# The unique *Legionella longbeachae* capsule favors intracellular replication and immune evasion

**DOI:** 10.1101/2023.11.22.568341

**Authors:** Silke Schmidt, Sonia Mondino, Laura Gomez-Valero, Pedro Escoll, Danielle P. A. Mascarenhas, Augusto Gonçalves, Christophe Rusniok, Martin Sachse, Maryse Moya- Nilges, Thierry Fontaine, Dario S. Zamboni, Carmen Buchrieser

**Affiliations:** Institut Pasteur, Université Paris Cité, Biologie des Bactéries Intracellulaires, CNRS UMR 6047, Paris, France; Sorbonne Université, Collège Doctoral, Paris, France; Department of Cell Biology, Medical School of Ribeirão Preto, FMRP/USP, Ribeirão Preto, Brazil; UTechS UBI, Centre de Ressources et Recherches Technologiques, Institut Pasteur, Paris, France; Biologie et Pathogénicité fongiques, Institut Pasteur, Paris, France

**Author notes:** For correspondence: Carmen Buchrieser Institut Pasteur, Biologie des Bactéries Intracellulaires 28, rue du Dr. Roux, 75724 Paris Cedex 15, France. Current address: Laboratory of Molecular and Structural Microbiology, Institut Pasteur de Montevideo, Montevideo, Uruguay (Sonia Mondino).

## Abstract

*Legionella longbeachae* and *Legionella pneumophila* cause Legionnaires’ disease despite species-specific differences in environmental niches, disease epidemiology, and genomic content. Here, we characterized a new *L. longbeachae* virulence factor, a capsule that is expressed in post-exponential growth phase as shown by electron microscopy. Analysis of the capsule composition *via* HLPC revealed the presence of a highly anionic polysaccharide absent in a capsule mutant. The capsule is crucial for replication and virulence *in vivo* in a mouse model of infection and in the natural host *Acanthamoeba castellanii*. It has anti-phagocytic function when encountering innate immune cells, it is involved in a low cytokine responses in mice and in human monocyte derived macrophages and helps to dampen the innate immune response. The *L. longbeachae* capsule is a novel virulence factor, unique among known *Legionella* species, that aids *L. longbeachae* to survive in specific niches and partly confers *L. longbeachae* its unique infection characteristics.

## Introduction

*Legionella longbeachae* is a rod-shaped Gram-negative bacterium that can cause Legionnaires’ disease, a severe form of pneumonia. Like other *Legionella* species, *L. longbeachae* is a facultative intracellular bacterium that causes the disease by inhalation of contaminated aerosols ^1^. *Legionella* spp. are typically found in aquatic environments, either as free-living bacteria or in biofilm communities ^2,3^. *Legionella longbeachae*, however, is predominantly isolated from potting soils and infections have been associated with gardening activities ^4,5^. The clinical symptoms of Legionnaires’ disease caused by *L. longbeachae* are similar to those caused by *L. pneumophila*, the most common causative agent of Legionnaires’ disease ^6^. Like *L. pneumophila, L. longbeachae* critically depends on a highly conserved type IVB Dot/Icm secretion system (T4SS) to establish an intracellular infection and to build a replication niche, the so-called *Legionella*-containing vacuole (LCV) ^7–10^. Analyses of the genomic content of the *Legionella* genus genome showed that it is highly diverse with an astounding 18,000 predicted T4SS effector proteins ^11^. For *L. longbeachae,* about 220 effectors were predicted, but only about 34% are shared with the *L. pneumophila* effector repertoire ^12,13^.

Interestingly, common laboratory mouse strains are resistant to *L. pneumophila* replication, except A/J mice which allow replication of *L. pneumophila* due to Naip5 mutations. In contrast, *L. longbeachae* effectively replicates in the lungs, disseminates, and causes death in A/J, BALBc, and C57BL/6 mice ^14,15^. It was hypothesized that this is due to the lack of flagella in *L. longbeachae*, as the detection of flagella expressed by *L. pneumophila* leads to a rapid activation of the NLRC4 inflammasome and clearing of the infection in BALBc or C57BL/6 mice ^16–18^. However, a flagellum-deficient *L. pneumophila* strain Paris is not lethal for C57BL/6 mice and it induces a robust pro-inflammatory cytokine response *in vitro* ^17,19^. In contrast, mice infected with wild type *L. longbeachae* die within 6 days after infection. *L. longbeachae* spreads from the primary site of infection (the lungs) to the blood and spleens of the animals. In contrast to *L. pneumophila, L. longbeachae* only induces a low pro-inflammatory cytokine response *in vitro* ^19^. A unique feature identified in the *L. longbeachae* genome is the presence of a gene cluster of 48 kb predicted to code for a capsule ^13^. It comprises 33 genes that are annotated as glycosyltransferases, enzymes for the synthesis of nucleotide sugar precursors, and an ABC transporter, likely for the export of the capsule. The transporter was found to be homologous to the capsule transporter of *Neisseria meningitidis* ^13^. We thus hypothesized that the capsule encoded in the *L. longbeachae* genome may be responsible for the enhanced virulence of *L. longbeachae* in mice as compared to *L. pneumophila* ^19^.

Capsules can protect bacteria from adverse environmental conditions, or antimicrobial agents such as antibiotics or antimicrobial peptides ^20–24^. Likewise, in pathogenic bacteria, capsules can mimic surface polysaccharides of mammalian cells, thus avoiding recognition by the host immune system ^25^. For example, the capsule of *E. coli* K1 contains polysialic acids that are anti-phagocytic and protect the bacteria from complement-mediated killing ^26,27^.

In this study, we visualized and characterized the *L. longbeachae* capsule and analyzed its functional role in infection. Comparing a knockout mutant in the capsule transporter and a wild type (WT) strain, we reveal that the capsule plays a key role during infection in mammalian and protozoan hosts. Furthermore, we provide evidence that the capsule is responsible for the low cytokine response in infected macrophages and mice. Thus, we demonstrate that the capsule is a novel virulence feature of *L. longbeachae*, which is unique among the known *Legionella* species.

## Results

### The L. longbeachae capsule cluster is unique among Legionella species

We first analyzed the presence of the putative capsule cluster and orthologous genes among 58 different *Legionella* species (Figure 1). This analysis shows that *L. longbeachae* is the only *Legionella* species to encode the entire 48 kb capsule cluster. Two other species, *L. massiliensis* and *L. gormanii,* carry genes similar to the ABC-type transporter genes *ctrBCD* and *bexD*. However, these two species lack most of the glycosyltransferases encoded by *L. longbeachae.* We did not find similar genes in the genomes of *L. pneumophila,* except for an orthologous gene of *llo3163,* a hydroxyacid dehydrogenase similar to *serA*, indicating that the capsule cluster is specific for *L. longbeachae* (see Supplementary Table 1).

**Figure 1:**
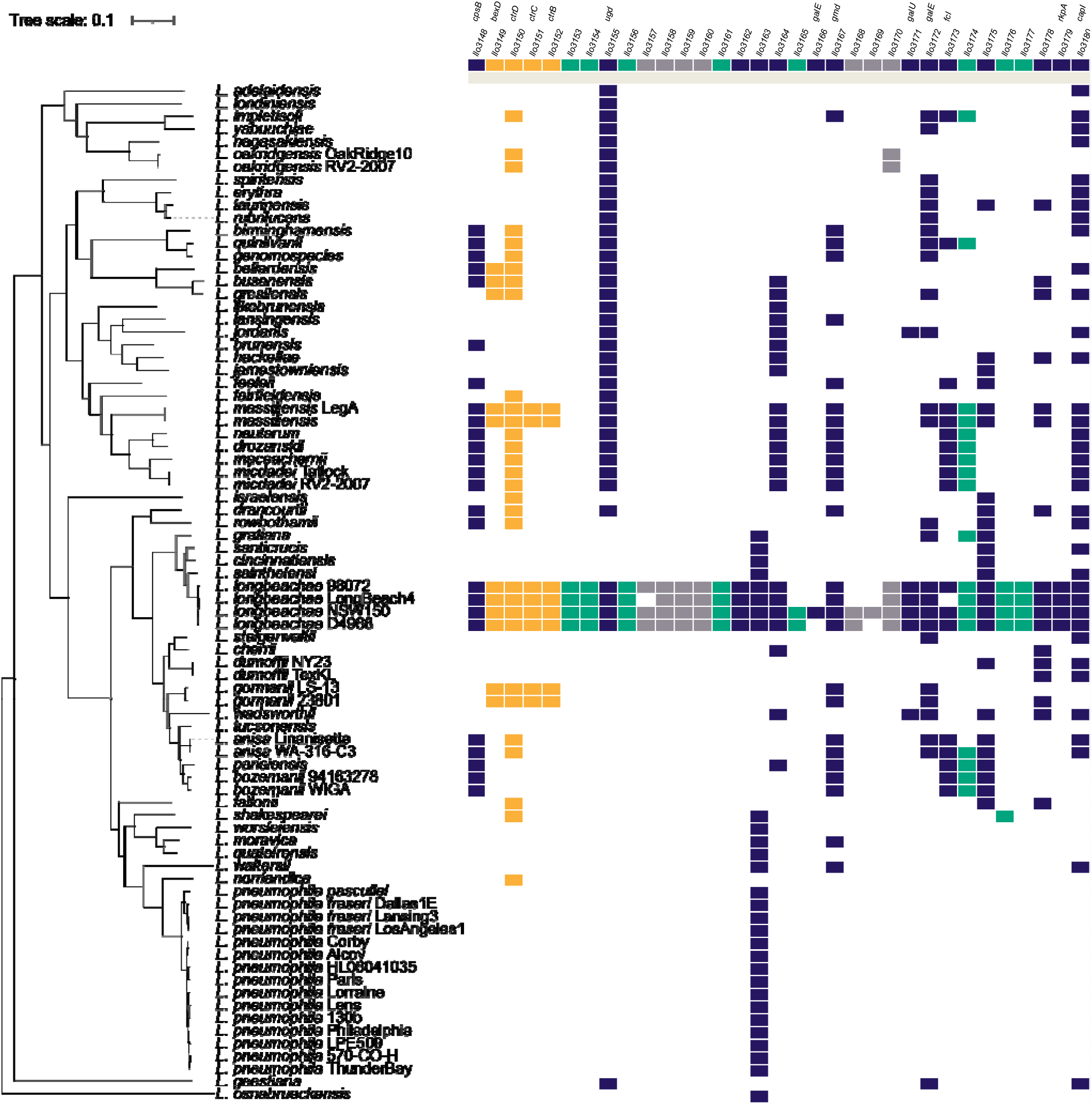
Presence or absence of orthologous genes coding for the capsule in 80 *Legionella* species/strains. Colored squares represent orthologous genes in comparison to the *L. longbeachae* NSW150 genome (depicted at the top). Orthologous gene reconstruction was done with PanOCT: amino acid percentage identity cutoff was 30%, BLAST e-value cutoff 10^−5^, and minimum percentage match length of subject and query was 65%.

To deepen these analyses, we selected twelve publicly available *L. longbeachae* strains, comprising nine strains from serogroup 1 (sg1) and three strains from serogroup 2 (sg2). All strains contain a similar capsule cluster whose genomic organization and genomic location are conserved (Figure S1). Only a small region in the central part of the capsule cluster shows few differences among the glycosyltransferases and a duplication of *galE* in the NSW150 and B41211CHC genomes (Figure 1 and Figure S1). All twelve strains encode genes for the predicted ABC type transporter *ctrBCD* and *bexD* (*ctrA*-like). Furthermore, they all encode glycosyltransferases and nucleotide sugar precursor genes belonging to the same groups of enzymes (Supplementary Table 2). This extremely high conservation in gene content, sequence similarity, and in the genomic location indicate that the acquisition of the capsule cluster dated back to a common ancestor of *L. longbeachae*.

### The L. longbeachae capsule cluster genes share homology to soil-dwelling bacteria

The evolutionary history of the capsule cluster in *L. longbeachae* is not known. Thus, we performed BLAST analysis on all its genes using *L. longbeachae* strain NSW150 (sg1) as the reference. Supplementary Table 1 lists the best hits obtained by BLAST search. Across the cluster, we find homologous genes from β-proteobacteria often found in soils, and of γ- and δ- proteobacteria such as the soil-dwelling *Burkholderia* spp., *Geobacter* spp., or *Pseudomonas* spp., but also from *Nitrococcus mobilis* or *Alteromonas* spp., which are present in marine environments. These similarities suggest that the capsule cluster was acquired from soil-dwelling bacteria.

### Electron microcopy analyses confirm that L. longbeachae expresses a capsule that is absent from L. pneumophila

To further analyze the *L. longbeachae* capsule and to study its functional role, we constructed a knockout mutant in the capsule transporter gene *ctrC* (*llo3151*). We chose *ctrC* as a target, since a knockout mutant of a homologous gene in *Campylobacter jejuni* abrogated capsule expression ^28^. When grown in ACES-buffered yeast extract broth (BYE) medium with or without apramycin, the Δ*ctrC* mutant did not display any significant growth differences as compared to the *L. longbeachae* WT (Figure S2). To confirm that *L. longbeachae* indeed expresses a capsule and that the identified gene cluster is responsible for its expression, we used transmission electron microscopy (TEM). The *L. longbeachae* WT and the Δ*ctrC* mutant as well as *L. pneumophila* WT (negative control) were grown in BYE until the optical density OD_600_ reached mid-log phase and late-log phase, referred to as exponential (E) phase and post-exponential (PE) phase, respectively. Cells were fixed and stained with cationized ferritin. TEM revealed a capsular layer surrounding the *L. longbeachae* WT (Figure 2A). The capsule was observed only in PE phase, indicating a growth phase dependent expression (Figure S3A). Its thickness was estimated to be between 50-100 nm, which is similar to ferritin-stained capsules observed in *E. coli* K30 ^29^. In contrast, the Δ*ctrC* mutant strain was devoid of such a layer (Figure 2A) and *L. pneumophila* WT cells were not stained by the ferritin dye (Figure 2A). Complementation of the Δ*ctrC* mutant strain with a plasmid containing *ctrC* as well as the downstream genes *ctrD* and *bexD* under the control of their native promoter restored capsule expression (Figure 2B). We included the downstream genes as macrocolonies of a Δ*ctrC* mutant strain complemented with *ctrC* and *ctrD* only barely restored the WT phenotype compared to those complemented with the longer construct (Figure S3B). However, only 50% of the imaged cells expressed a capsular structure indicating that regulation of capsule expression may be more complex (Figure S3C). Taken together, our results confirm that, in contrast to *L. pneumophila*, *L. longbeachae* expresses a capsule on its surface and that *ctrC* is essential for its expression on the cell surface.

**Figure 2:**
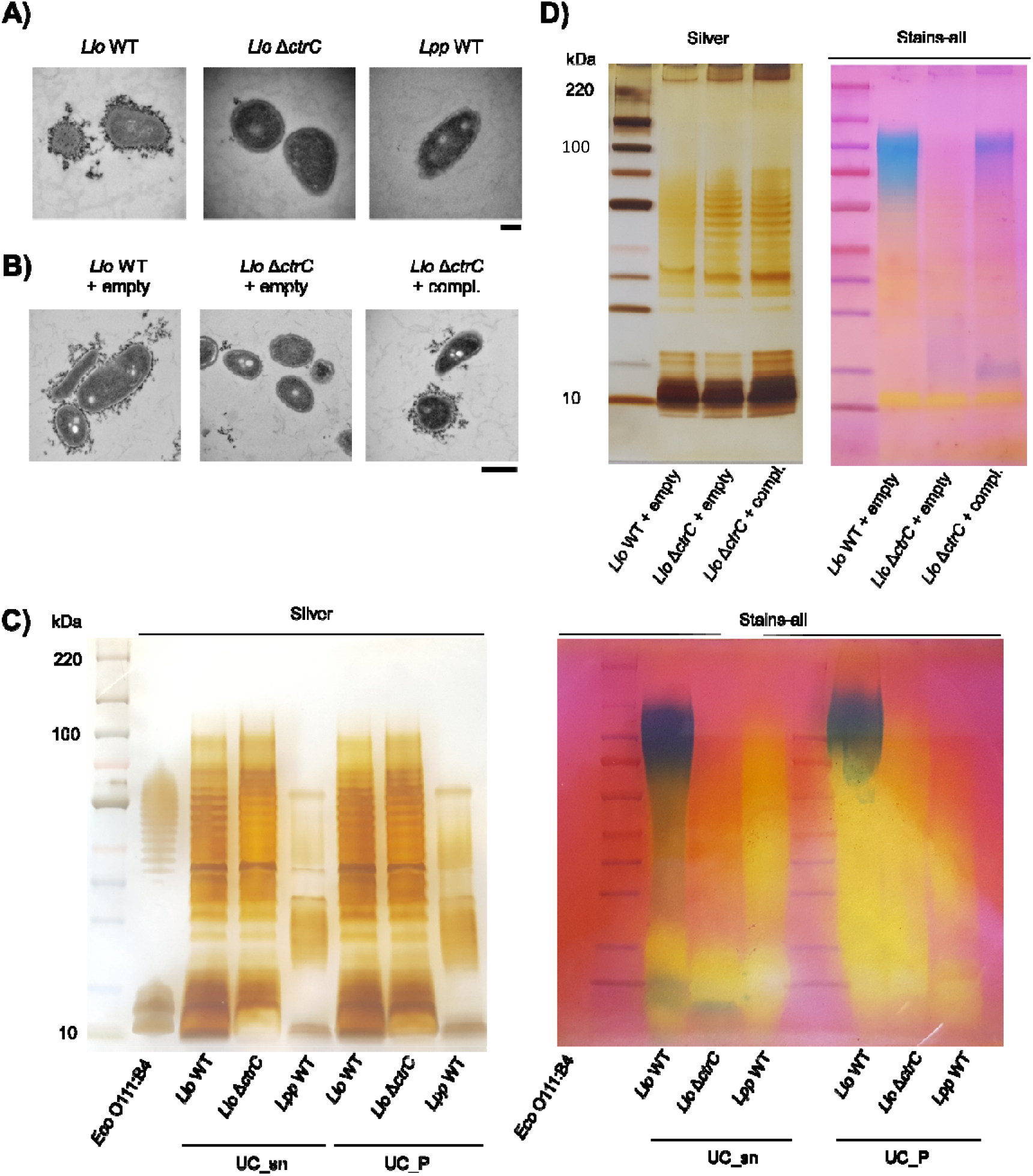
The *L. longbeachae* capsule is expressed in exponential growth phase and contains a highly anionic polysaccharide that cannot be stained by silver staining. A) *L. longbeachae* WT, Δ*ctrC* and *L. pneumophila* WT (negative control) were grown in BYE medium to post-exponential phase. Fixed cells were stained with cationized ferritin. TEM images are representative of n=3 independent experiments. Scale bar = 200 nm. **B)** *L. longbeachae* WT or Δ*ctrC* harboring an empty plasmid (pBCKS) or the complementation plasmid (SSM083) were fixed in post-exponential phase and processed like samples in A). Representative imgaes of n=3 independent experiments. Scale bar = 500 nm. **C)** Polysaccharides were extracted from *Llo* WT, Δ*ctrC* or *Lpp* WT by enzymatic isolation and subjected to ultracentrifugation. SDS gels were stained with silver nitrate or Stains-all dye. Purified LPS from *E. coli* O111:B4 (Sigma) was used as a control for silver staining. **D)** Extracts from *Llo* WT or Δ*ctrC* harboring the empty plasmid (pBCKS) or the complemented mutant were stained by silver nitrate or Stains-all.

### The highly anionic L. longbeachae capsule is resistant to silver staining

To isolate the *L. longbeachae* capsule and to characterize its composition, we first used phenol-extraction for polysaccharide isolation followed by HPLC analyses. This approach allowed the identification of galactosamine, glucosamine, mannose and possibly quinovosamine in both the WT and the capsule mutant extracts (Figure S4A). After elution of the column, we detected high peaks in the WT sample that might correspond to phosphosugars. However, when visualizing these samples by silver or Alcian blue staining, we observed that phenol extraction seems to collapse the LPS structure, similarly to what has been seen for *L. pneumophila* ^30^. Thus, *L. longbeachae* LPS nor CPS can be extracted by phenol (Figure S4B). An enzymatic isolation method ^31^ and silver staining revealed that *L. longbeachae* expresses a long O-antigen chain between 30-70 kDa. In contrast, *L. pneumophila* expresses two shorter O-antigen fractions as shown previously, confirming that enzymatic isolation preserves the LPS structure in both *Legionella* species (Figure 2C) ^32^. However, no differences between the *L. longbeachae* WT and capsule mutant were observed. Therefore, we tested other dyes to visualize CPS. When using Stains-all dye, a carbocyanine dye that stains highly anionic compounds such as glucosaminoglycans ^33^, the enzymatically prepared *L. longbeachae* extracts showed a strong signal at 100 kDa in the WT and the complemented strain, but neither the capsule mutant nor *L. pneumophila* (Figure 2C). When we complemented the capsule mutant, Stains-all reveals that the band observed in WT extracts is restored in the complemented mutant as well (Figure 2D). Again, silver staining did not reveal any CPS in the complemented strain (Figure 2D). The presence of a highly anionic PS corroborates our findings from staining with cationized ferritin for TEM and may explain our unique peak seen in HPLC. Taken together, the *L. longbeachae* capsule consists of highly anionic PS that cannot be stained by conventional silver staining methods.

### The capsule is expressed in vitro and during infection in a growth phase dependent manner

1. *L. pneumophila* is known to have distinct gene transcription profiles *in vitro* and during infection depending on the growth phase ^34^. Thus, to learn when the genes encoding the capsule of *L. longbeachae* are expressed and if their expression is growth phase dependent, we performed RNAseq analyses. The *L. longbeachae* WT and the Δ*ctrC* mutant strain were grown in BYE until E or PE phase, and we sequenced total RNA to follow their transcriptional profile in both growth phases. Almost all genes of the capsule cluster in the *L. longbeachae* WT strain were significantly downregulated in PE as compared to E phase (Figure 3A). This correlates with the visualization of the capsule in the PE phase (Figure S3A). Thus, expression of the capsule on the cell surface is growth phase dependent. Moreover, when comparing *L. longbeachae* WT and the Δ*ctrC* mutant strain, apart from the expression of *ctrC* and the downstream genes of the capsule transporter operon, no significant differences in the transcription of the other capsule genes were observed neither in E phase (Figure S5A) nor in PE phase (Figure S5B, Supplementary Table 3). Transcription of the capsule in E phase and its expression in PE phase suggest that *L. longbeachae* synthesizes its capsule later in infection, probably to be prepared for cell attachment and a new infection of host cells, and/or for survival in the environment.

To test this hypothesis, we analyzed the capsule expression during infection, using a dual reporter to follow the transcription of the capsule transporter operon *in cellulo*. This dual reporter contains a constitutively expressed red fluorescent protein (mKate2) and two copies of superfolder GFP (sfGFP) tagged with a degradation tag (sfGFP_deg_), each under the control of the native promoter of the capsule transporter. As a negative control we used a plasmid containing constitutively expressed mKate2 and two copies of sfGFP_deg_ without the native capsule promoter. These plasmids were transformed into *L. longbeachae* WT, which was used to infect THP-1 cells or *Acanthamoeba castellanii* (Figure 3B, Figure S6). We followed the expression of sfGFP and mKate2 over time by live confocal imaging. GFP was detected at 14 hours after infection in the promoter containing constructs but not the negative control. Our results show that the capsule is expressed in both eukaryotic hosts. Thus, the capsule is universally transcribed *in vitro* and upon infection.

**Figure 3:**
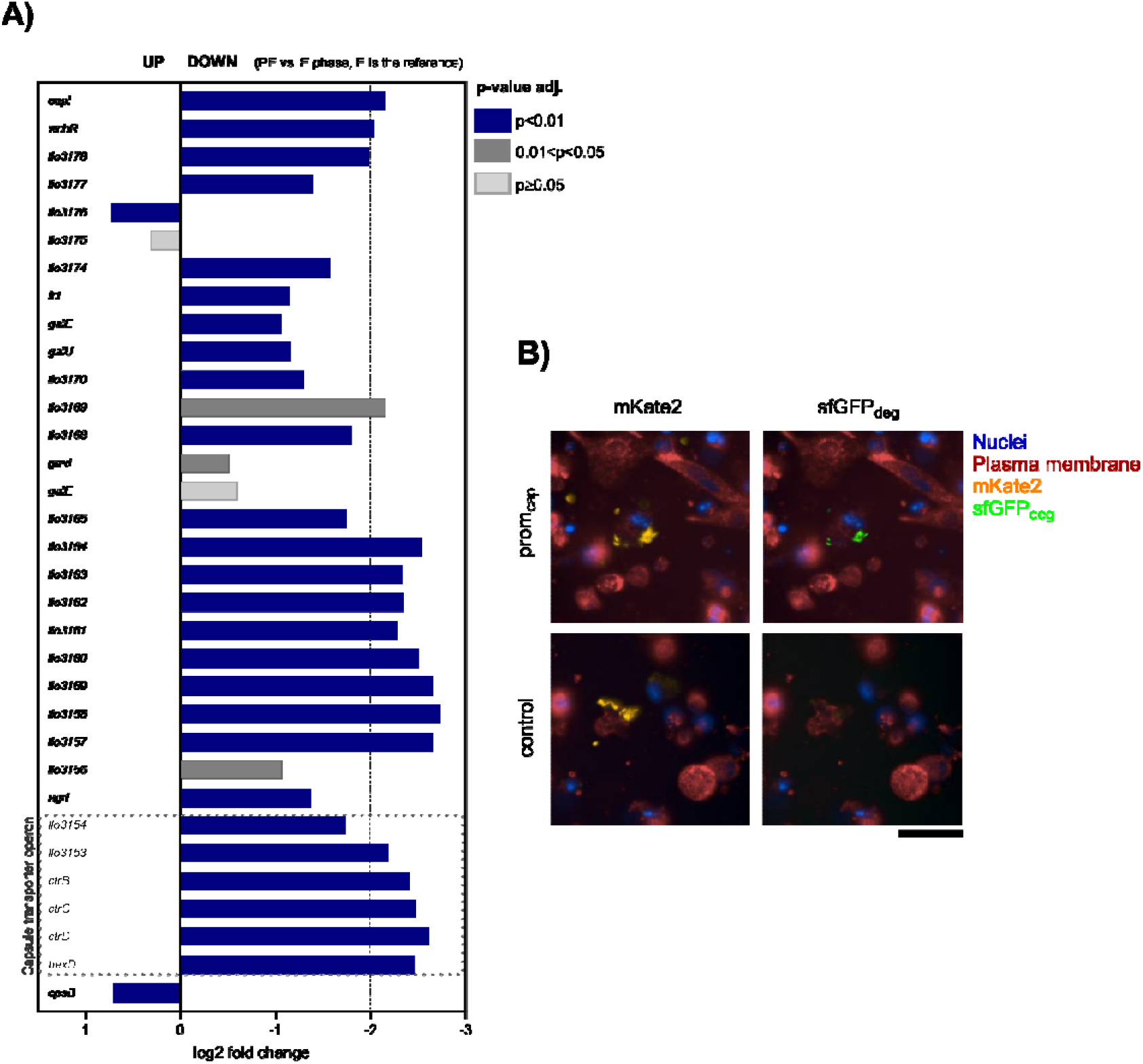
The *L. longbeachae* capsule is transcribed in liquid medium and *in cellulo*, but it is not expressed in a Δ*ctrC* mutant. A) RNAseq of WT transcripts (n=4); E vs. PE (E is the reference); consider relevant genes with log2 fold change of ±2 and adjusted p value ≤ 0.05. **B)** THP-1 cells were infected with wild type *L. longbeachae* expressing constitutive mKate2 and inducible sfGFPdeg under the control of the capsule promoter (prom_cap_). A control plasmid containing constitutive mKate2 and two copies of sfGFPdeg without the capsule promoter was included. Cells were imaged at 22 hours post infection. Scale bar = 50 µm.

### The capsule is crucial for virulence in vivo in mice and in the environmental host Acanthamoeba castellanii

To learn if the *L. longbeachae* capsule plays a role in virulence *in vivo*, we infected mice with either the *L. longbeachae* WT or the Δ*ctrC* mutant and different bacterial loads and followed their survival over 10-15 days. Upon infection with the WT strain, all animals succumb to the infection within 5 to 6 days, similar to what has been reported previously ^19^. However, all mice infected with the Δ*ctrC* mutant survived the infection (Figure 4A, Figure S7A). *L. longbeachae* WT replicates to higher numbers in the murine lungs as compared to the capsule mutant, but both seem to reach the blood stream as we measured comparable levels of CFUs in the spleen (Figures S7B, C). To confirm that the observed phenotype is due to the missing capsule, we infected mice with the *L. longbeachae* Δ*ctrC* mutant either complemented or harboring an empty plasmid. All mice infected with the mutant strain carrying the empty plasmid survived the infection. In contrast, 40% of mice infected with the complemented strain died, indicating that the capsule partially restored virulence *in vivo* (Figure 4B). In contrast, when we infected human macrophage-like THP-1 cells or murine bone marrow-derived macrophages *in vitro*, we observed only a slight replication delay of the mutant as compared to the WT (Figures S8A, B), indicating that successful infection by *L. longbeachae* depends on the interaction of immune cells with the capsule *in vivo*. Taken together, the lack of the capsule renders the bacteria avirulent in mice.

**Figure 4:**
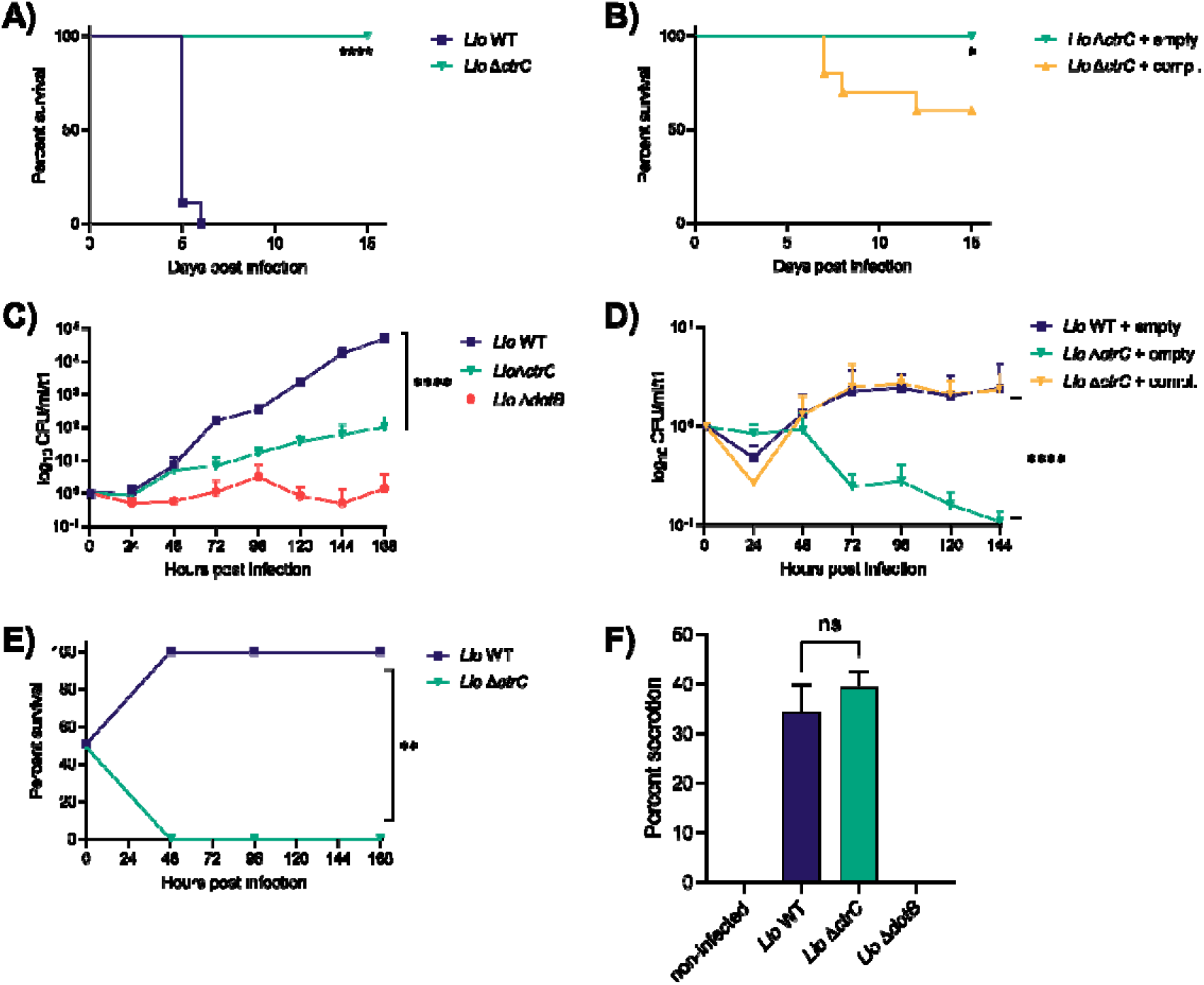
The capsule mutant is impaired in replication and avirulent *in vivo*, despite a functional Dot/Icm type IV secretion system. A) Female C57BL/6 mice were infected with 10^7^ CFU of *Llo* WT or Δ*ctrC* via the nasal route. Survival was monitored over 15 days. Each group contained 9 mice. **B)** Infection of female C57BL/6 mice with the capsule mutant harboring an empty plasmid (pBCKS) or the complementation plasmid (SSM083) with 10^7^ CFUs. 10 mice per group. Statistical significance was determined by Log-rank (Mantel-Cox) test. *, p≤0.05; ****, p≤0.0001. **C)** *Acanthamoeba castellanii* trophozoites were infected at MOI 0.1 at 20°C over 7 days. Samples were taken every 24 hours and CFUs are normalized to the input control. Data show means ± SD for n=3 independent experiments. **D)** *A. castellanii* trophozoites were infected with *Llo* WT or ΔctrC harboring an empty control plasmid (pBCKS) or the complementation plasmid (SSM083) at an MOI of 1. Samples were taken every 24 hours and CFUs normalized to t=1. Data show means ± SD for n=3 independent experiments. **E)** *A. castellanii* trophozoites were infected at MOI XYZ with equal amounts of *Llo* WT or Δ*ctrC*. Samples were taken every 24 hours and normalized to the input control. Data show % survival for n=2 independent experiments. Statistical significance was tested by non-linear regression analysis using a straight-line model. **F)** Translocation of the known T4SS effector RomA was tested in THP-1 cells infected with *Llo* WT, Δ*ctrC*, or a T4SS mutant, Δ*dotB*. Data represent means ± SD of n=2 experiments. Statistical significance was tested by two-tailed t-test. ns, non-significant; *, p≤0.05, **, p≤0.01, ***, p≤0.001, ****, p≤0.0001.

We then infected *A. castellanii* with *L. longbeachae* WT, the Δ*ctrC* or the Δ*dotB* mutant strain. The latter is deficient in the T4SS and was used as a negative control. We followed replication of the bacteria over seven days and observed a strong replication defect of the capsule mutant as compared to the WT (Figure 4C). As expected, the Δ*dotB* mutant failed to replicate in *A. castellanii* but seems to persist over time indicated by the stable CFU counts towards the end of the experiment. Like in mouse infections, *A. castellanii* infected with the complemented strain restored replication to WT levels while the capsule mutant harboring the empty plasmid was impaired in replication (Figure 4D). We further tested the competitive fitness of the capsule mutant in *A. castellanii* as described previously ^35^. This showed that the capsule mutant is completely outcompeted by the *L. longbeachae* WT strain within 48 hours of infection, further underlining the importance of the capsule in virulence of *L. longbeachae* in its environmental host (Figure 4E).

An important question that arose from the *in vivo* infections was whether the strong virulence defect of the capsule mutant may be due to impaired effector secretion through the Dot/Icm T4SS in the Δ*ctrC* strain. We thus tested effector translocation using the beta-lactamase (BlaM) secretion assay as described previously ^11^. We used a BlaM-fusion to the known T4SS effector RomA ^36^ and measured BlaM secretion in infected THP-1 cells by flow cytometry after 2 hours of infection. Both, the *L. longbeachae* WT and the Δ*ctrC* mutant translocated RomA successfully into the host cells to similar levels, whereas the Δ*dotB* mutant failed to translocate the effector (Figure 4F). Thus, the virulence phenotype of the capsule mutant *in vivo* and in *A. castellanii* is not due to an impaired Dot/Icm T4SS, as this strain maintains the ability to secrete effectors upon infection.

### Encapsulated L. longbeachae is more sensitive to salt stress

Capsules may protect bacteria from adverse environmental conditions, and it has been shown that salt stress in *L. pneumophila* is linked to virulence ^37^. Indeed, a *L. pneumophila* mutant in the response regulator LqsR was shown to be less virulent but more resistant to salt stress than WT bacteria ^37,38^. To test the role of the capsule under salt stress, we grew the *L. longbeachae* WT and capsule mutant as well as *L. pneumophila* WT to PE phase in liquid culture and plated the bacteria on BCYE in the presence or absence of 50 mM or 100 mM NaCl, respectively (Figure S9). At high salt concentrations, both the capsule mutant and *L. pneumophila* exhibit a 3-log-fold growth difference. Surprisingly, however, there is also a 2-log-fold growth difference between the *L. longbeachae* WT, and the capsule mutant grown in the presence of 100 mM NaCl. Hence, similar to *L. pneumophila*, the analysis reveals that the less virulent *L. longbeachae* capsule mutant is more resistant to salt stress (Figure S9). This growth difference was not detectable at a lower concentration of 50 mM NaCl, indicating that salt concentration is an important factor for optimal growth of both *Legionella* species.

### The L. longbeachae capsule delays phagocytosis in human cells

Based on data published for other bacteria, we hypothesized that the capsule of *L. longbeachae* could be implicated in modulating phagocytosis or attachment to eukaryotic cells. To quantify phagocytosis, we incubated the bacteria with THP-1 cells and added gentamicin to kill extracellular bacteria to follow infection by plating bacteria at different timepoints. For attachment studies, we pre-treated half of the cells with cytochalasin D (cytoD), a known inhibitor of actin polymerization, and followed the infection by plating the bacteria recovered from the cytoD-as well as the non-treated cells. We observed that WT bacteria were phagocytosed more slowly over time than the capsule mutant (Figure 5A). In contrast, both strains attached to THP-1 cells at similar rates, likely due to the synchronization of the infection by centrifugation at the start of each experiment (Figure 5B). This suggests that the capsule is important to delay phagocytosis by host cells.

**Figure 5:**
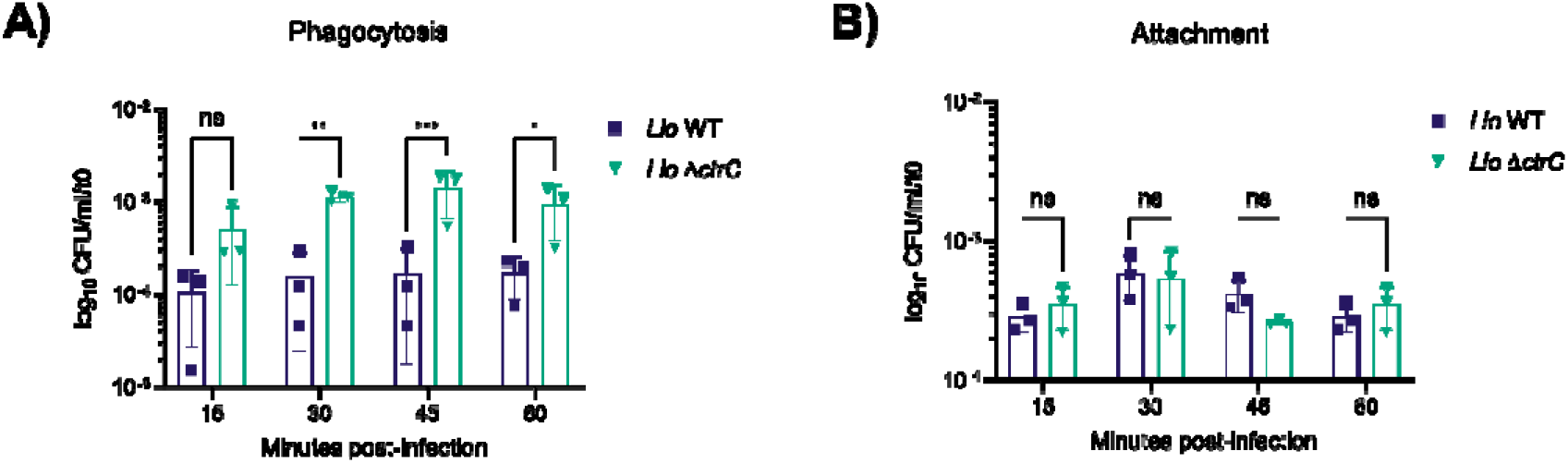
The capsule delays phagocytosis of host cells. A) For phagocytosis, differentiated THP-1 cells were infected with *Llo* WT or the *ctrC* mutant at MOI 10. Gentamycin was added at the indicated timepoints for 1 hour to kill extracellular bacteria and cells were lysed in water for CFU plating. Data show means ± SD of n=3 independent experiments normalized to the input control at t=0. **B)** For attachment, THP-1 cells were pre-treated with 2 µM cytochalasin D for two hours before infection with *Llo* WT or *ctrC* at MOI 10. At the indicated timepoints, gentamycin was added to half of the wells to kill extracellular bacteria. Cells were lysed in water and CFUs of gentamycin-treated cells (intracellular bacteria) and non-treated cells (total bacteria) were plated. Intracellular bacteria were deducted from total CFUs, normalized to the input control at t=0 and data plotted as means ± SD of n=3 independent experiments. Statistical significance was tested by two-way ANOVA with Tukey’s post-test. ns, non-significant; *, p≤0.05, **, p≤0.01, ***, p≤0.001.

### The L. longbeachae capsule impacts the pro-inflammatory cytokine response in primary innate immune cells

Capsules have been shown to mask cell structures like LPS or outer membrane proteins in bacteria. Due to their inherently low immunological recognition, capsules can have detrimental effects in infection settings such as reduced bacterial clearance or failure to mount an effective cytokine response against the pathogen ^25^. To learn whether the *L. longbeachae* capsule plays a role in the cytokine response, we infected primary human monocyte-derived macrophages (hMDMs) with the *L. longbeachae* WT and the capsule mutant strains and measured pro-inflammatory cytokine levels in cell supernatants using a high sensitivity SP-X human cytokine array (Simoa). We observed a significant increase in IL-6 levels in the capsule mutant compared to the complemented strain (Figure 6A). Moreover, cytokine levels of the complemented strain were as low as those of the *L. longbeachae* WT strain. In contrast, we did not detect significant differences in TNFα or IL-1β levels (Figure 6B-C). These cytokines are known mediators of inflammation in the context of *L. pneumophila* infections ^39,40^, and IL-6 and TNFα are highly induced upon infection with the *L. longbeachae* Δ*dotB* mutant. However, TNFα and IL-1β secretion in human macrophages do not seem to be driven by the capsule but are rather dependent on the presence of a functional Dot/Icm T4SS.

**Figure 6:**
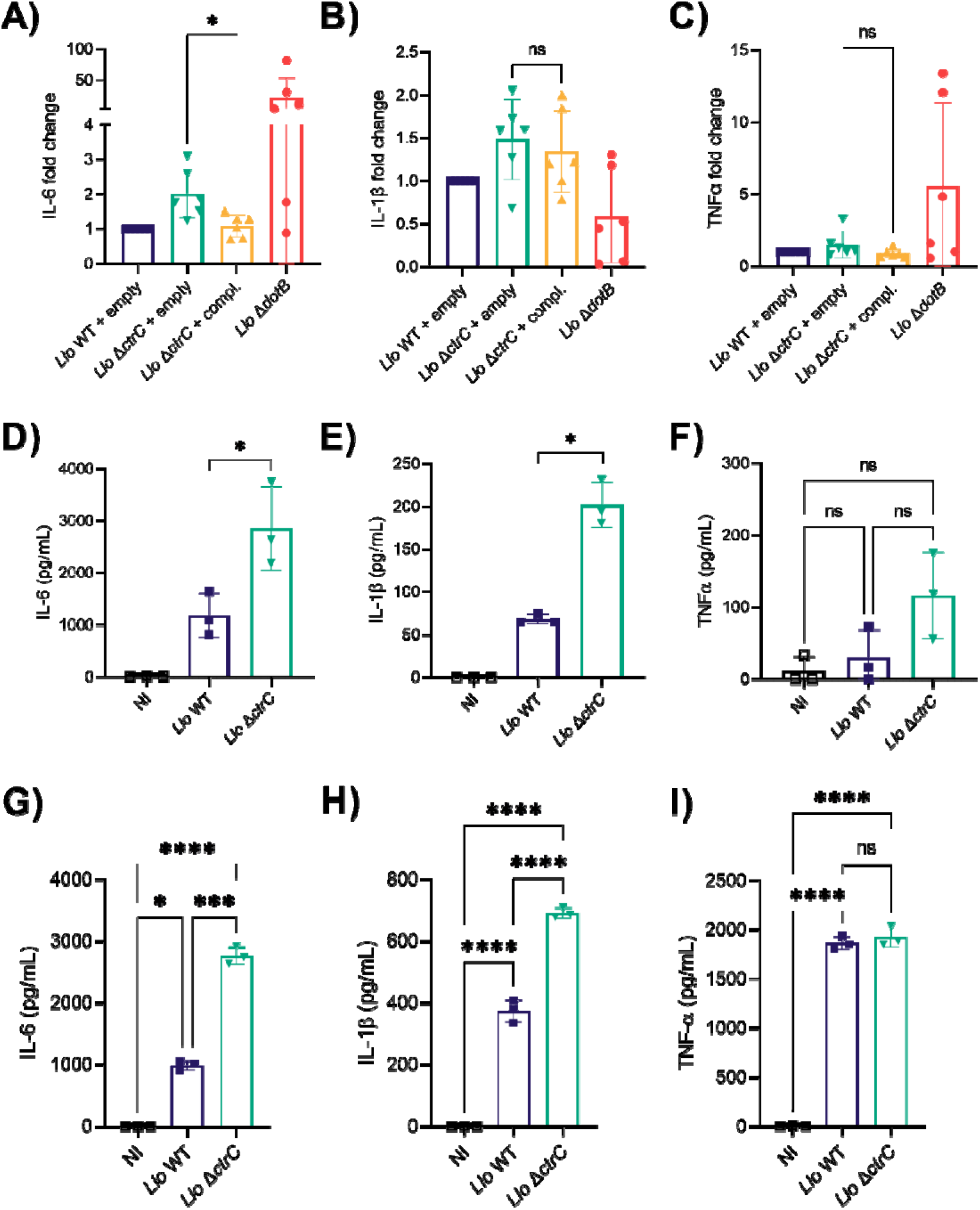
The capsule influences the cytokine response of primary macrophages and dendritic cells. A-D) Human monocyte derived macrophages (hMDMs) were infected at MOI 10 and cell supernatants were collected at 24 hours post-infection. High-sensitivity SP-X array was performed on cell supernatants. Data show means ± SD of fold change over *L. longbeachae* WT of n=6 independent experiments. Statistical significance was tested by pairwise t-tests; ns, non-significant; *, p≤0.05. **D-F)** Murine bone-marrow derived DCs were infected at MOI 10 and cell supernatants collected at 24 hours post-infection. Data show means ± SD of n=3 independent experiments. **G-I)** Murine bone-marrow derived macrophages were infected at MOI 10 and cell supernatants collected at 24 hours post-infection. Data show means ± SD of n=3 independent experiments. Statistical significance was tested by one-way ANOVA with Tukey’s post-test. ns, non-significant; *, p≤0.05; **, p≤0.01; ***, p≤0.001; ****, p≤0.0001.

*L. longbeachae* is lethal for mice and fails to induce a strong cytokine response *in vitro*^19^. Our results show that the capsule plays a crucial role in virulence, and we therefore tested whether cytokine secretion during infection is also impacted by its presence or absence. We infected murine dendritic cells (DCs) with *L. longbeachae* WT or the capsule mutant. Similar to human macrophages, the secretion of IL-6 was highly induced in the capsule mutant and lower in the *L. longbeachae* WT (Figure 6D). Secretion of IL-1β was also higher in murine DCs infected with the capsule mutant as compared to the WT (Figure 6E). Using murine bone marrow derived macrophages (BMDMs), higher IL-6 and IL-1β levels were detected in cells infected with the capsule mutant compared to WT infected or the complemented mutant (Figure 6G-H). However, we did not detect a significant difference in TNFα secretion in murine DCs or macrophages (Figure 6F, I). Thus, our results show that in both human and murine infection models the capsule dampens the secretion of pro-inflammatory cytokines.

## Discussion

In this study, we report that *L. longbeachae* expresses a capsule that is unique within the genus *Legionella* (Figure 1, Figure S1). Its genes seem to have been acquired from bacteria such as *Pseudomonas, Klebsiella,* or *Neisseria* which are present in soil environments or the soil-dwelling bacteria *Burkholderia* spp. and *Geobacter* spp., as evidenced by our comparative genomics analyses. This fits well with the environmental niche of *L. longbeachae*, as this bacterium is mostly isolated from moist soils and potting mixes ^4,41,42^. Importantly, the genetic organization of the cluster and its position in the genome are conserved among all *L. longbeachae* strains analyzed, suggesting that these genes have been acquired by horizontal gene transfer by a common ancestor of *L. longbeachae* (Figure S1). However, the exact organism from which it has been acquired, probably *en bloc* given the high conservation in all strains, is not known yet.

TEM analysis confirmed that *L. longbeachae* expresses a capsule, which can be stained by cationized ferritin, and that the *ctrC* mutant does not export the capsule (Figure 2A). CtrC is the inner membrane protein component in contact with CtrD, the nucleotide-binding domain containing ATPase of ABC capsule transporters. A deletion of the homologous *kpsM* gene in *C. jejuni* also abrogated capsule expression and impaired bacterial virulence in a ferret model of infection ^28,43^. Interestingly, RNAseq analyses showed that the capsule cluster is highly transcribed in E phase, and TEM analyses revealed that the *L. longbeachae* capsule is visible in PE phase (Figure 2B, S3). Thus, export of the capsule on the surface happens in later stages of bacterial growth, suggesting that the bacteria prepare themselves for a new infection and/or for survival in the environment. Indeed, using a fluorescence dual reporter to follow capsule transcription in living cells, we showed that the capsule transporter is expressed in both the human THP-1 cell line and in the environmental host *A. castellanii* late in infection.

Using conventional silver staining, we did not detect any fraction that corresponded to a possible CPS in the *L. longbeachae* WT, but we showed a similar LPS pattern in both the *L. longbeachae* WT and the Δ*ctrC* mutant (Figure S4). Hence, depletion of the capsule does not impact LPS expression, which might have been an explanation for the low virulence in the latter. Only when we applied Stains-all dye, we detected a band at around 100 kDa in the WT, which is absent in the mutant and in *L. pneumophila* (Figure 2C). Stains-all can be used for differential staining of highly anionic compounds such as glycosaminoglycans like hyaluronic acid, chondroitin sulfate, dermatan sulfate, or heparin ^33^. We possibly cannot detect the CPS by silver staining because it contains highly modified sugars, which cannot be oxidized by periodate during silver staining. Indeed, it has been reported that some *C. jejuni* strains express hyaluronic acid-like and teichoic acid like capsules, which cannot be stained by silver, due to a high degree of O-methyl phosphoramidate groups in these CPSs ^31,44,45^. Our results open new paths for further research into the biochemical composition of the *L. longbeachae* capsule to determine what confers the highly anionic charge and its anti-phagocytic function.

Capsules have also been shown to mediate resistance to environmental stresses such as osmotic stress ^46^. We tested hyperosmotic stress due to NaCl and showed that growth of the WT was highly impaired in the presence of 100 mM NaCl as compared to the capsule mutant and *L. pneumophila,* whose growth was less disturbed (Figure S9). This is similar to what has been shown for the less virulent LsqR mutant of *L. pneumophila,* and our data point to a link between virulence of *L. longbeachae* and osmotic stress ^38^. Capsules are hydrated shells surrounding bacteria, which may contain over 95% water ^47^ and which can protect them from hyperosmotic stress ^46,48,49^. According to our TEM studies, the capsule is likely negatively charged. Thus, it is plausible that Na^+^ ions present in high salt environments may disrupt the integrity of the capsule, causing membrane stress and leading to a growth defect in the WT.

Capsules have been shown to be important virulence factors of many Gram-negative bacteria that are human pathogens, such as *N. meningitidis* group B*, E. coli* K1 and K5*, K. pneumoniae*, or *Haemophilus influenzae* type b ^50–55^. Using a mouse model of infection revealed that the capsule is a crucial virulence factor of *L. longbeachae* (Figure 4A). Remarkably, the capsule mutant was avirulent in mice, a phenotype that was partially restored *in vivo* by complementation (Figure 4B). The virulence of *L. longbeachae* is linked to its capacity to replicate better in the lungs of infected mice (Figure S7B). Previous studies have shown that *L. longbeachae* infection is lethal in common laboratory mouse strains ^14,15,19^. Here, we provide evidence that the observed high virulence for mice is due to the capsule encoded by *L. longbeachae*. Thus, the *L. longbeachae* capsule is the first virulence factor described for human pathogenetic *Legionella* whose virulence phenotype *in vivo* is comparable with that of the loss of a functional Dot/Icm type IV secretion system.

Early studies on the pathogenicity of *L. pneumophila* revealed that the bacteria can replicate inside the amoeba and likely use the same mechanisms for infection of human cells^56^. Here, we show that the capsule mutant is impaired in intracellular replication in *A. castellanii*, and upon coinfection the WT rapidly outcompetes the capsule mutant (Figure 4C, E). Thus, the capsule of *L. longbeachae* is important for infection of both mammalian cells and environmental protozoa. Importantly, all *Legionella* species analyzed to date critically depend on the secretion of effector proteins into host cells *via* a highly conserved Dot/Icm T4SS, to establish infection and their replicative vacuole ^7,10,57–60^. A *L. longbeachae* Δ*dotB* mutant having a nonfunctional T4SS cannot replicate in THP-1 cells, A549 cells and HEK cells *in vitro* ^60^. Here, we show that it also fails to replicate in *A. castellanii* (Figure 4C). However, the capsule mutant is not impaired in effector translocation through the T4SS (Figure 4F). Thus, the observed infection phenotypes are not linked to a dysfunctional T4SS but to the lack of a capsule in the mutant strain.

Capsules have been shown to modulate phagocytosis by host cells by masking cellular receptors on the bacterial surface, or by impairing recognition of LPS by TLRs ^61–63^. Indeed, in *L. longbeachae*, the capsule is involved in delaying bacterial phagocytosis by THP-1 cells (Figure 5A). This shows that the *L. longbeachae* capsule is an anti-phagocytic factor and it shields bacterial outer membrane components from immune recognition. Further studies may shed light on whether the *L. longbeachae* capsule helps in evading the binding of complement factors, which may explain the difference in phagocytosis.

By avoiding immune recognition, bacterial capsules have been shown to dampen the host pro-inflammatory cytokine response ^52,61,64^. The *Salmonella enterica* serotype Typhi Vi capsule dampens the expression of pro-inflammatory cytokines IL-6, TNFα, and IL8, and it prevents recognition by TLR4 ^65,66^. A non-encapsulated strain of *K. pneumoniae* induces higher IL-6 levels in bronchoalveolar lavage fluid from infected mice than an encapsulated WT strain ^67^. Furthermore, it has been shown that the lack of capsule in *C. jejuni* leads to an increased release of IL-6 and TNFα in murine dendritic cells and macrophages ^68,69^. Similarly, the *L. longbeachae* capsule modulates cytokine release in hMDMs, inducing a lower response pro-inflammatory IL-6 in the WT as compared to the capsule mutant (Figure 6A). However, we did not detect any differences in secretion of TNFα or IL-1β in human cells. In contrast, when we infected murine dendritic cells and murine macrophages, the capsule mutant induced a higher cytokine response for IL-6 and IL-1β, but not TNFα (Figure 6D-I), suggesting that the absence of capsule permits the liberation of bacterial-derived immunostimulatory components that would be only recognized by murine cells and not by human cells. Such immunostimulatory bacterial-derived components and their receptor counterparts in the host are yet to be identified, however an existing example of such differences between murine and human macrophages is the NAIP5 inflammasome, present in mice and absent in humans, which allows the exclusive recognition of flagellin from *L. pneumophila* by murine macrophages ^70^. However, the leaked immunostimulatory component(s) cannot be flagellin as it is absent from *L. longbeachae*. Thus, the *L. longbeachae* capsule dampens the immune response in innate immune cells by a mechanism that might involve the masking of immunostimulatory bacterial-derived components and their release, avoiding their recognition by innate immune pattern-recognition receptors.

In conclusion, our study provides evidence for a novel virulence mechanism of *L. longbeachae,* the expression of a capsule. This capsule is important for *L. longbeachae* replication in the environmental host *A. castellanii* and in mice. During infection, the *L. longbeachae* capsule modulates phagocytosis and dampens the innate immune response favoring bacterial replication. Importantly, our findings provide exciting new insights into how *Legionella* escape recognition by host cells and their defenses, and it opens new avenues to explore the architecture of this unique capsular type and to discover its anti-phagocytic molecules.

## Methods

### Bacterial strains, growth conditions and cell culture

Bacterial strains used in this study are listed in Supplementary Table 4. *Escherichia coli* DH5α subcloning efficiency bacteria were grown in Luria-Bertani broth or on LB agar. *L. longbeachae* and *L. pneumophila* were cultured in N-(2-acetamido)-2-aminoethanesulfonic acid (ACES)-buffered yeast extract broth (BYE) or on ACES-buffered charcoal-yeast (BCYE) extract agar with antibiotics added where appropriate ^2^. To *L. longbeachae* Δ*ctrC*, 15 µg/ml apramycin (Sigma), for plasmid maintenance, 5 µg/ml of chloramphenicol were added. CDM or MDM was used as minimal medium according to published protocols ^3^. *Acanthamoeba castellanii* strain C3 (ATCC 50739) trophozoites were grown in PYG medium according to published protocols ^4,5^. As infection buffer PYG 712 medium [2% proteose peptone, 0.1% yeast extract, 0.1 M glucose, 4 mM MgSO_4_, 0.4 M CaCl_2_, 0.1% sodium citrate dihydrate, 0.05 mM Fe(NH_4_)_2_(SO_4_)_2_•6H_2_O, 2.5 mM NaH_2_PO_3_, 2.5 mM K_2_HPO_3_] without proteose peptone, yeast extract, and glucose was used. The human monocyte derived THP-1 cell line (ATCC TIB-202) was maintained in RPMI GlutaMax supplemented with 10% fetal calf serum (FCS) at 37°C and 5% CO_2_. For infections, undifferentiated THP-1 cells were seeded with RPMI and 50 µg/ml of Phorbol 12-myristate 13-acetate (PMA) to start differentiation into adherent cells. Cells were grown in the presence of PMA for three days and recovered overnight in fresh RPMI medium before infection.

### Construction of L. longbeachae Δ*ctrC*

The 0.5 kb flanking regions of *L. longbeachae ctrC* (*llo3151*) were amplified using the primer pairs P3/P4 (upstream segment) and P1/P2 (downstream segment) using genomic DNA of *L. longbeachae* NSW150 as a template (see primers listed in Supplementary Table 5). The apramycin cassette was amplified with primer pair Apra_s/Apra_as using pTOPO-ApraR plasmid as template. The three PCR fragments were ligated by PCR using primers P1/P3 and cloned into pGEM®-T easy vector (Promega). Plasmid was then digested with NotI, and the combined *ctrC*::Apramycin fragment was ligated into the suicide plasmid pLAW344 ^6^ cut with the same restriction enzyme. The resulting plasmid, pLGV012, was introduced into *L. longbeachae* NSW150 as described above, and transformants were plated onto BCYE agar with apramycin. Resulting colonies were grown in BYE with apramycin and plated onto BCYE agar supplemented with apramycin and 5% sucrose. Sucrose-resistant/chloramphenicol-sensitive clones were screened, and successful deletion was verified through PCR and whole genome sequencing.

### Transformation of *L. longbeachae*

1. *L. longbeachae* was grown on BCYE agar plates at 37°C. Bacteria from a fresh plate were washed three times with ice-cold 10% glycerol, and the final pellet was resuspended in ice-cold 10% glycerol. For transformation, a 400 µl aliquot of electrocompetent cells was freshly mixed with 300-600 ng of plasmid DNA and electroporated at 2.5 kV, 1000 Ω, and 25 µF. The cultures were recovered in BYE broth at 37°C with shaking for 16 h and then plated onto BCYE agar with the appropriate antibiotic.

### Transmission electron microscopy

Bacterial strains were grown overnight in BYE and OD_600_ was followed constantly to determine E phase (OD_600_ 2.0-2.5) or PE phase (OD_600_ 3.7-4.2). Samples of 10 ml were taken at E and PE phase and centrifuged for 15 minutes at 500 g. The medium was discarded, and cells were immediately fixed overnight in 0.1 M cacodylate fixation buffer containing 2.5% glutaraldehyde. Subsequently, fixed cells were washed in 0.2 M cacodylate buffer and stained with 1 mg/ml cationized ferritin (Sigma, F7879-2ML) for 30 minutes at room temperature. After another wash, cells were immobilized in 4% agar before osmium fixation. Agar blocks were sequentially dehydrated in increasing volumes of ethanol and embedded in resin and thin sections were sliced on a Leica UC7 ultramicrotome to 70 µm thickness. Resin slices were mounted and imaged on a Tecnai BioTWIN 20-120 kV transmission electron microscope.

### RNA sequencing and analysis

Bacteria were grown at 37°C in BYE and OD_600_ was measured to follow their growth. When bacteria reached exponential phase (OD_600_ 2.0-2.5), 2x5 ml were pelleted and snap frozen in dry ice and ethanol. Bacterial morphology was checked under a microscope. Similarly, pellets were collected from bacteria grown to post-exponential phase (OD_600_ 3.7-4.2) and stationary phase the following day. Bacterial pellets were resuspended in Qiazol (Qiagen) and RNA was isolated according to the miRNA Mini kit (Qiagen). The samples were Turbo DNase digested (Thermo Scientific) and rRNA was depleted using the RiboCop rRNA Depletion Kit for Gram-negative bacteria (Lexogen). Depleted RNA was metal-catalyzed heat-fragmented using the RNA Fragmentation kit (Ambion). RNA quantity was measured using a Qubit 2.0 (Invitrogen) and size distribution was confirmed between 100-200 nt by BioAnalyzer (Agilent Technologies). Fragmented RNA was subsequently processed according to the TruSeq mRNA sample preparation guide by Illumina. RNAseq was performed using Illumina NextSeq 550 multiplex sequencing (Illumina). For analyzing capsule gene expression, we used FASTQ files containing single-end reads generated by Illumina sequencing. Sequencing reads were processed with Cutadapt software (version 1.15) to remove adapters. Trimming was performed with Sickle (version 1.33, https://github.com/najoshi/sickle) with a quality threshold (Phred Score) of 20. Reads shorter than 20 nucleotides were discarded. Clean reads were aligned to the *Legionella longbeachae* NSW150 sequence using Bowtie2 (version 2.3.4.3), and only uniquely mapped reads were kept for the read counts. We used Samtools package (https://github.com/samtools/samtools) to build indexed BAM files from the mapping results. To count the number of reads overlapping each genomic feature, we used featureCounts from the Subread package (version 1.6.3). Only primary alignments were counted. Differential analysis between the different conditions was performed with the R package Sartools (version 1.3.0) using DESeq2 methods. The “median” option was used to compute size factors.

### Construction of complementation and dual reporter plasmids

Capsule cluster promoter was amplified using primers SSM_79 and SSM_80, and the region containing *ctrC* (*llo3151*) and *ctrD* (*llo3150*) was amplified using primers SSM_82 and SSM_85. Both PCRs were performed using genomic DNA of *L. longbeachae* NSW150 as a template. Fragments were then ligated by PCR, with primers SSM_86 and SSM_87. The obtained amplicon was cloned into pBCKS (Stratagene) by restriction-free cloning ^7^, resulting in plasmid SSM073. To construct plasmid SSM083, *bexD* (*llo3149*) was amplified using primers SSM_113 and SSM_114, and subcloned into SSM073 vector by restriction-free cloning ^7^. For construction of the reporter plasmid, the capsule transporter promoter (prom_cap_) was PCR amplified using primers SSO_025 and SSO_049 (prom_cap_1) or SSO_050 and SSO_049 (prom_cap_2) (listed in Table 2). Superfolder GFP (sfGFP, gift from David Bikard) was PCR amplified using primers SSO_051 and SSO_052 (sfGFP1) or SSO_053 and SSO_054 (sfGFP2). A C-terminal degradation tag was added to each sfGFP construct by restriction-free cloning ^7^ using primers SSO_045 and SSO_046 or SSO_047 and SSO_048. The GFP constructs were fused to the capsule transporter promoter by overlap PCR, one copy with restriction sites for BamHI and HindIII and the second copy with restriction sites HindIII and KpnI. Each copy was first ligated into pBCKS and inserts were confirmed by sequencing. The second copy was subsequently ligated into the first copy pBCKS vector using restriction sites HindIII and KpnI. One copy of mKate2 (Addgene #68441) was fused to a strong promoter from the Anderson collection (Table 2) by overlap PCR using primers SSO_065 and SSO_066 and ligated into the two copy pBCKS vector through restriction sites NotI and BamHI. A control plasmid was constructed by replacing the two promoter regions cap_prom_ with a triple stop codon by restriction-free cloning using primers SSO_621/SSO_622 and SSO_623/SSO_624, respectively. Plasmid DNA was isolated using Nucleospin Plasmid kit (Macherey Nagel) and all plasmids were confirmed by sequencing.

### Imaging of dual reporter in THP-1 cells and *A. castellanii*

For imaging of THP-1 cells, bacteria expressing the dual reporter constructs (pSS016 or pSS017) were grown to post-exponential phase in BYE medium with 5 µg/ml chloramphenicol. Differentiated THP-1 cells were infected at an MOI of 10 in a µClear 96-well plate (Greiner) for 1 hour. Cells were washed three times in PBS and supplemented with fresh RPMI medium. Subsequently, cells were stained with CellMask™ Deep Red Plasma Membrane Stain (Invitrogen, C10046) at 5 µg/ml for 20 minutes and washed once. Nuclei were stained with Hoechst dye (Invitrogen, H3570) at 1 µg/ml. Live imaging was performed at 40x magnification using the Opera Phenix confocal microscope (PerkinElmer). Images were obtained every hour and analyzed using the Harmony® high-content analysis software (PerkinElmer). *A. castellanii* were infected in infection buffer at an MOI of 10 for 1 hour at 37°C. After one hour, cells were extensively washed in PBS to remove extracellular bacteria and maintained in infection buffer at 37°C. Cells were imaged at 40x magnification using the EVOS inverted digital microscope (Thermo Fisher).

### Animals and *in vivo* infections

Mice used in this study were bred and maintained in institutional animal facilities of University of São Paulo – School of Medicine of Ribeirão Preto/SP. All mice were at least 8 weeks old at the time of infection and were in a C57BL/6 (Jax 000664) genetic background. For the survival and CFU experiments, approximately 10 and 7 mice per group were used, as indicated in the figures. For *in vivo* experiments, the mice were anesthetized with ketamine and xylazine (300 mg/kg and 30 mg/kg, respectively) by intraperitoneal injection followed by intranasal inoculation with 40 μl of RPMI 1640 containing bacteria. For CFU determination, the lungs were harvested and homogenized in 5 ml of RPMI 1640 in a tissue homogenizer (Power Gen 125; Thermo Scientific) ^8,9^. Lung homogenates were diluted in RPMI and plated on BCYE agar plates containing streptomycin for CFU determination as previously described. For survival determination, mice were observed once a day with the measurement of their weight ^10^. The care of the mice followed the institutional guidelines on ethics in animal experiments.

### Bone marrow-derived dendritic cells and macrophages

Bone marrow-derived dendritic cells (BMDCs) and bone marrow-derived macrophages (BMDMs) were generated from C57BL/6 mice as previously described ^11,12^. Mice were euthanized and bone marrow cells were obtained from femurs and tibias. BMDCs were harvested from femurs and differentiated with RPMI 1640 (Gibco, Thermo Fisher) containing 20% Fetal Bovine Serum (FBS, Gibco) and 20 ng/ml of recombinant GM-CSF (eBioscience), 2 mM L-glutamine (Sigma-Aldrich), 15 mM Hepes (Gibco) and 100 U/ml penicillin-streptomycin (Sigma-Aldrich) at 37°C with 5% CO_2_ for 7 days. The non-adherent/loosely adherent fraction was harvested by collecting the culture supernatant and carefully washing the plate with PBS. After centrifugation of the total volume collected, BMDCs were resuspended in RPMI 1640 supplemented with 10% FBS and plated as indicated. BMDMs were harvested from femurs and differentiated with RPMI 1640 (Gibco, Thermo Fisher) containing 20% FBS and 30% L929-Cell Conditioned Medium (LCCM), 2 mM L-glutamine (Sigma-Aldrich), 15 mM Hepes (Gibco) and 100 U/ml penicillin-streptomycin (Sigma-Aldrich) at 37°C with 5% CO_2_ for 7 days. Of note, in some experiments LCCM was replaced for 10% of a conditional medium from 3T3 cells stably expressing mouse MCSF. Cells were detached with cold PBS, resuspended in RPMI 1640 supplemented with 10% FBS and plated as indicated for each method.

### Replication and competition assays in *Acanthamoeba castellanii*

Amoeba trophozoites grown at 20°C were washed in infection buffer, seeded at 1x10^6^/ml and left to adhere for one hour prior to infection. Bacteria were resuspended in infection buffer to an MOI of 0.1 and left to infect for 1 hour. Dilutions of an aliquot of input bacteria was plated on BCYE to determine CFUs used for infection (t0). After one hour of infection, amoeba were washed three times in PBS to remove extracellular bacteria and resuspended in infection buffer. Samples of 500 µl were taken at this timepoint (t1), centrifuged at 14000 rpm for 3 minutes and vortexed for one minute to break up amoebae. Samples were subsequently taken every 24 hours for seven days. Experiments were carried out in triplicates and CFUs were counted after three to four days of growth on BCYE at 37°C. Competition assay was carried out as previously described ^13^. Briefly, *A. castellanii* (5 × 10^6^ per flask) in infection buffer was infected at an MOI of 0.1 with a 1:1 mix of wild type *L. longbeachae* and Δ*ctrC* mutant bacteria. The infected amoebae were grown for seven days at 37°C. Every two days, a sample of lysed amoebae was diluted 1:100 and used to infect a fresh flask of amoebae (100 μl homogenate per flask). Dilutions were plated on BCYE agar plates containing apramycin or not, to determine CFUs.

### Replication assays in THP-1 cells

Infection assays of THP-1 cells (ATTC™: TIB-202) were done as previously described ^14^. Briefly, cells were seeded and differentiated into macrophage-like adherent cells in 12-well tissue culture trays (Falcon, BD lab ware) at a density of 2 x 10^5^ cells/well. Stationary phase *L. longbeachae* were resuspended in serum free medium and added to cells at an MOI of 10. After 2 h of incubation, cells were treated with 100 [g/ml gentamycin for 30 minutes to kill extracellular bacteria. Infected cells were then washed before incubation with serum-free medium. At 2, 24, 48, and 72 h, THP-1 cells were lysed with 0.1% Triton X-100. The infection efficiency of the different *L. longbeachae* strains was monitored by determining the number of CFUs after plating on BCYE agar.

### Bacterial replication in BMDMs

For CFU determination, macrophages were seeded at 2×10^5^ cells/well in 24-well plates and cultivated in RPMI 1640 with 10% FBS. Cultures were infected at a multiplicity of infection (MOI) of 10, centrifuged for 5 minutes at 200 ×g at room temperature. After 1 hour of infection, BMDMs were washed twice with PBS, and 1 ml of medium was added to each well. For CFU determination, the cultures were lysed in sterile water, and the cell lysates were combined with the cell culture supernatant from the respective wells. Lysates plus supernatants from each well were diluted in water, plated on BCYE agar plates, and incubated for 4 days at 37°C for CFU determination ^8,9^.

### Beta-lactamase translocation assay

Translocation of T4SS effectors was performed as previously described ^15^. Plasmids pXDC61 or SSM012 were electroporated into *L. longbeachae* strains or *L. pneumophila* strain Paris as a positive control. Triplicate wells of THP-1 cells were seeded in 96-well plates at 10^5^ cells/well and differentiated into adherent cells with 50 µg/ml PMA for three days. Cells were recovered in RPMI without PMA for another night. Freshly transformed bacteria were induced by addition of 1 mM IPTG and THP-1 cells were infected at MOI 50 with stationary phase bacteria. Cells were subsequently centrifuged to synchronize the infection. At 1h30 after infection cell were incubated with CCF4-AM (Life Technologies) and 0.1 M probenecid at room temperature in the dark. After another 1.5 hours, cells were detached with non-enzymatic cell dissociation solution (Sigma). Flow cytometry was performed using a MACSQuant VYB system (Miltenyi Biotec), with excitation at 405 nm (violet) and emission collection with filters at 525/50 nm (Green) and 450/50 nm (Blue). Flow data were analyzed by FlowJo 10 software (LLC). Gates for green and blue fluorescence were set based on uninfected cells without any treatment. Western blots were performed on protein lysates from induced bacteria to confirm expression of blaM.

### Attachment and phagocytosis assays

THP-1 cells (ATTC™: TIB-202) were seeded at 4x10^5^ cells/well in 12 well plates in RPMI-10% FCS and differentiated for three days with 50 nM PMA. After 96 hours of differentiation, RPMI without PMA was used to recover the cells overnight. For attachment assays, cells were treated with 2 µM cytochalasin D (Sigma) for two hours prior to infection. Bacterial strains were grown in BYE overnight and bacterial growth was monitored until the bacteria reached post-exponential growth phase (OD_600_ 3.7-4.2). The bacteria were diluted to reach an MOI of 10 in RPMI with or without 2 µM cytochalasin D for attachment assays. At the indicated timepoints, cells were washed three times with PBS to remove non-adhered bacteria and half of the wells were treated with 100 µg/ml gentamicin for one hour to kill extracellular bacteria (control for internalized bacteria). Cells were lysed by addition of ddH_2_O for 15’ at 37°C. Dilutions of CFUs were plated for both the gentamicin-treated and untreated wells to determine total CFUs vs. adhered CFUs, respectively. For phagocytosis assays, cells were washed at the indicated timepoints and treated with 100 µg/ml gentamicin for one hour to kill extracellular bacteria. Cells were lysed in ddH_2_O for 15 minutes at 37°C and dilutions were plated to determine CFUs. An aliquot of bacteria used for infections was plated for each strain to determine CFUs at t0.

### Osmotic stress assay

Bacteria were grown in BYE to PE phase (OD_600_ 3.7-4.2) and diluted to 2x10^9^/ml. Ten-fold dilutions were spotted in triplicates onto BCYE plates supplemented with 50 mM NaCl, or 100 mM NaCl, or plain BCYE plates and bacteria were grown at 37°C for 6 days.

### Polysaccharide extractions, gel electrophoresis and HPLC analysis

Bacteria were grown to PH phase in BYE medium and washed in PBS. For phenol extraction, cells were treated with 45% hot phenol for 30 minutes according to previous protocols ^16^. The aquaeous phase was subsequently extracted and extensively dialyzed against water to remove residual phenol. Extracts were treated with DNase, RNase, and proteinase K, each overnight. After dialysis, extracts were ultracentrifuged (Beckman Coulter Optima) at 100 000 rpm for two hours. Samples were freeze-dried and resuspended in ultrapure water for analysis by Dionex and SDS gel electrophoresis. For enzymatic extractions of polysaccharides, bacterial pellets were treated with mutanolysin (Sigma) and lysozyme (Sigma) according to published protocols ^17^. Cell debris was removed by centrifugation and extracts were treated with DNase, RNase, and proteinase K, and subsequently dialyzed against water. Following ultracentrifugation, extracts were resuspended in ultrapure water.

SDS gel electrophoresis was performed using pre-cast mPAGE™ 4-20% bis-tris gels (Merck) and 20 µl of PS extracts were mixed in Laemmli buffer and boiled at 95°C for 10 minutes. For silver staining, SDS gels were fixed in 45% ethanol solution containing 0.5% periodic acid (Sigma). After extensive washes in water, gels were submerged in 0.1% silver nitrate solution in water for 15 minutes. Gels were rinsed in water and developed in 3% sodium carbonate solution with added formaldehyde to fix the silver dye. Developer was neutralized by addition of citric acid (Sigma). For Alcian blue staining, gels were stained in 0.1% Alcian blue solution in 40% ethanol/5% acetic acid for 1 hour and destained in acetic acid solution ^18^. For Stains-all staining, gels were fixed in 50% ethanol/10% acetic acid solution for 1 hour and extensively washed in water. A 0.1% Stains-all solution was prepared in water and gels were stained in the dark for 30 minutes. Gels were subsequently destained in water.

For analysis of monosaccharides. monosaccharides were first released by acid hydrolysis (TFA 4 N, 4 hours at 100°C or HCl 6 N, 6 hours at 100°C). After vacuum drying of the hydrolysate, monosaccharides were identified and quantified by high performance anion exchange chromatography (HPAEC) with a pulsed electrochemical detector and an anion exchange column (CarboPAC PA-1, 4.6 x 250 mm, Dionex) using 18 mM NaOH as mobile phase at a flow rate of 1 mL/min; glucosamine and galactosamine were used as standards ^19^.

### Isolation of human monocyte-derived macrophages

Ficoll gradient centrifugation (Lympholyte®, Cedarlane) was performed to isolate human peripheral blood mononuclear cells (PMBCs) from freshly extracted blood from healthy donors. To isolate CD14+ cells, anti-hCD14 magnetic beads were used and isolated cells were differentiated to human monocyte-derived macrophages (hMDMs) by addition of 50 ng/ml rhM-CSF (R&D Systems) to XVivo medium (Lonza). After 3 days, medium was exchanged, and cells were differentiated for another 3 days in fresh XVivo supplemented with rhM-CSF. All donors gave written consent under the agreement C-CPSL UNT –No. 15/EFS/023 between the Institut Pasteur and EFS (L’Établissement français du sang), in accordance with articles L1243-4 and R1243-61 of the French Public Health Code and approved by the French Ministry of Science and Technology. Supply and handling of human blood cells followed official guidelines of the agreement between the Institut Pasteur, EFS, and the regulation of blood donation in France.

### ELISA of human pro-inflammatory cytokines

Differentiated hMDMs were seeded at 3x10^4^ cells per well in XVivo (Lonza) and infected with different bacterial strains at an MOI of 10. Cell supernatants were collected at 24 hours post-infection and centrifuged to remove cell debris and extracellular bacteria. The cleared cellular supernatants were rapidly frozen on dry ice and stored at -80 °C until cytokine measurements were performed. ELISA was performed using the SP-X CorPlex™ Human Cytokine Panel 1 (Quanterix), a sandwich-based multiplex ELISA with higher sensitivity and broad range of detection of 10 human pro-inflammatory cytokines. The SP-X array was performed from 6 independent experiments according to the manufacturer’s instructions, including triplicate standard wells and duplicate wells for each sample. The data was analyzed using the proprietary SP-X analysis software.

### ELISA of murine pro-inflammatory cytokines

For ELISA experiments, BMDMs and BMDCs were seeded in 24-well plates (5x10^5^ cells/well). Infections were performed in RPMI 1640 supplemented with 10% FBS. At the indicated time points, the supernatants were collected for cytokine determination using ELISA kits according to the manufacturer’s recommendations (R&D and BD Bioscience).

## Supporting information

Supplementat Figures and Tables for Schmidt et al

## Data Availability

The sequence reads of the RNAseq libraries of the WT and the capsule mutant strain have been deposited in the NCBI Gene Expression Omnibus (GEO) database (Accession number pending).

## Acknowledgements

Work in the C.B. laboratory is financed by the Institut Pasteur and funding has been received from the French Government (grants ANR-10-LABX-62-IBEID and ANR-15-CE17-0014-03 to C.B.) and the “Fondation pour la Recherche Médicale” (grant EQU201903007847 to C.B.). S.S. is a scholar in the Pasteur-Paris University (PPU) International Ph.D. program and received stipends from the Institut Pasteur and the “Fondation pour la Recherche Médicale” (FDT202204015116). We thank the Institut Pasteur UTechs CB platform for cytometry and biomarkers for assistance with SP-X ELISA measurements. We gratefully acknowledge the kind financial support of the Institut Pasteur (Paris) and the Région Ile-de-France (program DIM1Health). Further, we want to thank Hayley Newton for kindly gifting us the *Llo* Δ*dotB* mutant and David Bikard for sfGFP. Work in the D.S.Z. laboratory is financed by The São Paulo Research Foundation (FAPESP grant 2019/11342-6).

## Author contributions

SS, SM, LGV, PE, DPAM, AG, CR, MS, MMN, and TF performed experiments and data analysis. SS, SM, TF and CB designed the experiments. SS and CB wrote the original draft. DSZ and CB generated funding for this study. All authors approved the final version.

