## Supplementat Figures and Tables for Schmidt et al for "The unique *Legionella longbeachae* capsule favors intracellular replication and immune evasion"

**Figure S1:** The capsule cluster is conserved among different *L. longbeachae* strains.
**Figure S2:** The *L. longbeachae* WT and capsule mutant grow at similar rates in BYE medium.
**Figure S3:** Quantification of capsule complementation by TEM and growth phase dependent expression of the capsule in the *L. longbeachae* WT

**Figure S4:** Characterization and visualization of *L. longbeachae* cell wall polysaccharides.

**Figure S5:** RNAseq data of *L. longbeachae* WT vs. capsule mutant in E and PE phase

**Figure S6:** The *L. longbeachae* capsule is transcribed upon infection of *Acanthamoeba castellanii*.

**Figure S7:** The *L. longbeachae* capsule mutant replicates less in lungs of infected mice.

**Figure S8:** The *L. longbeachae* WT and capsule mutant replicate to similar levels in THP-1 cells and murine BMDMs

**Figure S9:** Growth of *L. longbeachae* WT and capsule mutant strains under salt stress, compared to *L. pneumophila*

**Table S1** List of the highest hits of the *L. longbeachae* capsule cluster genes against the NCBI database (Trembl)

**Table S2 CPS gene identitiy among *L. longbeachae* strains compared to NSW150**

**Table *S*4:** Bacteria and plasmids used in this study

**Table S5:** Primers used in this study

### SUPPLEMENTARY FIGURES

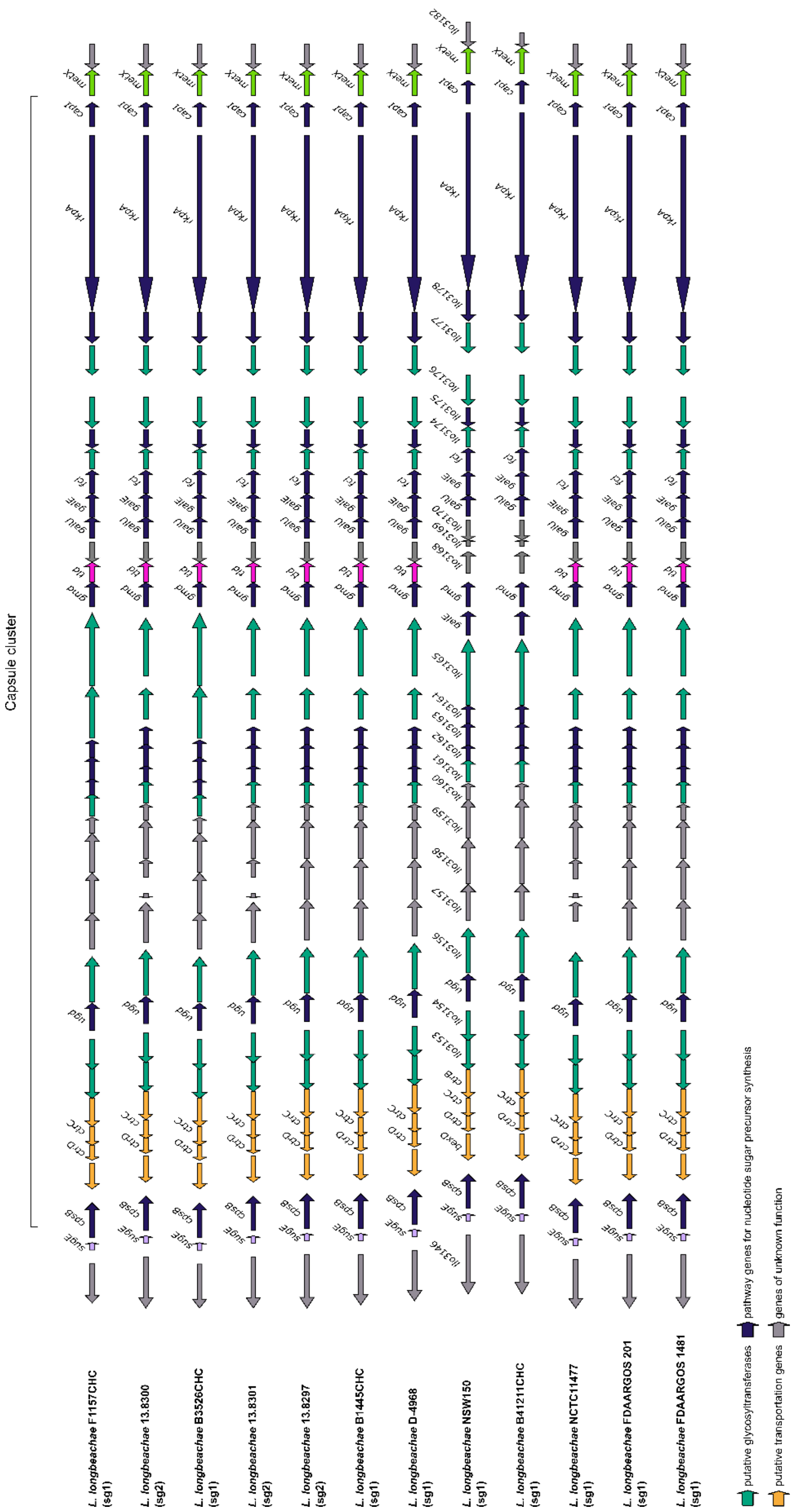


**Figure S1: The capsule cluster is conserved among different *L. longbeachae* strains.** Visualization of the gene content of the cluster capsule in serogroup 1 and serogroup 2

*Legionella longbeachae* strains using GeneSpy[1]. Genes coding for the capsule plus two flanking genes on either side are represented. To obtain homogeneous and comparable annotations, all the genome sequences were reannotated using PROKKA. Based on the new annotation files, gene names and biochemical functions are used to infer families, and a color is attributed for each one.

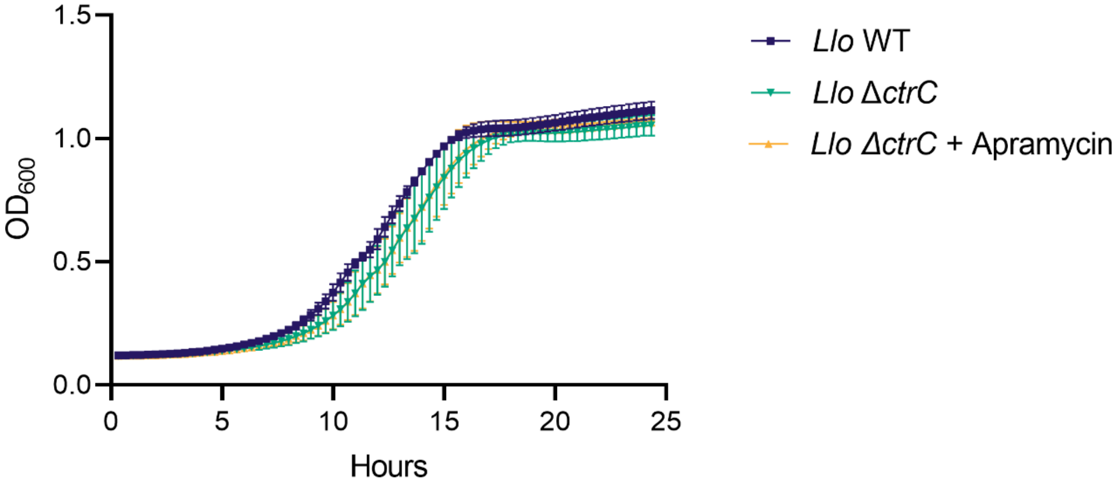


**Figure S2: The *L. longbeachae* WT and capsule mutant grow at similar rates in BYE medium.** *Llo* WT or ΔctrC ± apramycin were grown in BYE medium at 37°C and $OD_{600}$ was followed using a BioTek™ Synergy Plate Reader 2. Data show means ± SD of n=3 independent experiments.

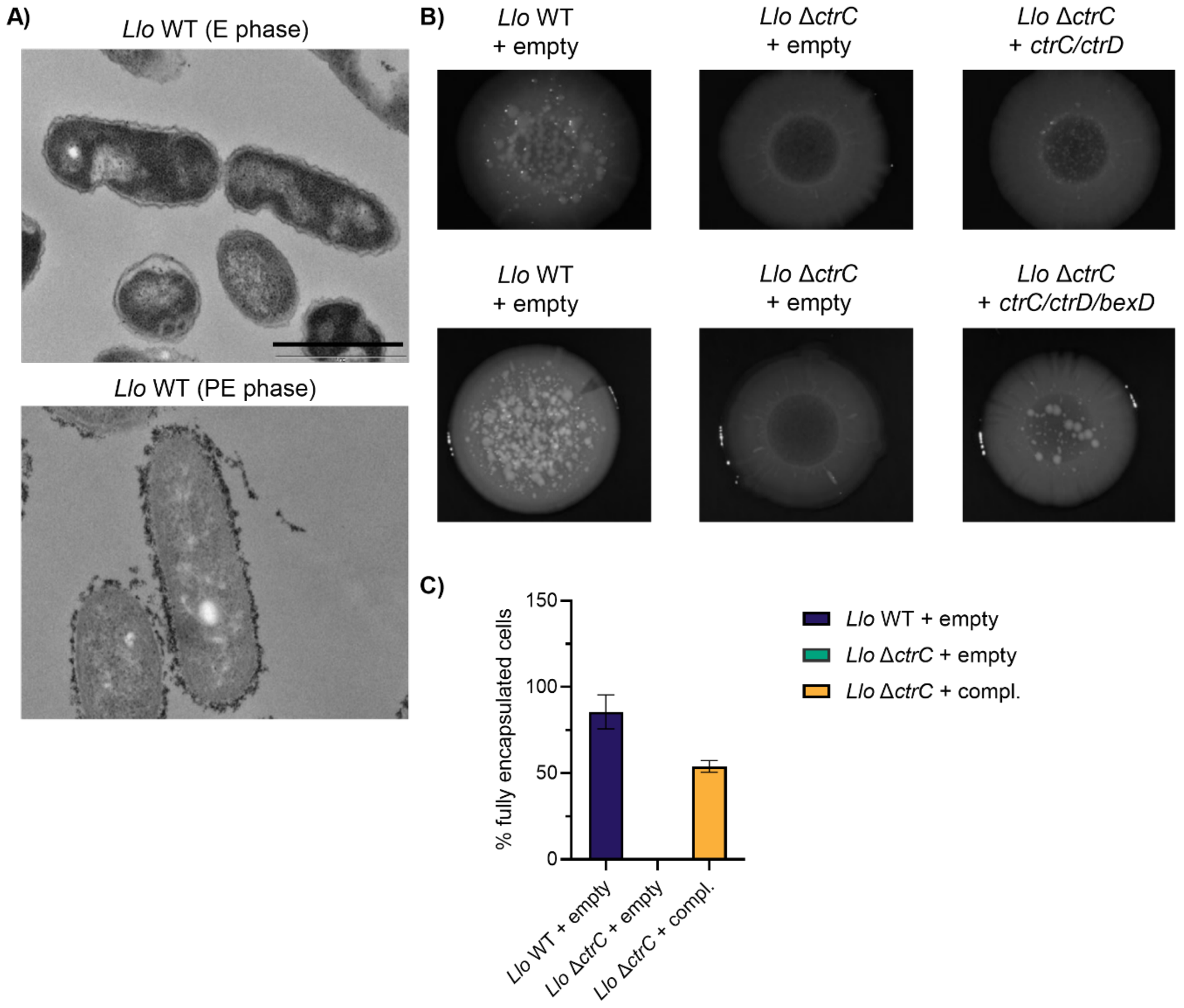


**Figure S3: Quantification of capsule complementation by TEM and growth phase dependent expression of the capsule in the *L. longbeachae* WT. A)** *Llo* WT bacteria were grown to E phase ($OD_{600}$ 2.0-2.5) or PE phase ($OD_{600}$ 3.7-4.2), fixed and stained with cationized ferritin for TEM imaging. Scale bar = 1µm. **B)** Quantification of encapsulated bacteria from TEM images presented in Figure 2B. **C)** Macrocolonies of *Llo* WT or Δ*ctrC* harboring an empty control plasmid (pBCKS) or the complementation plasmids (SSM073 or SSM083). Cells were grown to PE phase and 10 µl were spotted onto BCYE plates for 6 days. Colonies were imaged using a Leica M80 Stereo Microscope with top light and 1x magnification.

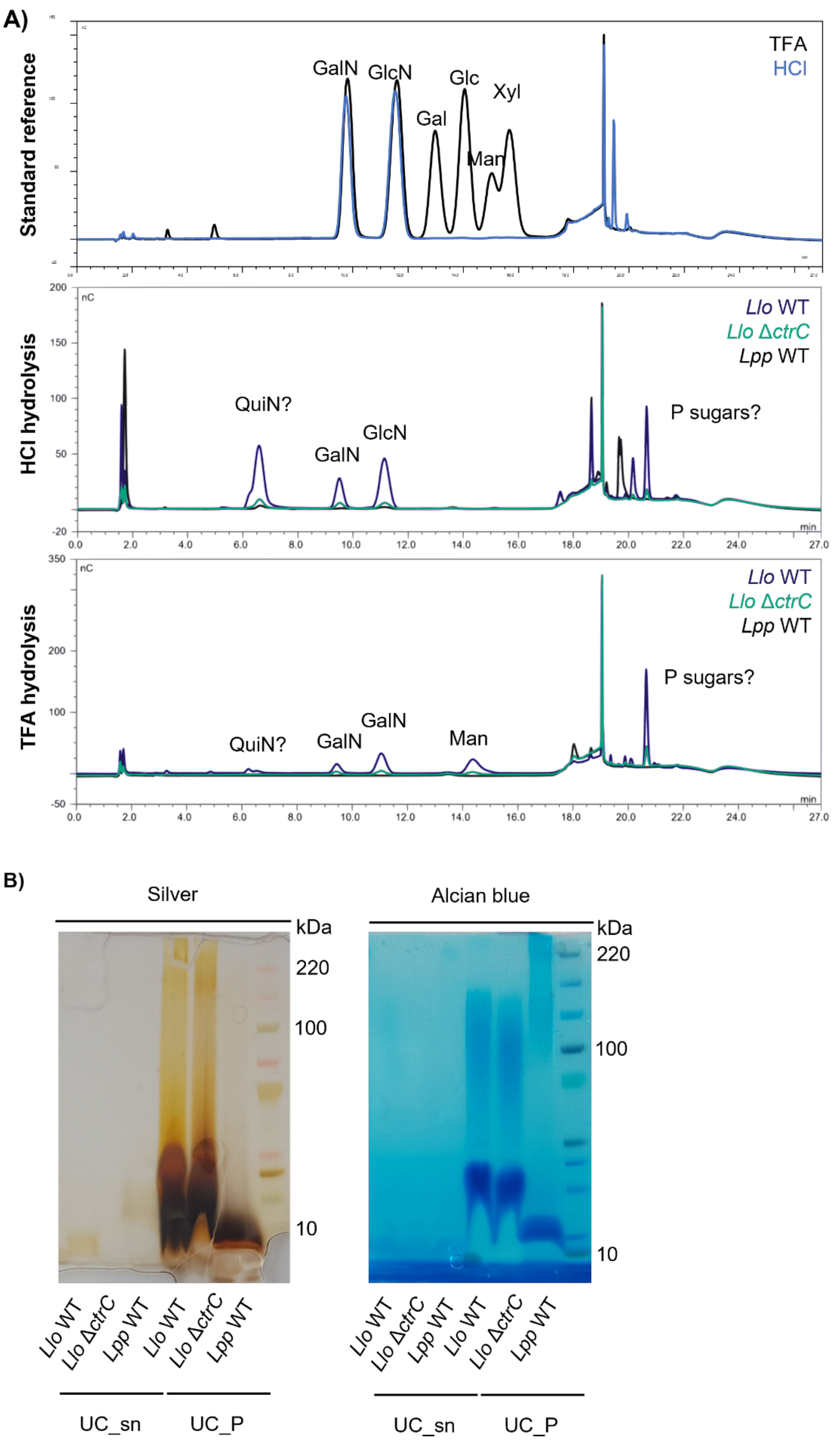


**Figure S4: Characterization and visualization of *L. longbeachae* cell wall polysaccharides.** **A)** Elution profiles of High-Performance Anion Exchange Chromatography. Top graph, standard reference hydrolyzed with TFA. Middle and bottom panel, bacterial phenol extracts hydrolyzed with HCl and TFA. Blue line, *Llo* WT; red line, *Llo* Δ*ctrC*; black line, *Lpp* WT. **B)** SDS gel electrophoresis of phenol extracts stained with silver nitrate or Alcian blue. QuiN, quinovosamine; GalN, galactosamine; GlcN, glucosamine; Gal, galactose; Glc, glucose; Man, mannose; Xyl, xylose; UC_sn, supernatants after ultracentrifugation; UC_P, pellets after ultracentrifugation.

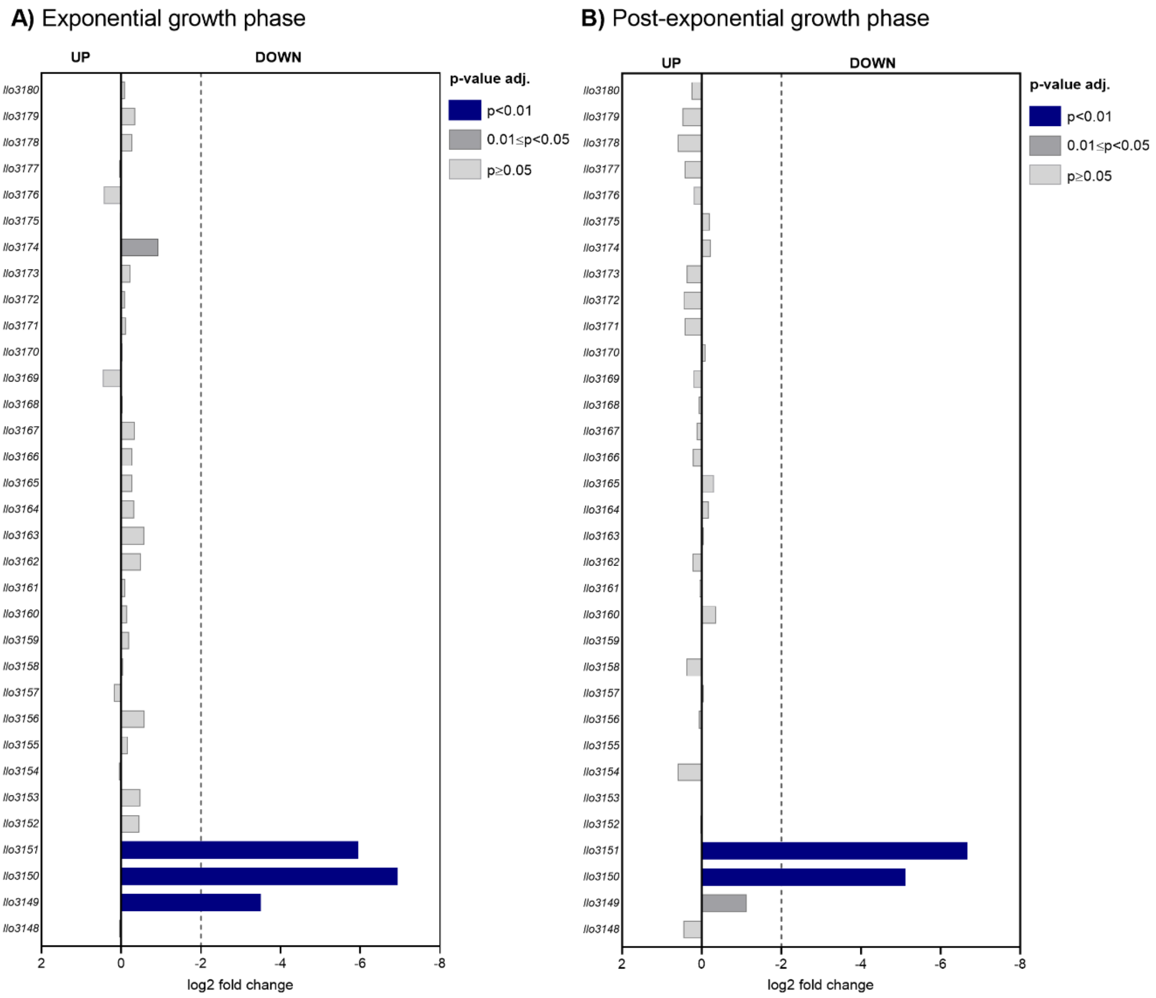


**Figure S5: RNAseq data of *L. longbeachae* WT vs. capsule mutant in E and PE phase (WT is the reference). A)** Comparison of E phase bacteria. **B)** Comparison of PE phase bacteria. WT vs. Δ*ctrC* (WT is the reference); consider relevant genes with log2 fold change of ±2 and adjusted p value ≤ 0.05. n=4 independent experiments.

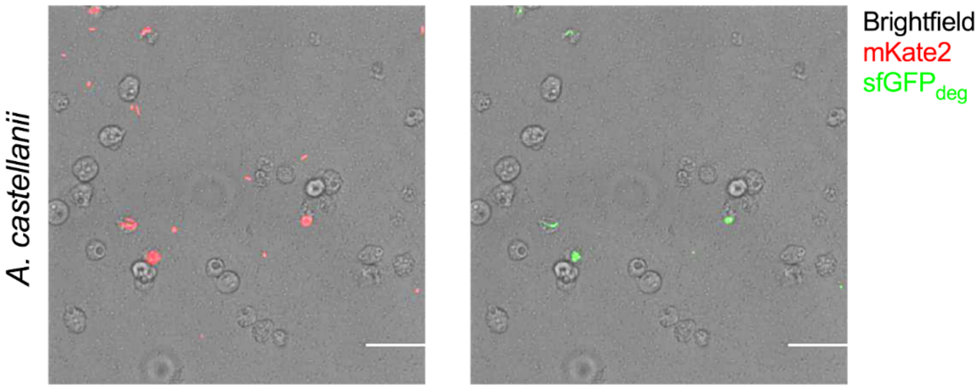


**Figure S6: The *L. longbeachae* capsule is transcribed upon infection of *Acanthamoeba castellanii.*** *A. castellanii* was infected with *Llo* WT bacteria harboring the dual reporter plasmid (pSS017) at MOI 10 and 37°C for 1 hour. Cells were imaged 24 hours post-infection using an EVOS inverted digital microscope. Scale bar = 100 µm.

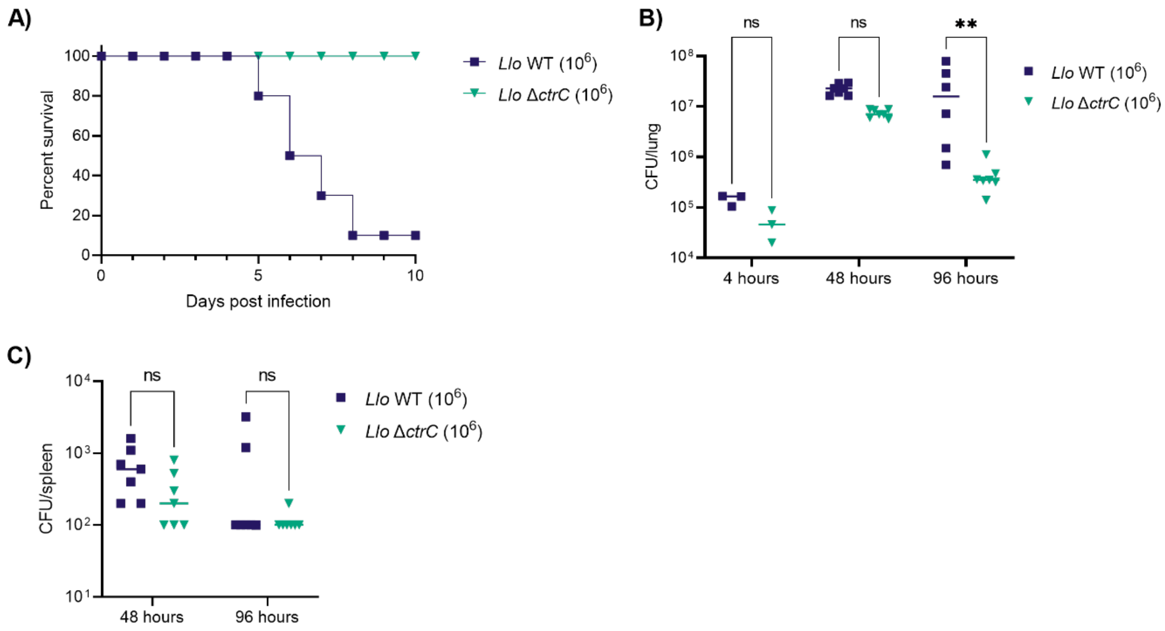


**Figure S7: The *L. longbeachae* capsule mutant replicates less in lungs of infected mice. A)** Female C57BL/6 mice were infected with $10^6$ bacteria and survival was monitored over ten days. 9 mice per group. **B)** CFUs from lungs of infected mice were plated at 4, 48, and 96 hours post-infection. Dots represent number of animals per group. **C)** CFUs from spleens of infected mice were plated at 48 and 96 hours post-infection. Dots represent number of animals per group. Statistical analysis was performed by two-way ANOVA with Tukey's post-test. ns, non-significant; **, $p \leq 0.01$.

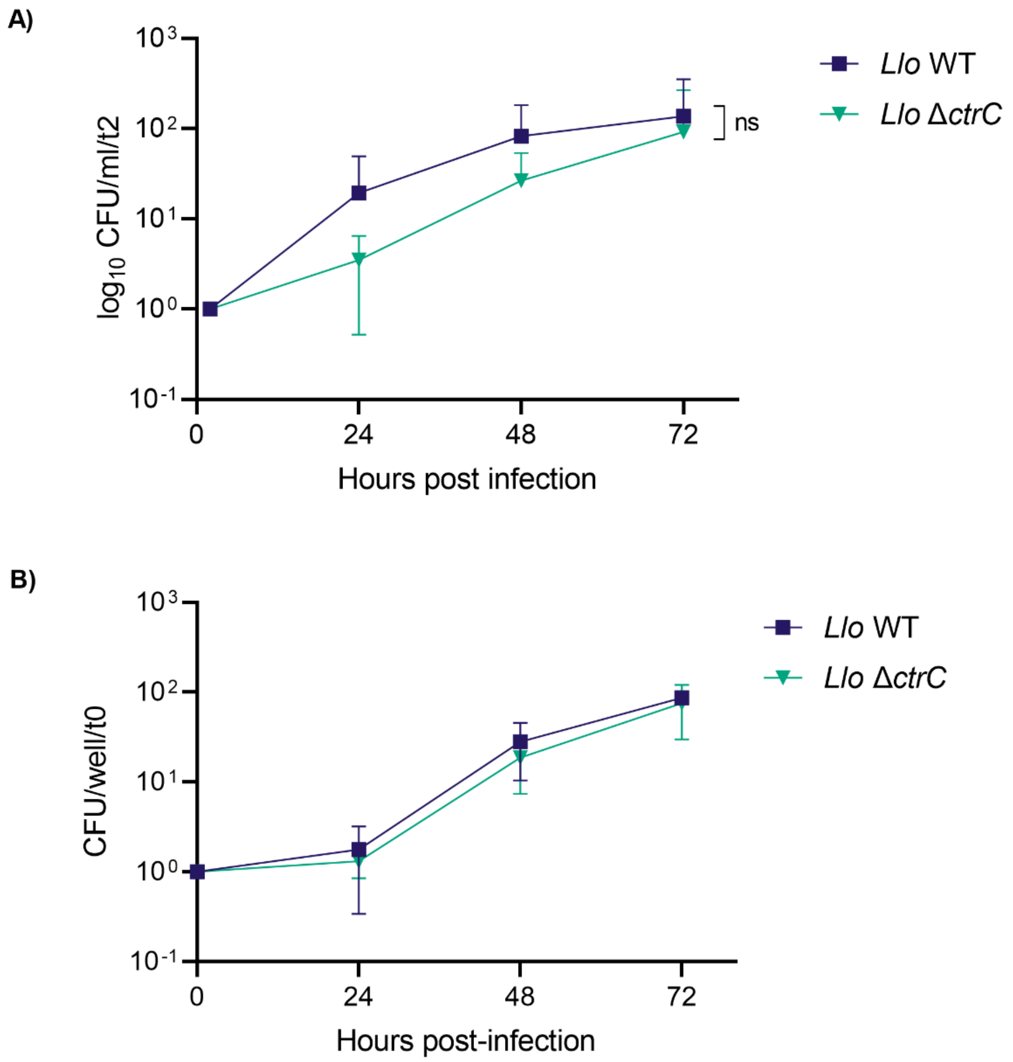


**Figure S8: The *L. longbeachae* WT and capsule mutant replicate to similar levels in THP-1 cells and murine BMDMs. A)** Differentiated THP-1 cells were infected at MOI 10 for 1 hour and treated with gentamycin to kill extracellular bacteria. CFUs were plated every 24 hours and normalized to the input control. Data show means ± SD of n=6 independent experiments **B)** Bone marrow-derived macrophages (BMDMs) were infected at MOI 10 and CFUs plated every 24 hours normalized to the input control. Data show means ± SD of n=3 independent experiments.

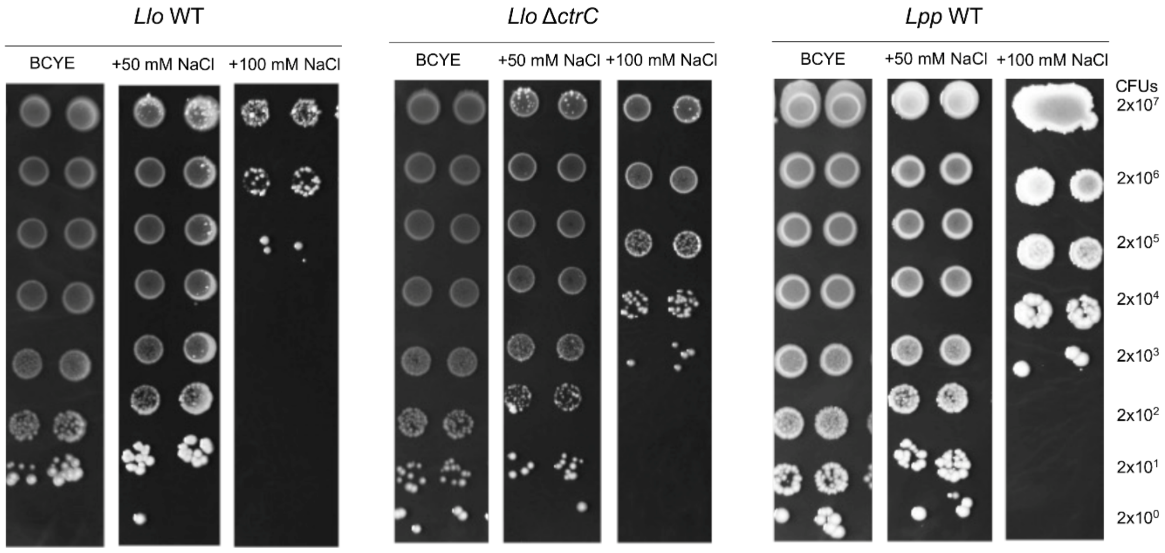


**Figure S9: Growth of *L. longbeachae* WT and capsule mutant strains under salt stress, compared to *L. pneumophila*.** Cells were grown to PE phase (OD600 3.7-4.2) in BYE medium and spotted onto BCYE plates ± NaCl. Representative images of n=3 independent experiments.

**Table S4: Bacteria and plasmids used in this study.**

| **Bacteria** | | |
|---|---|---|
| **Name** | **Source** | **Additional information** |
| *E. coli* DH5α | Invitrogen | Ref. 18265017 |
| *L. longbeachae* strain NSW150 | [4] | Wild type strain, serogroup 1 |
| *L. longbeachae* Δ*ctrC* | This study | KO of *ctrC* in NSW150 background |
| *L. longbeachae* Δ*dotB* | [20] | Genomic deletion of *dotB* |
| *L. pneumophila* strain Paris | [21] | Wild type strain, serogroup 1 |
| **Plasmids** | | |
| **Name** | **Source** | **Additional information** |
| pGEM®-T easy vector | Promega | Cloning vector, $Amp^R$ |
| pBCKS+ | Stratagene | lacZ, $Cm^R$ |
| TOPO-mKate2 | Addgene | Ref. 68441, mKate2, $Kan^R$ |
| pLAW344 | [6] | Suicide plasmid incl. *sacB* cassette, $Cm^R$ |
| pLGV012 | This study | pLAW344-*ctrC*::Apramycin |
| pXDC61 | [22] | N-terminal blaM, $Cm^R$ |
| pSSM012 | [23] | pXDC61-blaM-RomA |
| pSSM073 | This study | Complementation of *ctrC*, including *ctrD,* under the control of the native capsule promoter $prom_{cap}$, $Cm^R$ |
| pSSM083 | This study | Complementation of *ctrC*, including *ctrD* and *bexD,* native capsule promoter $prom_{cap}$, $Cm^R$ |
| pSS016 | This study | Dual reporter without $prom_{cap}$, $Cm^R$ |
| pSS017 | This study | Dual reporter incl. $prom_{cap}$, $Cm^R$ |

### Table S5: Primers used in this study.

| Oligo name | Sequence | Direction |
|---|---|---|
| Apra_as | CCCTCCAACGTCATCTCGTTCTC | reverse |
| Apra_s | CATCAGCAAAAGGGGATGATAAGTTT | forward |
| P1 | TCCCGAGCTCAGTGAAGTCT | forward |
| P2 | AAACTTATCATCCCCTTTTGCTGATGACGCACCCATTACTCCATTC | reverse |
| P3 | TTGGGAAAACGCTCAGAAAC | forward |
| P4 | GAGAACGAGATGACGTTGGAGGGGGGCTCAAGAGCAAACCATA | reverse |
| SSM_79 | GGATCCCTCTCTCCCTTTACGGCGGTAT | forward |
| SSM_80 | TTGTTAGGAAATGCACATTTTGCATCGACACCAATCCTTAATGTCAAAA | reverse |
| SSM_82 | GGTACCTTAAGTTTGCTTGTTGTAAAATTCG | forward |
| SSM_85 | ATGCAAAATGTGCATTTCCTAAC | reverse |
| SSM_86 | GGCCGCTCTAGAACTAGTGGATCCCTCTCTCCCTTTACGGCG | forward |
| SSM_087 | ACTAAAGGGAACAAAAGCTGGGTACCTTAAGTTTGCTTGTTGTAAAATTCG | reverse |
| SSM_113 | GGAAGCTTACGAATTTTACAACAAGCAAACTTAAATAAGTTTTTACAGGAGTTAATTTTT AAAGTGATAAAG | forward |
| SSM_114 | ACTAAAGGGAACAAAAGCTGGGTACCCTAAGCCCTTATAACCTGTGTTGC | reverse |
| SSO_025 | GGCCGCTCTAGAACTAGTGGATCCACTCTCTCCCTTTACGGCG | forward |
| SSO_049 | CAGTTCTTCACCTTTACTCATCGACACCAATCCTTAATGTCAAAATTTC | reverse |
| SSO_050 | CTACAAACCAGGCATCAAATAGTAAAAGCTTACTCTCTCCCTTTACGGCG | forward |
| SSO_049 | CAGTTCTTCACCTTTACTCATCGACACCAATCCTTAATGTCAAAATTTC | reverse |
| SSO_051 | ATGAGTAAAGGTGAAGAACTGTTCAC | forward |
| SSO_052 | CGCCGTAAAGGGAGAGAGTAAGCTTTTACTATTTGATGCCTGGTTTGTAG | reverse |
| SSO_053 | GAACAAAAGCTGGGTACCTTACTATTTGATGCCTGGTTTGTAG | reverse |
| SSO_045 | TGGATGAACTCTACAAACCAGGCATCAAAGCTGCTAATGATGAAAATTATGCTGATGCT | forward |
| SSO_046 | GAGGTCGACGGTATCGATAAGCTTTTACTAAGAAGCATCAGCATAATTTTCATCATTAG C | reverse |
| SSO_047 | CATTTTTATCTATAATATTGGCAAATCTGCTACAGCGCTGCTAATGATGAAAATTATGCT G | forward |
| SSO_048 | CATTATTATTTATCCTGATTGATTCAGGTTATTTACTAAGAAGCATCAGCATAATTTTCA TCA | reverse |
| SSO_065 | GCGGTGGCGGCCGCTTTACAGCTAGCTCAGTCCTAGGTATTATGCTAGCGAATTCGCTA GATTTAAGAAGGAGATATACATATGGTGAGCGAGCTGATTAAG | forward |
| SSO_066 | GGAGAGAGTGGATCCTCATCTGTGCCCCAGTTTGC | reverse |
| SSO_621 | GATGATGGATCCTGACTAACTAGCAGTAAAGGTGAAGAACTGTTCACC | forward |
| SSO_622 | GATGATAAGCTTTTAAGAAGCATCAGCATAATTTTCATC | reverse |
| SSO_623 | GATGATAAGCTTTGACTAACTAGCAGTAAAGGTGAAGAACTGTTCACC | forward |
| SSO_624 | GATGATGGTACCTTAAGAAGCATCAGCATAATTTTCATC | reverse |
