## Supplemental Table S1 for "The unique *Legionella longbeachae* capsule favors intracellular replication and immune evasion"

**Supplementary Table 1** List of the highest hits of the *L. longbeachae* capsule cluster genes against the NCBI database (Trembl).

**Genomic Object Editor: llo3148**

| PB id | Ident % | Eval | Gene | Description | EC number | Keywords | Organism |
|---|---|---|---|---|---|---|---|
| Q48462 | 56.81 | 0 | manC | Mannose-1-phosphate guanylyltransferase | 2.7.7.13 | Capsule biogenesis/degradation, [illegible] | Klebsiella pneumoniae |
| Q01410 | 57.08 | 0 | manC | Mannose-1-phosphate guanylyltransferase | 2.7.7.13 | GTP-binding, Lipopolysaccharide [illegible] | Salmonella montevideo |
| P37753 | 56.38 | 0 | manC | Mannose-1-phosphate guanylyltransferase | 2.7.7.13 | GTP-binding, Lipopolysaccharide [illegible] | Escherichia coli |
| P26404 | 56.13 | 0 | rfbM | Mannose-1-phosphate guanylyltransferase RfbM | 2.7.7.13 | Direct protein sequencing, GTP-binding, [illegible] | Salmonella typhimurium (strain LT2 / SGSC1412 / ATCC 700720) |
| Q8X7P1 | 56.14 | 0 | manC1 | Mannose-1-phosphate guanylyltransferase 1 | 2.7.7.13 | GTP-binding, Nucleotide-binding, [illegible] | Escherichia coli O157:H7 |
| P07874 | 55.25 | 0 | algA | Mannose-6-phosphate isomerase / Mannose-1-phosphate [illegible] | 5.3.1.8, 2.7.7.13 | Alginate biosynthesis, Cobalt, Direct protein [illegible] | Pseudomonas aeruginosa (strain ATCC 15692 / DSM 22644 / CIP 104116 / [illegible] |
| P24174 | 55.93 | 0 | manC | Mannose-1-phosphate guanylyltransferase | 2.7.7.13 | Capsule biogenesis/degradation, [illegible] | Escherichia coli (strain K12) |
| P26340 | 55.93 | 0 | manC | Mannose-1-phosphate guanylyltransferase ManC | 2.7.7.13 | Capsule biogenesis/degradation, [illegible] | Salmonella typhimurium (strain LT2 / SGSC1412 / ATCC 700720) |
| B0RVK6 | 56.68 | 0 | xanB | Mannose-6-phosphate isomerase / Mannose-1-phosphate guanylyl [illegible] | 5.3.1.8, 2.7.7.13 | Exopolysaccharide synthesis, GTP-binding, [illegible] | Xanthomonas campestris pv. campestris (strain B100) |
| O85342 | 56.03 | 0 | manC2 | Mannose-1-phosphate guanylyltransferase 2 | 2.7.7.13 | GTP-binding, Lipopolysaccharide [illegible] | Escherichia coli O157:H7 |
| P0C7J3 | 56.47 | 0 | xanB | Mannose-6-phosphate isomerase / Mannose-1-phosphate guanylyl [illegible] | 5.3.1.8, 2.7.7.13 | Exopolysaccharide synthesis, GTP-binding, [illegible] | Xanthomonas campestris pv. campestris (strain ATCC 33913 / DSM [illegible] |
| Q07024 | 56.01 | 0 | rfbA | Putative mannose-1-phosphate guanylyltransferase | 2.7.7.13 | GTP-binding, Lipopolysaccharide [illegible] | Vibrio cholerae serotype O1 (strain ATCC 39315 / El Tor Inaba N16961) |

**Genomic Object Editor: llo3149**

| PB id | Ident % | Eval | Gene | Description | EC number | Keywords | Organism |
|---|---|---|---|---|---|---|---|
| D3HMB3 | 100 | 0 | bexD | Capsule polysaccharide export protein bexD | _ | Reference proteome, Signal | Legionella longbeachae serogroup 1 (strain NSW150) |
| A0A1P8FL76 | 52.28 | 2E-127 | _ | Capsular biosynthesis protein | _ | Reference proteome, Signal, Transport | Betaproteobacteria bacterium GR16-43 |
| A0A1I3G2S8 | 54.64 | 1E-126 | _ | Polysaccharide export outer membrane protein | _ | Signal | Collimonas sp. OK307 |
| A0A1T4QGS8 | 51.37 | 5E-123 | _ | Polysaccharide export outer membrane protein | _ | Signal | Geobacter thiogenes |
| B3E8Y1 | 52.47 | 7E-123 | _ | Polysaccharide export protein | _ | Reference proteome, Signal, Transport | Geobacter lovleyi (strain ATCC BAA-1151 / DSM 17278 / SZ) |
| Q39JA8 | 51.61 | 7E-123 | _ | Polysaccharide export protein | _ | Membrane, Signal, Transmembrane, Transmembrane helix | Burkholderia lata (strain ATCC 17760 / DSM 23089 / LMG 22485 / NCIMB 9086 / R18194 / 383) |
| A0A1I9YRW4 | 51.76 | 6E-122 | _ | Capsular biosynthesis protein | _ | Signal, Transport | Paraburkholderia sprentiae WSM5005 |
| A0A2S5SX59 | 51.32 | 1E-121 | _ | Capsular biosynthesis protein | _ | Reference proteome, Signal, Transport | Zhizhongheella caldifontis |
| A0A0J9DW18 | 51.1 | 1E-121 | _ | Capsular biosynthesis protein | _ | Signal | Ralstonia sp. MD27 |
| Q7BMG8 | 50.68 | 4E-120 | wcbC | Capsular biosynthesis protein | _ | Signal | Burkholderia pseudomallei |
| A0A0F7R2Q8 | 42.63 | 1E-98 | cpx15D | Capsular polysaccharide export protein D | _ | Signal | Actinobacillus pleuropneumoniae |
| Q44132 | 42.74 | 5E-98 | cpxD | CpxD | _ | Signal | Actinobacillus pleuropneumoniae |
| A0A059WCS1 | 42.74 | 2E-97 | cpxD | Sugar ABC transporter substrate-binding protein CpxD | _ | Signal | Actinobacillus pleuropneumoniae serovar 8 str. 405 |
| A0A059WNF9 | 41.84 | 3E-97 | cpxD | Sugar ABC transporter substrate-binding protein CpxD | _ | Signal | Actinobacillus pleuropneumoniae |
| Q9RP76 | 42.63 | 1E-95 | cpxD | CpxD | _ | Signal | Mannheimia haemolytica |
| Q7WS64 | 40.53 | 3E-93 | bexD | BexD | _ | Signal | Haemophilus influenzae |
| Q714V0 | 40.79 | 2E-92 | bexD | BexD | _ | Signal | Haemophilus influenzae |
| Q9L9L6 | 41.42 | 3E-87 | cexD | CexD | _ | Signal | Pasteurella multocida |
| Q8KS22 | 39.94 | 1E-86 | ctrA | Capsular transport protein | _ | Signal | Neisseria meningitidis |

**Genomic Object Editor: llo3150**

| PB id | Ident % | Eval | Gene | Description | EC number | Keywords | Organism |
|---|---|---|---|---|---|---|---|
| D3HMB4 | 100 | 1,00E-153 | ctrD | Capsule polysaccharide export ATP-binding protein ctrD (Capsular-polysaccharide-transporting ATPase) | 3.6.3.38 | ATP-binding, Hydrolase, Nucleotide-binding, Reference proteome | Legionella longbeachae serogroup 1 (strain NSW150) |
| A0A1I9YRU6 | 67.59 | 7,00E-108 | _ | ATP-binding protein | _ | ATP-binding, Cell inner membrane, Cell membrane, Membrane, Nucleotide-binding, Translocase | Paraburkholderia sprentiae WSM5005 |
| A0A1V2XQM1 | 68.52 | 3,00E-107 | _ | ATP-binding protein | _ | ATP-binding, Cell inner membrane, Cell membrane, Membrane, Nucleotide-binding, Translocase, Transport | Burkholderia cenocepacia |
| A0A2U9SFH7 | 68.06 | 2,00E-106 | _ | ABC transporter ATP-binding protein | _ | ATP-binding, Cell inner membrane, Cell membrane, Membrane, Nucleotide-binding, Translocase, Transport | Burkholderia sp. JP2-270 |
| B2UAD6 | 68.22 | 1,00E-105 | _ | ABC transporter related | _ | ATP-binding, Cell inner membrane, Cell membrane, Membrane, Nucleotide-binding, Reference proteome, Translocase | Ralstonia pickettii (strain 12J) |
| U3GDS8 | 68.22 | 1,00E-105 | _ | ABC transporter domain-containing protein | _ | ATP-binding, Cell inner membrane, Cell membrane, Membrane, Nucleotide-binding, Translocase | Ralstonia sp. 5_2_56FAA |
| A0A117DUD2 | 68.69 | 8,00E-105 | _ | ABC transporter-like protein | _ | ATP-binding, Cell inner membrane, Cell membrane, Membrane, Nucleotide-binding, Translocase | Ralstonia sp. NT80 |
| S9RTK8 | 68.69 | 8,00E-105 | _ | ATP-binding protein | _ | ATP-binding, Cell inner membrane, Cell membrane, Membrane, Nucleotide-binding, Translocase | Ralstonia sp. AU12-08 |
| A0A1G8E0F9 | 67.76 | 4,00E-104 | _ | Capsular polysaccharide transport system ATP-binding protein | _ | ATP-binding, Cell inner membrane, Cell membrane, Membrane, Nucleotide-binding, Translocase | Paraburkholderia phenazinium |

| | | | | | | | |
|---|---|---|---|---|---|---|---|
| U2FWZ6 | 67.45 | 9,00E-104 | _ | Capsular polysaccharide ABC transporter, ATP-binding protein KpsT | _ | ATP-binding, Cell inner membrane, Cell membrane, Membrane, Nucleotide-binding, Translocase, Transport | Burkholderia sp. AU4i |
| A0A1N6LM37 | 67.29 | 4,00E-103 | _ | Capsular polysaccharide transport system ATP-binding protein | _ | ATP-binding, Cell inner membrane, Cell membrane, Membrane, Nucleotide-binding, Translocase, Transport | Burkholderia sp. GAS332 |
| K7QQ86 | 63.85 | 6,00E-97 | ctrD | CtrD | _ | ATP-binding, Nucleotide-binding | Kingella kingae |
| B3FHD0 | 63.85 | 1,00E-94 | ctrD | ATP-binding cassette domain-containing protein | _ | ATP-binding, Nucleotide-binding | Neisseria meningitidis |
| B3FHE0 | 63.38 | 9,00E-94 | ctrD | ATP-binding protein | 3.6.3.38 | ATP-binding, Hydrolase, Nucleotide-binding | Neisseria meningitidis |
| Q9S6K7 | 61.5 | 6,00E-92 | cpxA | Capsule polysaccharide export transport system ATP-binding protein | _ | ATP-binding, Nucleotide-binding | Actinobacillus pleuropneumoniae |
| Q7B3Y6 | 60.28 | 9,00E-91 | bexA | BexA | _ | ATP-binding, Nucleotide-binding | Haemophilus influenzae |
| Q44135 | 61.03 | 9,00E-91 | cpxA | CpxA | _ | ATP-binding, Nucleotide-binding | Actinobacillus pleuropneumoniae |
| O85464 | 60.09 | 1,00E-90 | hexA | ABC transporter ATP-binding protein | _ | ATP-binding, Nucleotide-binding | Pasteurella multocida |
| Q9L9L9 | 60.56 | 3,00E-90 | cexA | CexA | _ | ATP-binding, Nucleotide-binding | Pasteurella multocida |
| Q93UJ9 | 62.25 | 2,00E-89 | wzt2 | Wzt2 | _ | ATP-binding, Cell inner membrane, Cell membrane, Membrane, Nucleotide-binding, Translocase, Transport | Burkholderia pseudomallei |
| Q9RP79 | 60.28 | 2,00E-89 | cpxA | Leukotoxin translocation ATP-binding protein LktB | 7.4.2.5 | ATP-binding, Nucleotide-binding | Mannheimia haemolytica |

**Genomic Object Editor: llo3151**

| PB id | Ident % | Eval | Gene | Description | EC number | Keywords | Organism |
|---|---|---|---|---|---|---|---|
| P32015 | 45.82 | 9,00E-74 | ctrC | Capsule polysaccharide export inner-membrane protein CtrC | _ | Capsule biogenesis/degradation, Cell inner membrane, Polysaccharide transport, Transmembrane | Neisseria meningitidis serogroup B (strain MC58) |
| P57012 | 45.82 | 4,00E-71 | ctrC | Capsule polysaccharide export inner-membrane protein CtrC | _ | Capsule biogenesis/degradation, Cell inner membrane, Polysaccharide transport, Transmembrane | Neisseria meningitidis serogroup A / serotype 4A (strain DSM 15465 / Z2491) |
| P19391 | 40.89 | 1,00E-66 | bexB | Capsule polysaccharide export inner-membrane protein BexB | _ | Capsule biogenesis/degradation, Cell inner membrane, Polysaccharide transport, Transmembrane | Haemophilus influenzae |
| P19390 | 40.4 | 8,00E-66 | bexB | Capsule polysaccharide export inner-membrane protein BexB | _ | Capsule biogenesis/degradation, Cell inner membrane, Polysaccharide transport, Transmembrane | Haemophilus influenzae |
| P22235 | 40 | 6,00E-65 | bexB | Capsule polysaccharide export inner-membrane protein BexB | _ | Capsule biogenesis/degradation, Cell inner membrane, Polysaccharide transport, Transmembrane | Haemophilus influenzae |
| P24584 | 26.64 | 5,00E-23 | kpsM | Polysialic acid transport protein KpsM | _ | Cell inner membrane, Cell membrane, Membrane, Transmembrane, Transmembrane helix, Transport | Escherichia coli |
| P23889 | 26.23 | 8,00E-22 | kpsM | Polysialic acid transport protein KpsM | _ | Cell inner membrane, Cell membrane, Membrane, Transmembrane, Transmembrane helix, Transport | Escherichia coli |

**Genomic Object Editor: llo3152**

| PB id | Ident % | Eval | Gene | Description | EC number | Keywords | Organism |
|---|---|---|---|---|---|---|---|
| P57034 | 41 | 5,00E-90 | ctrB | Capsule polysaccharide export inner-membrane protein CtrB | _ | Capsule biogenesis/degradation, Cell inner membrane, Polysaccharide transport, Transmembrane | Neisseria meningitidis serogroup A / serotype 4A (strain DSM 15465 / Z2491) |
| P22930 | 39.72 | 2,00E-85 | bexC | Capsule polysaccharide export inner-membrane protein BexC | _ | Capsule biogenesis/degradation, Cell inner membrane, Polysaccharide transport, Transmembrane | Haemophilus influenzae |
| P32014 | 38.78 | 2,00E-83 | ctrB | Capsule polysaccharide export inner-membrane protein CtrB | _ | Capsule biogenesis/degradation, Cell inner membrane, Polysaccharide transport, Transmembrane | Neisseria meningitidis serogroup B (strain MC58) |
| P62586 | 25.07 | 7,00E-35 | kpsE | Capsule polysaccharide export inner-membrane protein KpsE | _ | Capsule biogenesis/degradation, Cell inner membrane, Polysaccharide transport, Transmembrane | Escherichia coli |
| P42501 | 25.07 | 2,00E-34 | kpsE | Capsule polysaccharide export inner-membrane protein KpsE | _ | Capsule biogenesis/degradation, Cell inner membrane, Polysaccharide transport, Transmembrane | Escherichia coli |
| P43111 | 23.98 | 2,00E-18 | vexD | Vi polysaccharide export inner-membrane protein VexD | _ | Capsule biogenesis/degradation, Cell inner membrane, Polysaccharide transport, Transmembrane | Salmonella typhi |

**Supplementary Table 1** List of the highest hits of the *L. longbeachae* capsule cluster genes against the NCBI database (Trembl).

| | | | | | | | |
|---|---|---|---|---|---|---|---|
| Q48452 | 20.75 | 0.063 | _ | Putative tyrosine-protein kinase in cps region | 2.7.10.- | ATP-binding, Cell inner membrane, Exopolysaccharide synthesis, Nucleotide-binding, Phosphoprotein, Transferase Transmembrane | Klebsiella pneumoniae |
| Q9F7B1 | 21.72 | 0.25 | wzc | Tyrosine-protein kinase wzc | 2.7.10.- | ATP-binding, Cell inner membrane, Exopolysaccharide synthesis, Nucleotide-binding, Phosphoprotein, Transferase Transmembrane | Salmonella typhimurium (strain LT2 / SGSC1412 / ATCC 700720) |

**Genomic Object Editor: llo3153**

| PB id | Ident % | Eval | Gene | Description | EC number | Keywords | Organism |
|---|---|---|---|---|---|---|---|
| D3HMB7 | 100 | 0 | _ | Putative glycosyl transferase group 1 | _ | Reference proteome, Transferase | Legionella longbeachae serogroup 1 (strain NSW150) |
| F5YHI6 | 25.36 | 2,00E-30 | _ | Glycosyltransferase, family 1 | _ | Coiled coil, Reference proteome, Transferase | Treponema primitia (strain ATCC BAA-887 / DSM 12427 / ZAS-2) |
| A0A2P6CAZ8 | 26.72 | 3,00E-30 | _ | Glycos_transf_1 domain-containing protein | _ | _ | Polaribacter butkevichii |
| A0A193SEV6 | 28.15 | 3,00E-29 | _ | Putative mannosyltransferase B | _ | Glycosyltransferase, Transferase | Klebsiella pneumoniae |
| I9BN45 | 27.16 | 5,00E-29 | _ | Glycos_transf_1 domain-containing protein | _ | Coiled coil | Bacteroides fragilis CL05T12C13 |
| E4VU69 | 26.98 | 3,00E-28 | _ | Glycosyltransferase, group 1 family protein | 2.4.-.- | Coiled coil, Glycosyltransferase, Transferase | Bacteroides fragilis 3_1_12 |
| O84909 | 30.59 | 0.00000000000007 | wbpY | Glycosyltransferase WbpY | _ | Transferase | Pseudomonas aeruginosa |
| Q47594 | 25.25 | 0.000000000007 | mtfB | Mannosyltransferase | _ | Glycosyltransferase, Transferase | Escherichia coli |
| Q9RMT9 | 25.25 | 0.000000000007 | wbdB | WbdB | _ | _ | Klebsiella pneumoniae |
| C8YZ35 | 27.32 | 0.000000009 | wejJ | WejJ | _ | _ | Escherichia coli |
| M4QPQ6 | 31.2 | 0.00000003 | wbdA | Mannosyltransferase | _ | Glycosyltransferase, Transferase | Escherichia coli |
| Q9RMU0 | 34.02 | 0.0000002 | wbdA | WbdA | _ | _ | Klebsiella pneumoniae |
| C8YZ32 | 29.1 | 0.0000002 | wejI | WejI | _ | _ | Escherichia coli |
| Q00481 | 25.59 | 0.0000002 | _ | Glycosyltransferase | _ | Transferase | Salmonella enterica |
| Q93CS4 | 41.54 | 0.000001 | wbaX | Putative glycosyl transferase | _ | Transferase | Shigella boydii |
| Q9LC66 | 31.53 | 0.000002 | wbdA | Mannosyltransferase | _ | Glycosyltransferase, Transferase | Klebsiella pneumoniae |
| Q47593 | 31.53 | 0.000002 | mtfA | Mannosyltransferase A | _ | Glycosyltransferase, Transferase | Escherichia coli |
| M4QN28 | 31.53 | 0.000002 | wbdA | Mannosyltransferase | _ | Glycosyltransferase, Transferase | Escherichia coli |

**Genomic Object Editor: llo3154**

| PB id | Ident % | Eval | Gene | Description | EC number | Keywords | Organism |
|---|---|---|---|---|---|---|---|
| D3HMB8 | 100 | 0 | _ | Putative glycosyl transferase, group 1 | _ | Coiled coil, Reference proteome, Transferase | Legionella longbeachae serogroup 1 (strain NSW150) |
| A0A254RBS0 | 36.96 | 2,00E-76 | _ | Uncharacterized protein | _ | Membrane, Transmembrane, Transmembrane helix | Fibrobacter sp. UWR2 |
| A0A1V4R768 | 32.1 | 1,00E-64 | _ | Glycos_transf_1 domain-containing protein | _ | _ | Candidatus Cloacimonas sp. 4484_140 |
| A0A2E0ZAU5 | 34.06 | 3,00E-56 | _ | Colanic acid biosynthesis glycosyltransferase WcaL | _ | Transferase | Anaerolineaceae bacterium |
| A0A0P9DL51 | 31.5 | 6,00E-52 | _ | Colanic acid biosynthesis glycosyl transferase | _ | Reference proteome, Transferase | Kouleothrix aurantiaca |
| F2NHK4 | 32.51 | 6,00E-51 | _ | Glycosyl transferase group 1 | _ | Reference proteome, Transferase | Desulfobacca acetoxidans (strain ATCC 700848 / DSM 11109 / ASRB2) |
| A0A2V2RIU7 | 31.47 | 2,00E-50 | _ | Colanic acid biosynthesis glycosyltransferase WcaL | _ | Transferase | Acidobacteria bacterium |
| A0A193SBW5 | 28.34 | 9,00E-27 | wclQ | Glycosyltransferase protein | 2.4.1.57 | Glycosyltransferase, Transferase | Klebsiella pneumoniae |
| A4F3K9 | 26.62 | 0.0000000000006 | aerI | Putative glycosyltransferase | _ | Transferase | Planktothrix agardhii NIVA-CYA 126 |
| Q204F0 | 24.43 | 0.00000000006 | cps2G | Cps1/2G | _ | Glycosyltransferase, Transferase | Streptococcus suis |
| Q9RHD0 | 25.96 | 0.00000000007 | wbpU | Glycosyl transferase-like protein | 2.4.1.21 | Glycosyltransferase, Transferase | Pseudomonas aeruginosa |
| Q6L735 | 26.35 | 0.0000000004 | _ | Glycosyltransferase | _ | Glycosyltransferase, Transferase | Streptomyces kanamyceticus |
| Q8KWP7 | 29.7 | 0.000000001 | cps9vG | Capsular polysaccharide biosynthesis protein Cps4H | 2.4.1.21 | Glycosyltransferase, Transferase | Streptococcus pneumoniae |
| D2KXE7 | 22.45 | 0.000000003 | epsG | Putative glycosyltransferase | _ | Transferase | Lactobacillus fermentum |
| Q8GJ89 | 25.38 | 0.000000004 | wbyC | Putative glycosyltransferase | _ | Transferase | Yersinia pseudotuberculosis |
| Q9EVX4 | 25.2 | 0.000000007 | cpsG | Putative hexose transferase | _ | Transferase | Streptococcus salivarius |
| Q00481 | 27.03 | 0.00000001 | _ | Glycosyltransferase | _ | Transferase | Salmonella enterica |

**Genomic Object Editor: llo3155**

| PB id | Ident % | Eval | Gene | Description | EC number | Keywords | Organism |
|---|---|---|---|---|---|---|---|
| D3HMB9 | 100 | 0 | ugd | UDP-glucose 6-dehydrogenase | 1.1.1.22 | NAD, Oxidoreductase, Reference proteome | Legionella longbeachae serogroup 1 (strain NSW150) |
| A0A088U1Y7 | 65.98 | 0 | _ | UDP-glucose 6-dehydrogenase | 1.1.1.22 | NAD, Oxidoreductase | Burkholderia cenocepacia |
| B3QQB3 | 65.46 | 0 | _ | UDP-glucose 6-dehydrogenase | 1.1.1.22 | NAD, Oxidoreductase | Chlorobaculum parvum (strain NCIB 8327) |
| A0A0B0YZU4 | 64.95 | 0 | ugd | UDP-glucose 6-dehydrogenase | 1.1.1.22 | NAD, Oxidoreductase | Escherichia coli |
| D2ZEC2 | 65.21 | 0 | _ | UDP-glucose 6-dehydrogenase | 1.1.1.22 | NAD, Oxidoreductase | Enterobacter cancerogenus ATCC 35316 |
| A0A132F643 | 65.72 | 0 | _ | UDP-glucose 6-dehydrogenase | 1.1.1.22 | NAD, Oxidoreductase | Burkholderia pseudomultivorans |
| A0A2J0NPG5 | 64.95 | 0 | _ | UDP-glucose 6-dehydrogenase | 1.1.1.22 | NAD, Oxidoreductase | Enterobacter mori |
| V3QX60 | 64.95 | 0 | _ | UDP-glucose 6-dehydrogenase | 1.1.1.22 | NAD, Oxidoreductase | Enterobacter sp. MGH 24 |
| A0A0D7LW76 | 64.69 | 0 | _ | UDP-glucose 6-dehydrogenase | 1.1.1.22 | NAD, Oxidoreductase | Citrobacter freundii |
| A0A0M7DFE8 | 64.69 | 0 | ugd | UDP-glucose 6-dehydrogenase | 1.1.1.22 | NAD, Oxidoreductase | Enterobacter cloacae |
| A0A0H3CLX6 | 64.69 | 0 | _ | UDP-glucose 6-dehydrogenase | 1.1.1.22 | NAD, Oxidoreductase | Enterobacter cloacae subsp. cloacae (strain ATCC 13047 / DSM 30054 / NBRC 13535 / NCDC 279-56) |

| | | | | | | | |
|---|---|---|---|---|---|---|---|
| A0A0J9WZA6 | 64.18 | 9,00E-178 | ugd | UDP-glucose 6-dehydrogenase | 1.1.1.22 | 3D-structure, NAD, Nucleotide-binding, Oxidoreductase | Klebsiella pneumoniae subsp. pneumoniae NTUH-K2044 |
| O06519 | 63.92 | 5,00E-177 | ugd | UDP-glucose 6-dehydrogenase | 1.1.1.22 | NAD, Oxidoreductase | Escherichia coli |
| Q6U8B9 | 63.66 | 5,00E-177 | ugd | UDP-glucose 6-dehydrogenase | 1.1.1.22 | NAD, Oxidoreductase | Raoultella terrigena |
| Q56625 | 62.11 | 6,00E-176 | _ | UDP-glucose 6-dehydrogenase | 1.1.1.22 | NAD, Oxidoreductase | Vibrio cholerae O139 |
| Q9RP54 | 63.14 | 1,00E-174 | ugd | UDP-glucose 6-dehydrogenase | 1.1.1.22 | NAD, Oxidoreductase | Escherichia coli |
| M9P0X4 | 60.31 | 1,00E-169 | ugd | UDP-glucose 6-dehydrogenase | 1.1.1.22 | NAD, Oxidoreductase | Providencia alcalifaciens |

**Genomic Object Editor: llo3156**

| PB id | Ident % | Eval | Gene | Description | EC number | Keywords | Organism |
|---|---|---|---|---|---|---|---|
| D3HMC0 | 100 | 0 | _ | Putative glycosyl transferase group 1 | _ | Reference proteome, Transferase | Legionella longbeachae serogroup 1 (strain NSW150) |
| H2FUE7 | 47.1 | 0 | _ | Glycosyl transferase group 1 | _ | Reference proteome, Transferase | Oceanimonas sp. (strain GK1) |
| A4BRU1 | 46.78 | 0 | _ | Glycosyltransferase | _ | Reference proteome, Transferase | Nitrococcus mobilis Nb-231 |
| A0A1I3JCB7 | 36.83 | 2,00E-114 | _ | Glycosyl transferases group 1 | _ | Transferase | Nitrosomonas sp. Nm34 |
| A0A0P9U3K9 | 36.88 | 6,00E-113 | _ | Glycos_transf_1 domain-containing protein | _ | _ | Pseudomonas syringae pv. helianthi |
| A0A101D0Z2 | 35.6 | 1,00E-110 | _ | Glycos_transf_1 domain-containing protein | _ | _ | Halomonas sp. 54_146 |
| A0A1H2QA17 | 35.97 | 1,00E-107 | _ | Glycosyltransferase involved in cell wall bisynthesis | _ | Transferase | Nitrosomonas communis |
| A0A1I4MRS6 | 36.35 | 1,00E-106 | _ | Glycosyl transferases group 1 | _ | Transferase | Nitrosomonas communis |
| K9E0G4 | 35.75 | 2,00E-105 | _ | Glycos_transf_1 domain-containing protein | _ | Reference proteome | Massilia timonae CCUG 45783 |
| Q9RQU9 | 29.38 | 1,00E-31 | wbqA | Putative perosamine transferase | _ | Transferase | Caulobacter vibrioides |
| L7S459 | 34.65 | 0.00000004 | gwEuk | Glycose transferase group 1 domain protein | _ | Transferase | Phytophthora hibernalis |
| M4QPQ6 | 28.67 | 0.00000006 | wbdA | Mannosyltransferase | _ | Glycosyltransferase, Transferase | Escherichia coli |
| L7S4R0 | 34.65 | 0.0000001 | gwEuk | Glycose transferase group 1 domain protein | _ | Transferase | Phytophthora ramorum |
| L7SZH1 | 33.66 | 0.0000003 | gwEuk 30.30.1 | Glycosyltransferase group 1 domain | _ | Transferase | Phytophthora lateralis |
| Q9RMU0 | 23.38 | 0.000002 | wbdA | WbdA | _ | _ | Klebsiella pneumoniae |

**Genomic Object Editor: llo3157**

| PB id | Ident % | Eval | Gene | Description | EC number | Keywords | Organism |
|---|---|---|---|---|---|---|---|
| D3HMC1 | 100 | 0 | _ | Uncharacterized protein | _ | Coiled coil, Reference proteome | Legionella longbeachae serogroup 1 (strain NSW150) |
| A0A2S4KG72 | 25.84 | 1,00E-33 | _ | Uncharacterized protein | _ | Coiled coil | Diaphorobacter sp. LR2014-1 |
| A0A2D4Y0U4 | 22.8 | 3,00E-24 | _ | Uncharacterized protein | _ | Coiled coil | Sphingomonadaceae bacterium |
| A0A011NI14 | 28.01 | 1,00E-21 | _ | Uncharacterized protein | _ | Coiled coil | Candidatus Accumulibacter sp. BA-92 |
| A0A2T7UAV0 | 23.98 | 2,00E-21 | _ | Uncharacterized protein | _ | Coiled coil | Limnohabitans planktonicus II-D5 |
| A0A1H8X4C5 | 34.33 | 1,00E-17 | _ | Uncharacterized protein | _ | _ | Pseudomonas sp. Snoq117.2 |
| A0A0K8NZ38 | 26.47 | 3,00E-17 | _ | Uncharacterized protein | _ | Coiled coil, Reference proteome | Ideonella sakaiensis (strain NBRC 110686 / TISTR 2288 / 201-F6) |
| D0QYN0 | 23.73 | 0.000000000001 | ccbE | CcbE | _ | Coiled coil | Avibacterium paragallinarum |
| D0QYN6 | 22.59 | 0.000000000003 | ccbE | CcbE | _ | Coiled coil | Avibacterium paragallinarum |
| Q9DUN0 | 31.97 | 0.00000001 | _ | Orf73 | _ | 3D-structure | Human herpesvirus 8 |
| Q91LX9 | 31.97 | 0.00000001 | _ | ORF73 | _ | _ | Human herpesvirus 8 |
| W5U981 | 24.81 | 0.00000001 | _ | Htt | _ | _ | Homo sapiens |
| Q76SB0 | 29.41 | 0.00000002 | _ | ORF 73 | _ | 3D-structure | Human herpesvirus 8 type M |
| Q98148 | 29.41 | 0.00000002 | _ | Kaposi's sarcoma-associated herpes-like virus ORF73 homolog | _ | 3D-structure | Human herpesvirus 8 |
| E5LC01 | 31.15 | 0.00000002 | ORF73 | LANA | _ | _ | Human herpesvirus 8 |
| Q9DUM3 | 31.97 | 0.00000002 | _ | Latent nuclear antigen | _ | 3D-structure | Human herpesvirus 8 |
| Q2KPA5 | 27.05 | 0.00000004 | _ | Clock | _ | Coiled coil, DNA-binding, Repeat | Macrobrachium rosenbergii |
| D0QYL8 | 22.3 | 0.0000001 | acbE | AcbE | _ | Coiled coil | Avibacterium paragallinarum |
| B5SUM8 | 23.18 | 0.0000004 | MED15 | Mediator of RNA polymerase II transcription subunit 15 | _ | Activator, Coiled coil, Nucleus, Transcription, Transcription regulation | Hyla arborea |

**Genomic Object Editor: llo3158**

| PB id | Ident % | Eval | Gene | Description | EC number | Keywords | Organism |
|---|---|---|---|---|---|---|---|
| D3HMC2 | 100 | 0 | _ | Uncharacterized protein | _ | Coiled coil, Reference proteome | Legionella longbeachae serogroup 1 (strain NSW150) |
| A0A1R4GY81 | 23.72 | 9,00E-54 | _ | Uncharacterized protein | _ | Coiled coil, Reference proteome | Crenothrix polyspora |
| A0A1Y1IYW1 | 25.57 | 4,00E-50 | _ | Signal recognition particle receptor protein FtsY | _ | Coiled coil, Receptor | Comamonas testosteroni |
| A0A1I4MS07 | 24.75 | 1,00E-49 | _ | Uncharacterized protein | _ | Coiled coil | Nitrosomonas communis |
| A0A096GZV8 | 23.08 | 3,00E-45 | _ | Methyltransf_21 domain-containing protein | _ | Coiled coil | Comamonas testosteroni |
| A0A1I3JE51 | 24.08 | 4,00E-41 | _ | Uncharacterized protein | _ | Coiled coil | Nitrosomonas sp. Nm34 |
| D6Z4R0 | 24.66 | 6,00E-41 | _ | Chromosome segregation ATPase-like protein | _ | Coiled coil, Reference proteome | Desulfurivibrio alkaliphilus (strain DSM 19089 / UNIQEM U267 / AHT2) |
| A0A0W7Z290 | 25.41 | 8,00E-41 | _ | Uncharacterized protein | _ | Coiled coil, Reference proteome | Comamonas kerstersii |
| A0A0F7KD18 | 25.04 | 1,00E-39 | _ | Uncharacterized protein | _ | Coiled coil, Reference proteome | Nitrosomonas communis |
| A0A1F9IRE7 | 26.98 | 2,00E-38 | _ | Uncharacterized protein | _ | Coiled coil | Deltaproteobacteria bacterium RIFCSPLOWO2_02_FULL_53_8 |
| Q6X1Y7 | 22.33 | 4,00E-24 | lepB | Effector protein B | _ | Coiled coil, Transmembrane | Legionella pneumophila |
| Q5ZSM7 | 22.33 | 4,00E-24 | lepB | LepB | _ | Coiled coil, Transmembrane | Legionella pneumophila subsp. pneumophila (strain Philadelphia 1 / ATCC 33152 / DSM 7513) |
| Q7K5Q6 | 26.48 | 4,00E-23 | maebl | Erythrocyte binding protein 3 | _ | Coiled coil, Signal | Plasmodium falciparum |
| Q8T5C7 | 26.48 | 4,00E-23 | maebl | Chimeric erythrocyte-binding protein MAEBL | _ | Coiled coil, Signal, Transmembrane | Plasmodium falciparum |
| Q7K5Q5 | 26.48 | 4,00E-23 | maebl | Erythrocyte binding protein 2 | _ | Coiled coil, Signal | Plasmodium falciparum |
| E5LC01 | 20.78 | 1,00E-22 | ORF73 | LANA | _ | _ | Human herpesvirus 8 |
| Q6A178 | 26.8 | 2,00E-22 | mt1 | Myosin tail 1 protein | _ | Coiled coil | Cryptosporidium parvum |
| Q91LX9 | 21.05 | 3,00E-22 | _ | ORF73 | _ | _ | Human herpesvirus 8 |
| Q25893 | 26.88 | 5,00E-22 | LSA-1 | Liver stage antigen | _ | Membrane, Transmembrane, Transmembrane helix | Plasmodium falciparum |

| | | | | | | | |
|---|---|---|---|---|---|---|---|
| O44934 | 23.21 | 9,00E-22 | _ | Myosin heavy chain isoform A | _ | Actin-binding, ATP-binding, Coiled coil, Myosin, Nucleotide-binding | Doryteuthis pealeii |
| G1EIL6 | 20.49 | 1,00E-21 | tnks1bp1 | Tankyrase 1 binding protein 1 | _ | _ | Danio rerio |
| Q9U0S6 | 23.82 | 1,00E-20 | prm MHC | Pedal retractor muscle myosin heavy chain | _ | Actin-binding, ATP-binding, Coiled coil, Myosin, Nucleotide-binding | Mytilus galloprovincialis |
| A0A140UGH3 | 23.68 | 1,00E-20 | _ | Myosin 2 heavy chain striated muscle | _ | Actin-binding, ATP-binding, Coiled coil, Myosin, Nucleotide-binding | Aphonopelma |

**Genomic Object Editor: llo3159**

| PB id | Ident % | Eval | Gene | Description | EC number | Keywords | Organism |
|---|---|---|---|---|---|---|---|
| D3HMC3 | 100 | 0 | _ | Uncharacterized protein | _ | Reference proteome | Legionella longbeachae serogroup 1 (strain NSW150) |
| A4BRT8 | 41.15 | 5,00E-131 | _ | Uncharacterized protein | _ | Reference proteome | Nitrococcus mobilis Nb-231 |
| A0A1H3CQ84 | 58.71 | 1,00E-69 | _ | Uncharacterized protein | _ | Reference proteome | Roseicitreum antarcticum |
| A0A1X7A7H2 | 42.42 | 6,00E-66 | _ | Uncharacterized protein | _ | Coiled coil, Reference proteome | Limimaricola soesokkakensis |
| A0A1Q4CSG6 | 55.22 | 1,00E-64 | _ | Uncharacterized protein | _ | _ | Rhodobacterales bacterium 65-51 |
| A0A2E9GVE9 | 42.72 | 4,00E-48 | _ | Uncharacterized protein | _ | _ | Deltaproteobacteria bacterium |
| A0A2E6LM58 | 35.68 | 2,00E-30 | _ | Methyltransf_21 domain-containing protein | _ | Coiled coil | Gammaproteobacteria bacterium |
| A0A0F7KKU9 | 35.58 | 2,00E-29 | _ | Methyltransf_21 domain-containing protein | _ | Coiled coil, Reference proteome | Nitrosomonas communis |
| A0A1I3JDG0 | 35.58 | 2,00E-29 | _ | Methyltransferase, FkbM family | _ | Coiled coil, Methyltransferase, Transferase | Nitrosomonas sp. Nm34 |
| A0A1H2Q970 | 42.41 | 2,00E-27 | _ | Methyltransferase, FkbM family | _ | Methyltransferase, Transferase | Nitrosomonas communis |

**Genomic Object Editor: llo3160**

| PB id | Ident % | Eval | Gene | Description | EC number | Keywords | Organism |
|---|---|---|---|---|---|---|---|
| D3HMC4 | 100 | 4,00E-170 | _ | Putative lipopolysaccharide core biosynthesis protein | _ | Reference proteome | Legionella longbeachae serogroup 1 (strain NSW150) |
| A0A0P7WRL6 | 64.44 | 4,00E-105 | _ | Uncharacterized protein | _ | _ | Idiomarinaceae bacterium HL-53 |
| A0A1Q4CSD3 | 53.78 | 2,00E-81 | _ | Uncharacterized protein | _ | _ | Rhodobacterales bacterium 65-51 |
| A4BRT7 | 58.29 | 6,00E-81 | _ | Uncharacterized protein | _ | Reference proteome | Nitrococcus mobilis Nb-231 |
| A0A1H9L2A0 | 52.73 | 1,00E-79 | _ | Uncharacterized protein | _ | Reference proteome | Litorimicrobium taeanense |
| A0A254QKC4 | 52.65 | 2,00E-79 | _ | Uncharacterized protein | _ | Reference proteome | Phaeobacter sp. 22II1-1F12B |
| A0A1V0RJS1 | 54.13 | 5,00E-78 | _ | Uncharacterized protein | _ | _ | Roseovarius mucosus |
| K1XV28 | 52.27 | 6,00E-73 | _ | Uncharacterized protein | _ | _ | uncultured bacterium |
| A0YTV4 | 51.83 | 1,00E-72 | _ | TPR_REGION domain-containing protein | _ | Coiled coil, Reference proteome, TPR repeat | Lyngbya sp. (strain PCC 8106) |
| Q9XC98 | 26.44 | 0.003 | _ | Lipopolysaccharide core biosynthesis protein RfaZ | _ | Membrane, Transferase, Transmembrane, Transmembrane helix | Klebsiella pneumoniae |
| I7AU32 | 27.78 | 0.62 | waaZ | 3-deoxy-D-manno-oct-2-ulosonate III transferase WaaZ | 2.4.99.15 | Glycosyltransferase, Transferase | Escherichia coli |

**Genomic Object Editor: llo3161**

| PB id | Ident % | Eval | Gene | Description | EC number | Keywords | Organism |
|---|---|---|---|---|---|---|---|
| D3HMC5 | 100 | 0 | _ | Putative glycosyl transferase family 2 | _ | Reference proteome, Transferase | Legionella longbeachae serogroup 1 (strain NSW150) |
| Q07Z86 | 59.24 | 9,00E-140 | _ | Glycosyl transferase, family 2 | _ | Reference proteome, Transferase | Shewanella frigidimarina (strain NCIMB 400) |
| A0A0P8B481 | 60.88 | 1,00E-130 | _ | Family 2 glycosyltransferase | _ | Transferase | Idiomarinaceae bacterium HL-53 |
| A0A0F7M0S2 | 54.78 | 2,00E-122 | _ | Glycosyl transferase | _ | Reference proteome, Transferase | Spongiibacter sp. IMCC21906 |
| A0A1B7WX93 | 57 | 3,00E-119 | _ | Glycosyl transferase family 2 | _ | Transferase | Anabaena sp. MDT14b |
| A0A1G1H1U7 | 58.36 | 2,00E-117 | _ | Glycosyl transferase family 2 | _ | Transferase | Nitrospirae bacterium GWC2_57_9 |
| A0A1X7A7Q4 | 51.1 | 7,00E-113 | _ | N-glycosyltransferase | _ | Reference proteome, Transferase | Limimaricola soesokkakensis |
| A0A090SRU4 | 54.79 | 3,00E-112 | _ | Glyco_trans_2-like domain-containing protein | _ | _ | Vibrio maritimus |
| A0A1W9GAI3 | 54.61 | 1,00E-110 | _ | Glycosyl transferase family 2 | _ | Transferase | Nitrospira sp. SG-bin2 |
| A0A1V0RJU4 | 50.47 | 3,00E-109 | _ | Putative glycosyl transferase | _ | Transferase | Roseovarius mucosus |
| Q3ZK45 | 41.07 | 2,00E-20 | epsG | EpsG | _ | _ | Lactococcus lactis |
| O66259 | 30.32 | 0.00000000000002 | _ | Glycosyltransferase | _ | Transferase | Aggregatibacter actinomycetemcomitans |
| Q9XDQ0 | 30.67 | 0.0000000000001 | ORF14 | Putative glycosyltransferase | _ | Transferase | Aggregatibacter actinomycetemcomitans |
| Q54129 | 31.22 | 0.00000000002 | wbaN | Rhamnosyl transferase | _ | Transferase | Salmonella enterica |
| Q9AQA9 | 30 | 0.00000000005 | _ | Putative rhamnosyltransferase | _ | Transferase | Aggregatibacter actinomycetemcomitans |
| K4P2X6 | 33.33 | 0.0000003 | wbyL | WbyL | _ | _ | Yersinia similis |
| Q8GNC0 | 30.56 | 0.000002 | lgtA | N-acetylglucosamine glycosyltransferase | _ | Transferase | Haemophilus ducreyi |

**Genomic Object Editor: llo3162**

| PB id | Ident % | Eval | Gene | Description | EC number | Keywords | Organism |
|---|---|---|---|---|---|---|---|
| D3HMC6 | 100 | 3,00E-161 | _ | Putative acylneuraminate cytidylyltransferase | _ | Nucleotidyltransferase, Reference proteome, Transferase | Legionella longbeachae serogroup 1 (strain NSW150) |
| K2JYG6 | 70.45 | 2,00E-106 | _ | Acylneuraminate cytidylyltransferase | _ | Nucleotidyltransferase, Reference proteome, Transferase | Gallaecimonas xiamenensis 3-C-1 |
| A0A0F7M0J7 | 68.66 | 6,00E-104 | _ | CMP-N-acetylneuraminic acid synthetase | _ | Reference proteome | Spongiibacter sp. IMCC21906 |
| A0A0M1JD43 | 68.42 | 8,00E-101 | _ | Acylneuraminate cytidylyltransferase | _ | Nucleotidyltransferase, Reference proteome, Transferase | Achromatium sp. WMS3 |
| A0A0M1J3Y3 | 67.94 | 1,00E-99 | _ | Acylneuraminate cytidylyltransferase | _ | Nucleotidyltransferase, Reference proteome, Transferase | Achromatium sp. WMS3 |
| A0A011NPN0 | 65.9 | 1,00E-97 | _ | 3-deoxy-manno-octulosonate cytidylyltransferase | _ | Nucleotidyltransferase, Reference proteome, Transferase | Candidatus Accumulibacter sp. SK-11 |
| A0A250KNV9 | 66.21 | 2,00E-97 | _ | Acylneuraminate cytidylyltransferase | _ | Nucleotidyltransferase, Reference proteome, Transferase | Methylocaldum marinum |

**Supplementary Table 1** List of the highest hits of the *L. longbeachae* capsule cluster genes against the NCBI database (Trembl).

| | | | | | | | |
|---|---|---|---|---|---|---|---|
| A0A1H9KWA5 | 63.27 | 2,00E-96 | _ | CMP-N-acetylneuraminic acid synthetase | _ | Reference proteome | Litorimicrobium taeanense |
| A0A0N8KBF2 | 66.51 | 4,00E-96 | neuA | N-acylneuraminate cytidylyltransferase NeuA | _ | Nucleotidyltransferase, Transferase | Idiomarinaceae bacterium HL-53 |
| A0A254QKC9 | 65.75 | 2,00E-95 | _ | Acylneuraminate cytidylyltransferase | _ | Nucleotidyltransferase, Reference proteome, Transferase | Phaeobacter sp. 22II1-1F12B |
| Q933W2 | 23.64 | 0.0000001 | neuA1 | Acylneuraminate cytidylyltransferase | _ | Nucleotidyltransferase, Transferase | Campylobacter jejuni |
| Q077S2 | 21.98 | 0.0000002 | nnaC | Acylneuraminate cytidylyltransferase | 2.7.7.43 | Nucleotidyltransferase, Transferase | Escherichia coli |

**Genomic Object Editor: llo3163**

| PB id | Ident % | Eval | Gene | Description | EC number | Keywords | Organism |
|---|---|---|---|---|---|---|---|
| D3HMC7 | 100 | 0 | _ | Putative D-isomer specific 2-hydroxyacid dehydrogenase | _ | NAD, Oxidoreductase, Reference proteome | Legionella longbeachae serogroup 1 (strain NSW150) |
| B8CL20 | 68.06 | 8,00E-152 | _ | D-isomer specific 2-hydroxyacid dehydrogenase, catalytic region, D-isomer specific 2-hydroxyacid dehydrogenase, NAD-binding | _ | NAD, Oxidoreductase | Shewanella piezotolerans (strain WP3 / JCM 13877) |
| Q07Z88 | 67.2 | 1,00E-150 | _ | D-isomer specific 2-hydroxyacid dehydrogenase, NAD-binding | _ | NAD, Oxidoreductase, Reference proteome | Shewanella frigidimarina (strain NCIMB 400) |
| A0A0P7ZKZ6 | 66.56 | 8,00E-147 | serA | D-3-phosphoglycerate dehydrogenase | 1.1.1.95 | NAD, Oxidoreductase | Idiomarinaceae bacterium HL-53 |
| A0A0C3MQY8 | 65.05 | 2,00E-143 | _ | Phosphoglycerate dehydrogenase | _ | NAD, Oxidoreductase | Shewanella sp. cp20 |
| A4BRT3 | 60.91 | 3,00E-133 | _ | Phosphoglycerate dehydrogenase | _ | NAD, Oxidoreductase, Reference proteome | Nitrococcus mobilis Nb-231 |
| A0A254QKD0 | 60.33 | 3,00E-126 | _ | Phosphoglycerate dehydrogenase | _ | Oxidoreductase, Reference proteome | Phaeobacter sp. 22II1-1F12B |
| A0A0P1EMI7 | 58.55 | 1,00E-124 | serA_2 | D-3-phosphoglycerate dehydrogenase | 1.1.1.95 | Oxidoreductase, Reference proteome | Shimia marina |
| A0A1Q4CSG3 | 59.22 | 3,00E-124 | _ | Phosphoglycerate dehydrogenase | _ | Oxidoreductase | Rhodobacterales bacterium 65-51 |
| A0A1X7A7P1 | 58.17 | 1,00E-123 | tkrA_2 | Glyoxylate/hydroxypyruvate reductase B | 1.1.1.79 | Oxidoreductase, Pyruvate, Reference proteome | Limimaricola soesokkakensis |
| F8AEA4 | 35.64 | 8,00E-37 | gyaR | Glyoxylate reductase | 1.1.1.26 | 3D-structure, Cytoplasm, NAD, Oxidoreductase | Pyrococcus yayanosii (strain CH1 / JCM 16557) |
| Q2TL63 | 32.23 | 2,00E-32 | _ | D-3-phosphoglycerate dehydrogenase | 1.1.1.95 | Amino-acid biosynthesis, NAD, Oxidoreductase, Serine biosynthesis | Mesorhizobium ciceri |
| A8R0N0 | 30.55 | 1,00E-31 | ApPGDH | D-3-phosphoglycerate dehydrogenase | 1.1.1.95 | Amino-acid biosynthesis, NAD, Oxidoreductase, Serine biosynthesis | Aphanothece halophytica |
| U5TVU1 | 32.22 | 4,00E-27 | mcyI | McyI | _ | Oxidoreductase | Nostoc sp. 152 |
| G4XDR8 | 32.37 | 2,00E-26 | ptxD | Phosphite dehydrogenase | _ | 3D-structure, Oxidoreductase | Ralstonia sp. 4506 |

**Genomic Object Editor: llo3164**

| PB id | Ident % | Eval | Gene | Description | EC number | Keywords | Organism |
|---|---|---|---|---|---|---|---|
| Q8KWT4 | 31.47 | 5,00E-25 | bacC | Dihydroanticapsin 7-dehydrogenase | 1.1.1.385 | Antibiotic biosynthesis, NAD, Oxidoreductase | Bacillus subtilis |
| Q9WYG0 | 32.51 | 1,00E-24 | _ | Uncharacterized oxidoreductase TM_0325 | 1.-.-.- | Oxidoreductase, Reference proteome | Thermotoga maritima (strain ATCC 43589 / MSB8 / DSM 3109 / JCM 10099) |
| P39640 | 30 | 2,00E-23 | bacC | Dihydroanticapsin 7-dehydrogenase | 1.1.1.385 | 3D-structure, Antibiotic biosynthesis, NAD, Oxidoreductase, Reference proteome | Bacillus subtilis (strain 168) |
| P50199 | 28.81 | 2,00E-23 | gno | Gluconate 5-dehydrogenase | 1.1.1.- | Carbohydrate metabolism, Cytoplasm, Direct protein sequencing, NADP, Oxidoreductase, Reference proteome | Gluconobacter oxydans (strain 621H) |
| Q56318 | 29.27 | 9,00E-23 | _ | Uncharacterized oxidoreductase TM_0019 | 1.-.-.- | NADP, Oxidoreductase, Reference proteome | Thermotoga maritima (strain ATCC 43589 / MSB8 / DSM 3109 / JCM 10099) |
| P40288 | 28.23 | 1,00E-21 | _ | Glucose 1-dehydrogenase | 1.1.1.47 | 3D-structure, Direct protein sequencing, NADP, Oxidoreductase, Sporulation | Bacillus megaterium |
| Q51576 | 29.64 | 1,00E-21 | _ | Uncharacterized oxidoreductase PA3106 | 1.-.-.- | NADP, Oxidoreductase, Reference proteome | Pseudomonas aeruginosa (strain ATCC 15692 / DSM 22644 / CIP 104116 / JCM 14847 / LMG 12228 / 1C / PRS 101 / PAO1) |
| P39482 | 27.35 | 1,00E-21 | gdhI | Glucose 1-dehydrogenase 1 | 1.1.1.47 | Germination, NADP, Oxidoreductase, Sporulation | Bacillus megaterium |
| P46331 | 30.08 | 2,00E-21 | yxbG | Uncharacterized oxidoreductase YxbG | 1.-.-.- | NAD, Oxidoreductase, Reference proteome | Bacillus subtilis (strain 168) |
| Q92RN6 | 34.74 | 4,00E-21 | galD | Probable galactose dehydrogenase GalD | 1.1.1.- | NADP, Oxidoreductase, Reference proteome | Rhizobium meliloti (strain 1021) |
| P08074 | 30.36 | 6,00E-21 | Cbr2 | Carbonyl reductase | NADPH] 2 | 1.1.1.184 | 2455724, 7705352, 15489334, 8040004, 8999926, 21183079, 8805511 |
| P05406 | 31.64 | 8,00E-21 | fixR | Protein FixR | _ | Nitrogen fixation, Oxidoreductase, Reference proteome | Bradyrhizobium diazoefficiens (strain JCM 10833 / BCRC 13528 / IAM 13628 / NBRC 14792 / USDA 110) |
| Q53882 | 28.96 | 8,00E-21 | dauE | Aklaviketone reductase DauE | 1.1.1.362 | Antibiotic biosynthesis, NADP, Oxidoreductase | Streptomyces sp. (strain C5) |

**Genomic Object Editor: llo3165**

| PB id | Ident % | Eval | Gene | Description | EC number | Keywords | Organism |
|---|---|---|---|---|---|---|---|
| D3HMC9 | 100 | 0 | _ | Putative glycosyl transferase family 2 | _ | Reference proteome, Transferase | Legionella longbeachae serogroup 1 (strain NSW150) |
| A4BRT1 | 62.34 | 0 | _ | Uncharacterized protein | _ | Reference proteome | Nitrococcus mobilis Nb-231 |
| A0A1X7A785 | 58.81 | 0 | _ | Uncharacterized protein | _ | Coiled coil, Reference proteome | Limimaricola soesokkakensis |
| A0A239Q0E7 | 54.37 | 0 | _ | Methyltransferase domain-containing protein | _ | Coiled coil, Membrane, Methyltransferase, Reference proteome, Transferase, Transmembrane, Transmembrane helix | Amphiplicatus metriothermophilus |
| A0A2E7LM08 | 49.77 | 0 | _ | Glycosyl transferase family 2 | _ | Transferase | Dehalococcoidia bacterium |
| A0A1Q4CSM0 | 47.31 | 0 | _ | Uncharacterized protein | _ | _ | Rhodobacterales bacterium 65-51 |

**Supplementary Table 1** List of the highest hits of the *L. longbeachae* capsule cluster genes against the NCBI database (Trembl).

| | | | | | | | |
|---|---|---|---|---|---|---|---|
| A0A2M7GNM0 | 70.08 | 8,00E-129 | _ | Uncharacterized protein | _ | Coiled coil, Membrane, Transmembrane, Transmembrane helix | Rhodobacterales bacterium CG15_BIG_FIL_POST_REV_8_21_14_020_59_13 |
| K2KDH9 | 58.53 | 4,00E-99 | _ | Uncharacterized protein | _ | Reference proteome | Gallaecimonas xiamenensis 3-C-1 |
| B8CL24 | 59.92 | 1,00E-96 | _ | Uncharacterized protein | _ | _ | Shewanella piezotolerans (strain WP3 / JCM 13877) |

**Genomic Object Editor: llo3166**

| PB id | Ident % | Eval | Gene | Description | EC number | Keywords | Organism |
|---|---|---|---|---|---|---|---|
| D3HMD0 | 100 | 0 | galE | UDP-glucose 4-epimerase | 5.1.3.2 | Carbohydrate metabolism, Isomerase, NAD, Reference proteome | Legionella longbeachae serogroup 1 (strain NSW150) |
| D3HMD6 | 63.44 | 1,00E-160 | galE | UDP-glucose 4-epimerase | 5.1.3.2 | Carbohydrate metabolism, Isomerase, NAD, Reference proteome | Legionella longbeachae serogroup 1 (strain NSW150) |
| R4YS92 | 56.36 | 1,00E-131 | galE | UDP-glucose 4-epimerase | 5.1.3.2 | Carbohydrate metabolism, Isomerase, NAD, Reference proteome | Oleispira antarctica RB-8 |
| A0A2C4RG95 | 55.49 | 6,00E-129 | galE | UDP-glucose 4-epimerase | 5.1.3.2 | Carbohydrate metabolism, Isomerase, NAD | Bacillus sp. AFS043905 |
| A0A1H9KXM1 | 56.36 | 6,00E-127 | _ | UDP-glucose 4-epimerase | 5.1.3.2 | Carbohydrate metabolism, Isomerase, NAD | Butyrivibrio fibrisolvens |
| A0A1M6BZ40 | 56.06 | 1,00E-126 | _ | UDP-glucose 4-epimerase | 5.1.3.2 | Carbohydrate metabolism, Isomerase, NAD | Butyrivibrio fibrisolvens DSM 3071 |
| A0A1H9ELV8 | 55.76 | 5,00E-126 | _ | UDP-glucose 4-epimerase | 5.1.3.2 | Carbohydrate metabolism, Isomerase, NAD, Reference proteome | Butyrivibrio sp. TB |
| A0A098L0R3 | 55.79 | 1,00E-124 | _ | UDP-glucose 4-epimerase | 5.1.3.2 | Carbohydrate metabolism, Coiled coil, Isomerase, NAD | Geobacillus thermoleovorans B23 |
| A0A1G8VHA8 | 54.55 | 1,00E-124 | _ | UDP-glucose 4-epimerase | 5.1.3.2 | Carbohydrate metabolism, Coiled coil, Isomerase, NAD, Reference proteome | Lachnospiraceae bacterium G41 |
| M5WMM1 | 53.5 | 1,00E-114 | UGE | UDP-glucose 4-epimerase | 5.1.3.- | Carbohydrate metabolism, Isomerase, NAD, Reference proteome | Prunus persica |
| Q58IJ6 | 53.33 | 3,00E-113 | UGE1 | UDP-glucose 4-epimerase | 5.1.3.- | Carbohydrate metabolism, Coiled coil, Isomerase, NAD | Hordeum vulgare |
| Q9RP56 | 50.3 | 7,00E-113 | galE | UDP-glucose 4-epimerase | 5.1.3.2 | Carbohydrate metabolism, Isomerase, NAD | Escherichia coli |
| Q58IJ5 | 50.45 | 4,00E-110 | UGE2 | UDP-glucose 4-epimerase | 5.1.3.- | Carbohydrate metabolism, Isomerase, NAD | Hordeum vulgare |
| A0A2D0W0L9 | 49.24 | 1,00E-109 | gne | UDP-glucose 4-epimerase | 5.1.3.2 | Carbohydrate metabolism, Isomerase, NAD | Escherichia fergusonii |
| Q2QD27 | 50 | 2,00E-109 | gne | UDP-glucose 4-epimerase | 5.1.3.2 | Carbohydrate metabolism, Isomerase, NAD | Aeromonas hydrophila |
| M9VRS7 | 48.77 | 2,00E-109 | galE2 | UDP-glucose 4-epimerase | 5.1.3.2 | Carbohydrate metabolism, Isomerase, NAD | Streptococcus oralis |

**Genomic Object Editor: llo3167**

| PB id | Ident % | Eval | Gene | Description | EC number | Keywords | Organism |
|---|---|---|---|---|---|---|---|
| D3HMD1 | 100 | 0 | gmd | GDP-mannose 4,6-dehydratase | 4.2.1.47 | Lyase, NADP, Reference proteome | Legionella longbeachae serogroup 1 (strain NSW150) |
| A0A024HQL7 | 76.09 | 0 | bre-1 | GDP-mannose 4,6-dehydratase | 4.2.1.47 | Lyase, NADP, Reference proteome | Pseudomonas knackmussii (strain DSM 6978 / LMG 23759 / B13) |
| A0A0A7EDA1 | 75.66 | 0 | gmd | GDP-mannose 4,6-dehydratase | 4.2.1.47 | Lyase, NADP, Reference proteome | Pseudoalteromonas piratica |
| A0A0Q5FPZ0 | 75.22 | 0 | gmd | GDP-mannose 4,6-dehydratase | 4.2.1.47 | Lyase, NADP | Pseudomonas sp. Leaf127 |
| V9UXM5 | 74.34 | 0 | gmd | GDP-mannose 4,6-dehydratase | 4.2.1.47 | Lyase, NADP | Pseudomonas monteilii SB3101 |
| A0A177YY40 | 74.93 | 0 | gmd_1 | GDP-mannose 4,6-dehydratase | 4.2.1.47 | Lyase, NADP | Pseudomonas putida |
| A0A2J8I1D5 | 75.66 | 0 | gmd | GDP-mannose 4,6-dehydratase | 4.2.1.47 | Lyase, NADP | Vibrio diazotrophicus |
| A0A2S5IDD8 | 74.93 | 0 | gmd | GDP-mannose 4,6-dehydratase | 4.2.1.47 | Lyase, NADP | Pseudomonas aeruginosa |
| R1IXR6 | 75.37 | 0 | gmd | GDP-mannose 4,6-dehydratase | 4.2.1.47 | Lyase, NADP | Grimontia indica |
| A0A2V4KC88 | 74.93 | 0 | gmd | GDP-mannose 4,6-dehydratase | 4.2.1.47 | Lyase, NADP | Pseudomonas sp. MB-090624 |
| A0A147GBA3 | 74.64 | 0 | gmd | GDP-mannose 4,6-dehydratase | 4.2.1.47 | Lyase, NADP | Pseudomonas parafulva |
| F3KBD8 | 75.07 | 0 | gmd | GDP-mannose 4,6-dehydratase | 4.2.1.47 | Lyase, NADP, Reference proteome | gamma proteobacterium IMCC2047 |
| C8YZ33 | 74.49 | 0 | gmd | GDP-mannose 4,6-dehydratase | 4.2.1.47 | Lyase, NADP | Escherichia coli |
| D2KWB3 | 73.47 | 0 | gmd | GDP-mannose 4,6-dehydratase | 4.2.1.47 | Lyase, NADP | Pseudomonas savastanoi pv. glycinea |
| M4HNX2 | 71.51 | 6,00E-179 | bceN | GDP-mannose 4,6-dehydratase | 4.2.1.47 | Lyase, NADP | Burkholderia cepacia |
| K4NNR8 | 53.26 | 8,00E-121 | gmd | GDP-mannose 4,6-dehydratase | 4.2.1.47 | Lyase, NADP | Yersinia similis |
| Q9R966 | 51.63 | 4,00E-120 | gmd | GDP-mannose 4,6-dehydratase | 4.2.1.47 | Lyase, NADP | Brucella melitensis |
| O85352 | 53.1 | 4,00E-120 | gmd | GDP-mannose 4,6-dehydratase | 4.2.1.47 | Lyase, NADP | Caulobacter vibrioides |
| Q5ND85 | 52.26 | 8,00E-116 | gmd | GDP-mannose 4,6-dehydratase | 4.2.1.47 | Lyase, NADP | Yersinia sp. A125 KOH2 |
| A0A1B1R128 | 52.68 | 9,00E-116 | GMD2 | GDP-mannose 4,6-dehydratase | 4.2.1.47 | Lyase | Mortierella alpina |
| F1CLL6 | 51.15 | 1,00E-115 | gmd | GDP-mannose 4,6-dehydratase | 4.2.1.47 | Lyase, NADP | Yersinia pseudotuberculosis |
| Q6T1X7 | 53.89 | 1,00E-115 | gmd | GDP-mannose 4,6-dehydratase | 4.2.1.47 | Lyase, NADP | Aneurinibacillus thermoaerophilus |

**Genomic Object Editor: llo3168**

| PB id | Ident % | Eval | Gene | Description | EC number | Keywords | Organism |
|---|---|---|---|---|---|---|---|
| D3HMD2 | 100 | 0 | _ | Putative capsular polysaccharide biosynthesis protein | _ | Coiled coil, Reference proteome | Legionella longbeachae serogroup 1 (strain NSW150) |
| A0A1Q4CWN3 | 47.39 | 6,00E-88 | _ | WcbI domain-containing protein | _ | Coiled coil | Rhodobacterales bacterium 65-51 |
| A0A2H4UTY7 | 28.2 | 5,00E-24 | _ | WcbI domain-containing protein | _ | Reference proteome | Bodo saltans virus |
| A0A2N1DBJ2 | 26.25 | 0.0000000007 | _ | WcbI domain-containing protein | _ | Reference proteome | Paraglaciecola sp. MB-3u-78 |
| U7Q8X3 | 22.59 | 0.000000007 | _ | Nucleotide-diphospho-sugar transferase family protein | _ | Transferase | Lyngbya aestuarii BL J |
| A0A1M7TIW0 | 28.49 | 0.0000009 | _ | WcbI domain-containing protein | _ | Reference proteome | Desulfovibrio litoralis DSM 11393 |
| Q93UJ4 | 25.22 | 0.0001 | wcbI | WcbI | _ | _ | Burkholderia pseudomallei |
| Q63R74 | 24.78 | 0.0003 | wcbI | Putative capsular polysaccharide biosynthesis protein | _ | 3D-structure, Reference proteome | Burkholderia pseudomallei (strain K96243) |

**Genomic Object Editor: llo3169**

| PB id | Ident % | Eval | Gene | Description | EC number | Keywords | Organism |
|---|---|---|---|---|---|---|---|
| D3HMD3 | 99.03 | 1,00E-66 | _ | Uncharacterized protein | _ | Membrane, Reference protei | Legionella longbeachae serogroup 1 (strain NSW150) |
| NO MORE HITS! | | | | | | | |

**Genomic Object Editor: llo3170**

| PB id | Ident % | Eval | Gene | Description | EC number | Keywords | Organism |
|---|---|---|---|---|---|---|---|
| D3HMD4 | 100 | 0 | _ | SGL domain-containing protein | _ | Reference proteome | Legionella longbeachae serogroup 1 (strain NSW150) |

**Supplementary Table 1** List of the highest hits of the *L. longbeachae* capsule cluster genes against the NCBI database (Trembl).

| | | | | | | | |
|---|---|---|---|---|---|---|---|
| A0A1J8P5Y9 | 40.71 | 8,00E-59 | _ | Gluconolaconase | _ | _ | Candidatus Rickettsiella isopodorum |
| K2D0F4 | 38.67 | 4,00E-58 | _ | SMP-30/Gluconolaconase/LRE protein | _ | _ | uncultured bacterium |
| A0A1G0G7G2 | 38.67 | 4,00E-58 | _ | SGL domain-containing protein | _ | _ | Gammaproteobacteria bacterium RIFCSPHIGHO2_02_FULL_39_13 |
| A8PKQ3 | 38.43 | 9,00E-58 | _ | SMP-30/Gluconolaconase/LRE domain protein | _ | Reference proteome | Rickettsiella grylli |
| A0A1Y5DM83 | 34.91 | 9,00E-53 | _ | SGL domain-containing protein | _ | _ | Arcobacter sp. 31_11_sub10_T18 |
| A9KEW0 | 34.96 | 1,00E-52 | _ | Gluconolactonase | 3.1.1.17 | Hydrolase | Coxiella burnetii (strain Dugway 5J108-111) |
| A0A2E5LWQ5 | 40.32 | 2,00E-52 | _ | Gluconolaconase | _ | _ | Dehalococcoidia bacterium |
| A0A1G0GPR6 | 37.8 | 2,00E-52 | _ | SGL domain-containing protein | _ | _ | Gammaproteobacteria bacterium RIFCSPHIGHO2_12_FULL_36_30 |
| Q83AU0 | 34.59 | 2,00E-51 | _ | Gluconolactonase | 3.1.1.17 | Hydrolase, Reference proteome | Coxiella burnetii (strain RSA 493 / Nine Mile phase I) |
| A9CPS8 | 26.84 | 9,00E-19 | GNL | Lactonase | _ | _ | Euglena gracilis |
| E7BDE8 | 26.95 | 2,00E-17 | Dca | DCA protein | _ | _ | Drosophila guanche |
| Q8TA68 | 26.92 | 2,00E-17 | H-LRE | Luciferin-regenerating enzyme | _ | _ | Aquatica lateralis |
| Q86DU5 | 24.53 | 6,00E-17 | LRE | Luciferin-regenerating enzyme | _ | _ | Photinus pyralis |

**Genomic Object Editor: llo3171**

| PB id | Ident % | Eval | Gene | Description | EC number | Keywords | Organism |
|---|---|---|---|---|---|---|---|
| D3HMD5 | 100 | 0 | galU | UTP--glucose-1-phosphate uridylyltransferase | 2.7.7.9 | Nucleotidyltransferase, Reference proteome, Transferase | Legionella longbeachae serogroup 1 (strain NSW150) |
| A0A1J4QD25 | 66.43 | 7,00E-133 | _ | UTP--glucose-1-phosphate uridylyltransferase | 2.7.7.9 | Nucleotidyltransferase, Reference proteome, Transferase | Oceanisphaera psychrotolerans |
| K6YSZ7 | 62.85 | 1,00E-132 | galU | UTP--glucose-1-phosphate uridylyltransferase | 2.7.7.9 | Nucleotidyltransferase, Reference proteome, Transferase | Paraglaciecola arctica BSs20135 |
| A0A233RFF0 | 66.55 | 2,00E-132 | galU | UTP--glucose-1-phosphate uridylyltransferase | 2.7.7.9 | Nucleotidyltransferase, Reference proteome, Transferase | Oceanimonas doudoroffii |
| A0A135ZYV6 | 64.44 | 4,00E-132 | _ | UTP--glucose-1-phosphate uridylyltransferase | 2.7.7.9 | Nucleotidyltransferase, Reference proteome, Transferase | Paraglaciecola hydrolytica |
| R4YTW2 | 63.79 | 5,00E-132 | galU | UTP--glucose-1-phosphate uridylyltransferase | 2.7.7.9 | Nucleotidyltransferase, Reference proteome, Transferase | Oleispira antarctica RB-8 |
| A0A2S9VBQ5 | 65.23 | 6,00E-132 | galU | UTP--glucose-1-phosphate uridylyltransferase | 2.7.7.9 | Nucleotidyltransferase, Transferase | Alteromonas alba |
| A0A2D5LH48 | 65.23 | 6,00E-132 | galU | UTP--glucose-1-phosphate uridylyltransferase | 2.7.7.9 | Nucleotidyltransferase, Transferase | Alteromonas sp |
| A0A1E8FAQ7 | 65.34 | 9,00E-132 | _ | UTP--glucose-1-phosphate uridylyltransferase | 2.7.7.9 | Nucleotidyltransferase, Reference proteome, Transferase | Alteromonas lipolytica |
| A0A2D9S173 | 65.7 | 1,00E-131 | galU | UTP--glucose-1-phosphate uridylyltransferase | 2.7.7.9 | Nucleotidyltransferase, Transferase | Alteromonadaceae bacterium |
| E1SP06 | 63.7 | 2,00E-131 | _ | UTP--glucose-1-phosphate uridylyltransferase | 2.7.7.9 | Nucleotidyltransferase, Reference proteome, Transferase | Ferrimonas balearica (strain DSM 9799 / CCM 4581 / KCTC 23876 / PAT) |
| C7FFE7 | 65.47 | 3,00E-128 | _ | UTP--glucose-1-phosphate uridylyltransferase | 2.7.7.9 | Nucleotidyltransferase, Transferase | Proteus mirabilis |
| A7KAV1 | 65 | 4,00E-127 | _ | UTP--glucose-1-phosphate uridylyltransferase | 2.7.7.9 | Nucleotidyltransferase, Transferase | Aeromonas hydrophila |
| O85215 | 62.37 | 2,00E-126 | galU | UTP--glucose-1-phosphate uridylyltransferase | 2.7.7.9 | Nucleotidyltransferase, Transferase | Actinobacillus pleuropneumoniae |
| D4I3X5 | 63.7 | 6,00E-121 | galU | UTP--glucose-1-phosphate uridylyltransferase | 2.7.7.9 | 3D-structure, Nucleotidyltransferase, Transferase | Erwinia amylovora (strain CFBP1430) |
| Q70AL9 | 62.45 | 2,00E-120 | galU | UTP--glucose-1-phosphate uridylyltransferase | 2.7.7.9 | Nucleotidyltransferase, Transferase | Yersinia enterocolitica |
| Q84BL0 | 60.07 | 8,00E-120 | galU | UTP--glucose-1-phosphate uridylyltransferase | 2.7.7.9 | Nucleotidyltransferase, Transferase | Aeromonas hydrophila |
| Q848R8 | 59.04 | 1,00E-118 | galU | UTP--glucose-1-phosphate uridylyltransferase | 2.7.7.9 | Nucleotidyltransferase, Transferase | Aeromonas hydrophila |
| Q70AL8 | 59.29 | 2,00E-111 | galF | UTP--glucose-1-phosphate uridylyltransferase | 2.7.7.9 | Nucleotidyltransferase, Transferase | Yersinia enterocolitica |
| Q937X5 | 54.58 | 4,00E-97 | galF | Alpha-D-glucosyl-1-phosphate uridylyltransferase | 2.7.7.9 | Nucleotidyltransferase, Transferase | Edwardsiella ictaluri |
| Q8GNG1 | 53.98 | 2,00E-96 | galF | Alpha-D-glucosyl-1-phosphate uridylyltransferase | 2.7.7.9 | Lipopolysaccharide biosynthesis, Nucleotidyltransferase, Transferase | Escherichia coli |

**Genomic Object Editor: llo3172**

| PB id | Ident % | Eval | Gene | Description | EC number | Keywords | Organism |
|---|---|---|---|---|---|---|---|
| D3HMD6 | 100 | 0 | galE | UDP-glucose 4-epimerase | 5.1.3.2 | Carbohydrate metabolism, Isomerase, NAD, Reference proteome | Legionella longbeachae serogroup 1 (strain NSW150) |
| D3HMD0 | 63.44 | 1,00E-160 | galE | UDP-glucose 4-epimerase | 5.1.3.2 | Carbohydrate metabolism, Isomerase, NAD, Reference proteome | Legionella longbeachae serogroup 1 (strain NSW150) |
| R4YS92 | 60.42 | 6,00E-147 | galE | UDP-glucose 4-epimerase | 5.1.3.2 | Carbohydrate metabolism, Isomerase, NAD, Reference proteome | Oleispira antarctica RB-8 |
| A0A2E7E932 | 57.75 | 1,00E-131 | galE | UDP-glucose 4-epimerase | 5.1.3.2 | Carbohydrate metabolism, Isomerase, NAD | Oceanospirillaceae bacterium |
| I0JQM5 | 53.89 | 2,00E-129 | galE2 | UDP-glucose 4-epimerase | 5.1.3.2 | Carbohydrate metabolism, Isomerase, NAD, Reference proteome | Halobacillus halophilus (strain ATCC 35676 / DSM 2266 / JCM 20832 / NBRC 102448/ NCIMB 2269) |
| A0A173ZI12 | 55.45 | 4,00E-129 | galE_1 | UDP-glucose 4-epimerase | 5.1.3.2 | Carbohydrate metabolism, Isomerase, NAD, Reference proteome | Clostridium ventriculi |
| A0A1H8C287 | 55.32 | 2,00E-127 | _ | UDP-glucose 4-epimerase | 5.1.3.2 | Carbohydrate metabolism, Isomerase, NAD | Paenisporosarcina quisquiliarum |
| A0A1I2KKT0 | 53.59 | 3,00E-127 | _ | UDP-glucose 4-epimerase | 5.1.3.2 | Carbohydrate metabolism, Isomerase, NAD | Halobacillus alkaliphilus |
| Q2QD27 | 54.27 | 1,00E-120 | gne | UDP-glucose 4-epimerase | 5.1.3.2 | Carbohydrate metabolism, Isomerase, NAD | Aeromonas hydrophila |
| Q5JBH4 | 53.64 | 2,00E-116 | galE | UDP-glucose 4-epimerase | 5.1.3.2 | Carbohydrate metabolism, Isomerase, NAD | Escherichia coli |
| Q4KXC7 | 53.03 | 3,00E-115 | galE | UDP-glucose 4-epimerase | 5.1.3.2 | Carbohydrate metabolism, Isomerase, NAD | Escherichia coli |
| Q937X4 | 50.76 | 3,00E-114 | galE | UDP-glucose 4-epimerase | 5.1.3.2 | Carbohydrate metabolism, Isomerase, NAD | Edwardsiella ictaluri |

**Supplementary Table 1** List of the highest hits of the *L. longbeachae* capsule cluster genes against the NCBI database (Trembl).

| | | | | | | | |
|---|---|---|---|---|---|---|---|
| M9VRS7 | 49.25 | 6,00E-114 | galE2 | UDP-glucose 4-epimerase | 5.1.3.2 | Carbohydrate metabolism, Isomerase, NAD | Streptococcus oralis |
| A0A0F7YYT2 | 51.83 | 8,00E-114 | gne1 | UDP-glucose 4-epimerase | 5.1.3.2 | Carbohydrate metabolism, Isomerase, NAD | Acinetobacter baumannii AB5075 |
| M9P0X5 | 52.44 | 2,00E-113 | galE | UDP-glucose 4-epimerase | 5.1.3.2 | Carbohydrate metabolism, Isomerase, NAD | Providencia alcalifaciens |
| Q9X3S6 | 50.91 | 2,00E-113 | galE | UDP-glucose 4-epimerase | 5.1.3.2 | Carbohydrate metabolism, Isomerase, NAD | Neisseria meningitidis |
| O54385 | 53.05 | 3,00E-113 | galE | UDP-glucose 4-epimerase | 5.1.3.2 | Carbohydrate metabolism, Isomerase, NAD | Brucella abortus |
| Q9F8B2 | 51.66 | 4,00E-113 | galE | UDP-glucose 4-epimerase | 5.1.3.2 | Carbohydrate metabolism, Isomerase, NAD | Moraxella catarrhalis |

**Genomic Object Editor: llo3173**

| PB id | Ident % | Eval | Gene | Description | EC number | Keywords | Organism |
|---|---|---|---|---|---|---|---|
| D3HMD7 | 99.69 | 0 | fcl | GDP-L-fucose synthase | 1.1.1.271 | Isomerase, Multifunctional enzyme, NADP, Oxidoreductase, Reference proteome | Legionella longbeachae serogroup 1 (strain NSW150) |
| W8R8L8 | 69.62 | 3,00E-167 | fcl | GDP-L-fucose synthase | 1.1.1.271 | Isomerase, Multifunctional enzyme, NADP, Oxidoreductase | Pseudomonas stutzeri |
| A0A2N8SP15 | 69.3 | 3,00E-167 | fcl | GDP-L-fucose synthase | 1.1.1.271 | Isomerase, Multifunctional enzyme, NADP, Oxidoreductase | Pseudomonas stutzeri |
| A0A0D9AQH8 | 68.97 | 1,00E-166 | fcl | GDP-L-fucose synthase | 1.1.1.271 | Isomerase, Multifunctional enzyme, NADP, Oxidoreductase | Pseudomonas stutzeri |
| A0A2E2GGE9 | 69.28 | 4,00E-166 | fcl | GDP-L-fucose synthase | 1.1.1.271 | Isomerase, Multifunctional enzyme, NADP, Oxidoreductase | Pseudomonas sp |
| A0A2N1CMA6 | 68.97 | 4,00E-166 | fcl | GDP-L-fucose synthase | 1.1.1.271 | Isomerase, Multifunctional enzyme, NADP, Oxidoreductase | Pseudomonas sp. Choline-3u-10 |
| A0A172WSZ4 | 68.97 | 6,00E-166 | fcl | GDP-L-fucose synthase | 1.1.1.271 | Isomerase, Multifunctional enzyme, NADP, Oxidoreductase | Pseudomonas stutzeri |
| A0A1M5PMK2 | 68.97 | 2,00E-165 | fcl | GDP-L-fucose synthase | 1.1.1.271 | Isomerase, Multifunctional enzyme, NADP, Oxidoreductase | Pseudomonas xanthomarina DSM 18231 |
| A0A2W5D1Z8 | 69.09 | 5,00E-165 | fcl | GDP-L-fucose synthase | 1.1.1.271 | Isomerase, Multifunctional enzyme, NADP, Oxidoreductase | Pseudomonas kuykendallii |
| A0A078LSR6 | 68.44 | 7,00E-165 | fcl | GDP-L-fucose synthase | 1.1.1.271 | Isomerase, Multifunctional enzyme, NADP, Oxidoreductase, Reference proteome | Pseudomonas saudiphocaensis |
| K4NRM2 | 66.98 | 1,00E-161 | fcl | GDP-L-fucose synthase | 1.1.1.271 | Isomerase, Multifunctional enzyme, NADP, Oxidoreductase | Yersinia similis |
| Q56873 | 67.61 | 1,00E-158 | fcl | GDP-L-fucose synthase | 1.1.1.271 | Isomerase, Multifunctional enzyme, NADP, Oxidoreductase | Yersinia enterocolitica |
| Q5ND84 | 64.89 | 1,00E-152 | fcl | GDP-L-fucose synthase | 1.1.1.271 | Isomerase, Multifunctional enzyme, NADP, Oxidoreductase | Yersinia sp. A125 KOH2 |
| Q4KXD6 | 64.38 | 2,00E-151 | fcl | GDP-L-fucose synthase | 1.1.1.271 | Isomerase, Multifunctional enzyme, NADP, Oxidoreductase | Escherichia coli |
| Q5XL46 | 63.21 | 1,00E-150 | wcaG | GDP-L-fucose synthase | 1.1.1.271 | Isomerase, Multifunctional enzyme, NADP, Oxidoreductase | Klebsiella pneumoniae |
| A5Y7V9 | 63.44 | 1,00E-149 | fcl | GDP-L-fucose synthase | 1.1.1.271 | Isomerase, Multifunctional enzyme, NADP, Oxidoreductase, Reference proteome | Salmonella enterica subsp. enterica serovar Poona |
| Q9F7A3 | 62.81 | 8,00E-147 | fcl | GDP-L-fucose synthase | 1.1.1.271 | Isomerase, Multifunctional enzyme, NADP, Oxidoreductase | Salmonella typhimurium |

**Genomic Object Editor: llo3174**

| PB id | Ident % | Eval | Gene | Description | EC number | Keywords | Organism |
|---|---|---|---|---|---|---|---|
| D3HMD8 | 100 | 0 | _ | Putative glycosyltransferase | _ | Reference proteome, Transferase | Legionella longbeachae serogroup 1 (strain NSW150) |
| A0A011PZP5 | 50.53 | 1,00E-98 | kfoC_1 | Chondroitin polymerase | _ | Transferase | Candidatus Accumulibacter sp. BA-92 |
| A0A011PC78 | 53.31 | 9,00E-81 | kfoC_1 | Chondroitin polymerase | _ | Reference proteome, Transferase | Candidatus Accumulibacter sp. SK-11 |
| A0A011PMS9 | 51.82 | 3,00E-78 | kfoC_1 | Chondroitin polymerase | _ | Reference proteome, Transferase | Candidatus Accumulibacter sp. SK-12 |
| A4BRS0 | 48.95 | 5,00E-71 | _ | Glycosyltransferase | _ | Reference proteome, Transferase | Nitrococcus mobilis Nb-231 |
| A4CTD8 | 45.56 | 8,00E-67 | _ | Glycosyltransferase | _ | Transferase | Synechococcus sp. (strain WH7805) |
| A0A1R4GZF7 | 54.37 | 1,00E-66 | _ | Glyco_trans_2-like domain-containing protein | _ | Reference proteome | Crenothrix polyspora |
| L8BAN0 | 45.45 | 2,00E-66 | _ | Putative Glycosyltransferase | _ | Transferase | Rubrivivax gelatinosus S1 |
| D9PNG8 | 46.79 | 1,00E-61 | _ | Glycosyltransferase | _ | Transferase | sediment metagenome |
| A0A2V6Q4G8 | 45.12 | 8,00E-60 | _ | Glycosyltransferase | _ | Transferase | Candidatus Rokubacteria bacterium |
| Q6XQ52 | 29.11 | 2,00E-16 | wbsK | Glycosyl transferase family 2 | _ | Transferase | Escherichia coli |
| D0QYM5 | 30.39 | 4,00E-16 | acbD | AcbD | _ | _ | Avibacterium paragallinarum |
| D0QYL9 | 30.39 | 4,00E-16 | acbD | AcbD | _ | _ | Avibacterium paragallinarum |
| Q9RFX0 | 36.43 | 5,00E-16 | cps7H | Putative glycosyltransferase Cps7H | _ | Transferase | Streptococcus suis |
| K4P2X6 | 29.74 | 0.000000000000002 | wbyL | WbyL | _ | _ | Yersinia similis |
| F8RC13 | 34.13 | 0.000000000000005 | wpaD | WpaD | _ | _ | Providencia alcalifaciens |
| Q8GMK1 | 32.17 | 0.00000000000001 | wbsA | Glycosyl transferase family 2 | _ | Transferase | Escherichia coli |
| Q56869 | 28.95 | 0.000000000001 | wbcG | WbcG | _ | _ | Yersinia enterocolitica |

**Supplementary Table 1** List of the highest hits of the *L. longbeachae* capsule cluster genes against the NCBI database (Trembl).

| | | | | | | | |
|---|---|---|---|---|---|---|---|
| A3F4D9 | 31.48 | 0.0000000000004 | epsM | EpsM | _ | Membrane, Plasmid, Transmembrane, Transmembrane helix | Lactococcus lactis subsp. cremoris |
| F1CLM1 | 33.06 | 0.0000000000008 | wbzE | Putative glycosyl transferase | _ | Transferase | Yersinia pseudotuberculosis |

**Genomic Object Editor: llo3175**

| PB id | Ident % | Eval | Gene | Description | EC number | Keywords | Organism |
|---|---|---|---|---|---|---|---|
| D3HMD9 | 100 | 0 | _ | Putative polysaccharide biosynthesis dehydrogenase/reductase protein | _ | Membrane, Reference proteome, Transmembrane, Transmembrane helix | Legionella longbeachae serogroup 1 (strain NSW150) |
| A0A1G0WYL4 | 53.36 | 8,00E-88 | _ | Short-chain dehydrogenase | _ | _ | Legionellales bacterium RIFCSPHIGHO2_12_FULL_37_14 |
| A0A1G0GZU1 | 53.91 | 5,00E-86 | _ | Short-chain dehydrogenase | _ | Membrane, Transmembrane, Transmembrane helix | Gammaproteobacteria bacterium RIFCSPHIGHO2_12_FULL_40_19 |
| A0A1G0G9X5 | 50.4 | 7,00E-77 | _ | Uncharacterized protein | _ | Membrane, Transmembrane, Transmembrane helix | Gammaproteobacteria bacterium RIFCSPHIGHO2_02_FULL_39_13 |
| K2BBG0 | 50.4 | 7,00E-77 | _ | Polysaccharide biosynthesis dehydrogenase/reductase protein | _ | Membrane, Transmembrane, Transmembrane helix | uncultured bacterium |
| A0A2G6HQW6 | 48.21 | 1,00E-74 | _ | Short-chain dehydrogenase | _ | Membrane, Transmembrane, Transmembrane helix | Thiothrix nivea |
| A0A2G6DK36 | 47.81 | 4,00E-72 | _ | Short-chain dehydrogenase | _ | Membrane, Transmembrane, Transmembrane helix | Proteobacteria bacterium |
| A0A228K644 | 46.48 | 3,00E-71 | _ | SDR family oxidoreductase | _ | _ | Burkholderia sp. AU27893 |
| A0A2S5DYT0 | 46.48 | 3,00E-71 | _ | KR domain-containing protein | _ | _ | Burkholderia contaminans |
| A0A103ZCF2 | 44.92 | 2,00E-70 | _ | Capsular biosynthesis protein | _ | _ | Burkholderia cepacia |
| A0A1V6KPJ8 | 45.31 | 3,00E-70 | _ | Capsular biosynthesis protein | _ | _ | Burkholderia cenocepacia |
| Q9KHD1 | 31.87 | 2,00E-19 | _ | Putative beta-ketoacyl reductase | _ | _ | Streptomyces griseus subsp. griseus |
| C6ZD46 | 30.89 | 9,00E-19 | ydfG | NADP-dependent L-serine/L-allo-threonine dehydrogenase ydfG | 1.1.1.- | Oxidoreductase | Legionella jamestowniensis |
| Q5EGQ6 | 32.02 | 5,00E-18 | rkpH | RkpH | 1.1.1.56 | Oxidoreductase | Rhizobium fredii |
| B8XIJ9 | 28.72 | 9,00E-17 | _ | Ketoacyl-reductase like protein | _ | _ | Karlodinium veneficum |
| K7ZSN0 | 29.79 | 1,00E-16 | lgnI | Gluconate 5-dehydrogenase | _ | Reference proteome | Paracoccus laeviglucosivorans |
| Q8RR58 | 25.53 | 3,00E-16 | acrM | Acyl coenzyme A reductase | _ | Coiled coil | Acinetobacter sp. M-1 |
| Q7LZT0 | 27.92 | 6,00E-16 | _ | 3(or 17)beta-hydroxysteroid dehydrogenase I | 1.1.1.51 | Oxidoreductase | Anguilla japonica |
| Q9API9 | 29.69 | 6,00E-16 | phaB | Acetoacetyl-CoA reductase | 1.1.1.36 | Oxidoreductase | Methylorubrum extorquens |
| Q6RH38 | 27.92 | 0.000000000000001 | _ | 17b-hydroxysteroid dehydrogenase type I | _ | Oxidoreductase | Anguilla japonica |

**Genomic Object Editor: llo3176**

| PB id | Ident % | Eval | Gene | Description | EC number | Keywords | Organism |
|---|---|---|---|---|---|---|---|
| D3HME0 | 100 | 0 | _ | Putative glycosyltransferase | 2.4.1.- | Glycosyltransferase, Reference proteome, Transferase | Legionella longbeachae serogroup 1 (strain NSW150) |
| A0A2S5SX42 | 46.4 | 1,00E-100 | _ | Glycosyltransferase family 1 protein | _ | Reference proteome, Transferase | Zhizhongheella caldifontis |
| H8FUS8 | 43.8 | 2,00E-94 | _ | Putative glycosyltransferase | 2.4.-.- | Glycosyltransferase, Reference proteome, Transferase | Phaeospirillum molischianum DSM 120 |
| A0A259BE26 | 43.86 | 1,00E-93 | _ | Glycos_transf_1 domain-containing protein | _ | _ | Halothiobacillus sp. 24-54-40 |
| A0A0W0Z520 | 41.88 | 1,00E-91 | _ | Glycosyltransferase | 2.4.1.- | Glycosyltransferase, Reference proteome, Transferase | Legionella shakespearei DSM 23087 |
| A0A2N3B0J3 | 45.1 | 1,00E-85 | _ | Glycosyltransferase family 1 protein | _ | Transferase | Alphaproteobacteria bacterium HGW-Alphaproteobacteria-7 |
| A0A255Y7H4 | 38.96 | 8,00E-84 | _ | Glycos_transf_1 domain-containing protein | _ | Reference proteome | Sandarakinorhabdus cyanobacteriorum |
| A0A1X1PB15 | 43.84 | 9,00E-81 | _ | Glycos_transf_1 domain-containing protein | _ | Coiled coil | Burkholderia puraquae |
| T2N217 | 40.21 | 1,00E-80 | _ | Glycos_transf_1 domain-containing protein | _ | _ | Ralstonia sp. 5_2_56FAA |
| M4QN28 | 35.29 | 2,00E-64 | wbdA | Mannosyltransferase | _ | Glycosyltransferase, Transferase | Escherichia coli |
| Q47593 | 35.04 | 7,00E-64 | mtfA | Mannosyltransferase A | _ | Glycosyltransferase, Transferase | Escherichia coli |
| Q9LC66 | 35.04 | 7,00E-64 | wbdA | Mannosyltransferase | _ | Glycosyltransferase, Transferase | Klebsiella pneumoniae |
| Q9LC67 | 35.58 | 1,00E-62 | wbdA | Mannosyltransferase | _ | Glycosyltransferase, Transferase | Escherichia coli |
| C8YZ32 | 35.34 | 1,00E-60 | wejI | WejI | _ | _ | Escherichia coli |
| O84908 | 32.57 | 1,00E-33 | wbpX | Glycosyltransferase Gtf1 | _ | Transferase | Pseudomonas aeruginosa |
| Q93UK1 | 26.73 | 0.0000000000001 | wcbB | WcbB | _ | _ | Burkholderia pseudomallei |
| M4M6T9 | 29.17 | 0.0000000006 | _ | Glycosyl transferase group 1 | _ | Transferase | Acidiphilium sp. PM |
| O84909 | 31.76 | 0.0000002 | wbpY | Glycosyltransferase WbpY | _ | Transferase | Pseudomonas aeruginosa |
| Q93UJ8 | 35.65 | 0.000005 | wcbE | WcbE | _ | _ | Burkholderia pseudomallei |
| Q9RMT9 | 31.97 | 0.000005 | wbdB | WbdB | _ | _ | Klebsiella pneumoniae |

**Genomic Object Editor: llo3177**

| PB id | Ident % | Eval | Gene | Description | EC number | Keywords | Organism |
|---|---|---|---|---|---|---|---|
| D3HME3 | 100 | 0 | _ | Uncharacterized protein | _ | Membrane, Reference proteome, Transmembrane, Transmembrane helix | Legionella longbeachae serogroup 1 (strain NSW150) |
| A0A1G0GZV7 | 49.04 | 5,00E-130 | _ | Uncharacterized protein | _ | _ | Gammaproteobacteria bacterium RIFCSPHIGHO2_12_FULL_40_19 |
| A0A1T4X2W9 | 36.1 | 5,00E-73 | _ | Glycosyltransferase family 28 C-terminal domain-containing protein | _ | Coiled coil, Reference proteome, Transferase | Clostridium sp. USBA 49 |
| U2CYI4 | 34 | 2,00E-68 | _ | Glycosyltransferase family 28 protein | _ | Coiled coil, Reference proteome, Transferase | Clostridiales bacterium oral taxon 876 str. F0540 |
| A0A1T4Y1N0 | 32.46 | 2,00E-60 | _ | Glycosyltransferase family 28 C-terminal domain-containing protein | _ | Coiled coil, Reference proteome, Transferase | Caloramator quimbayensis |
| A0A2C6TA21 | 30.79 | 2,00E-51 | _ | Uncharacterized protein | _ | Coiled coil | Nostoc linckia z16 |
| A0A0C2V1S8 | 30.18 | 3,00E-49 | _ | Glyco_tran_28_C domain-containing protein | _ | Reference proteome | Paenibacillus sp. VKM B-2647 |
| A0A1I1D9I8 | 22.14 | 0.000000004 | _ | UDP-N-acetylglucosamine:LPS N-acetylglucosamine transferase | _ | Glycosyltransferase, Reference proteome, Transferase | Brevinema andersonii |

| | | | | | | | |
|---|---|---|---|---|---|---|---|
| A0A1J4VM15 | 21.36 | 0.00000005 | _ | Uncharacterized protein | _ | Glycosyltransferase, Membrane, Transferase, Transmembrane, Transmembrane helix | Candidatus Omnitrophica bacterium CG1_02_46_14 |
| A0A1V4IVN9 | 20.83 | 0.0000001 | ugtP | Processive diacylglycerol beta-glucosyltransferase | 2.4.1.- | Glycosyltransferase, Reference proteome, Transferase | Clostridium chromiireducens |

**Genomic Object Editor: llo3178**

| PB id | Ident % | Eval | Gene | Description | EC number | Keywords | Organism |
|---|---|---|---|---|---|---|---|
| D3HME4 | 100 | 0 | _ | Putative aminotransferase | 2.3.1.47 | Acyltransferase, Aminotransferase, Pyridoxal phosphate | Legionella longbeachae serogroup 1 (strain NSW150) |
| A0A1G0WY90 | 63.45 | 0 | _ | 8-amino-7-oxononanoate synthase | _ | Pyridoxal phosphate | Legionellales bacterium RIFCSPHIGHO2_12_FULL_37_14 |
| A0A157QUK5 | 61.34 | 0 | wcbT | Polyketide synthase | 2.3.1.- | Acyltransferase, Transferase | Bordetella trematum |
| A0A157SWX8 | 58.8 | 2,00E-180 | wcbT | Polyketide synthase | 2.3.1.- | Acyltransferase, Pyridoxal phosphate, Transferase | Bordetella ansorpii |
| A0A132F5H0 | 60.84 | 2,00E-179 | _ | 8-amino-7-oxononanoate synthase | _ | Pyridoxal phosphate | Burkholderia pseudomultivorans |
| A0A1I3G3G2 | 59.77 | 2,00E-179 | _ | 8-amino-7-oxononanoate synthase | _ | Pyridoxal phosphate | Collimonas sp. OK307 |
| A0A1B4F580 | 61.07 | 3,00E-179 | _ | 8-amino-7-oxononanoate synthase | _ | Pyridoxal phosphate | Burkholderia sp. LA-2-3-30-S1-D2 |
| A0A088UAL4 | 60.84 | 4,00E-179 | _ | Beta-eliminating lyase family protein | _ | Lyase, Pyridoxal phosphate | Burkholderia cenocepacia |
| A0A2A4CCM4 | 60.61 | 1,00E-178 | _ | 8-amino-7-oxononanoate synthase | _ | Pyridoxal phosphate | Burkholderia sp. IDO3 |
| Q5EGQ7 | 48.95 | 3,00E-134 | rkpG | RkpG | 2.3.1.29 | Acyltransferase, Transferase | Rhizobium fredii |
| Q52936 | 51.62 | 3,00E-123 | rkpG | Acyl-transferase | _ | Transferase | Rhizobium meliloti |
| A7BFV7 | 33.33 | 6,00E-66 | spt | Serine palmitoyltransferase | 2.3.1.50 | Acyltransferase, Pyridoxal phosphate, Transferase | Sphingobacterium spiritivorum |
| Q9AJN1 | 34.38 | 1,00E-65 | bioF | 8-amino-7-ketopelargonate synthase | 2.3.1.47 | Biotin biosynthesis, Pyridoxal phosphate, Transferase | Kurthia sp. 538-KA26 |
| Q9AJM7 | 34.82 | 2,00E-65 | bioFII | 8-amino-7-ketopelargonate synthase | 2.3.1.47 | Biotin biosynthesis, Pyridoxal phosphate, Transferase | Kurthia sp. 538-KA26 |
| A7BFV6 | 35.39 | 9,00E-65 | spt | 8-amino-7-oxononanoate synthase | 2.3.1.47, 2.3.1.50 | Acyltransferase, Pyridoxal phosphate, Transferase | Sphingobacterium multivorum |
| A7BFV8 | 32.14 | 2,00E-57 | spt | Serine palmitoyltransferase | 2.3.1.50 | Acyltransferase, Pyridoxal phosphate, Transferase | Bacteriovorax stolpii |
| B2XR73 | 32.03 | 2,00E-56 | LCB2 | Serine C-palmitoyltransferase | 2.3.1.50 | Endoplasmic reticulum, Lipid metabolism, Membrane, Pyridoxal phosphate, Sphingolipid metabolism, Transferase, Transmembrane | Nicotiana benthamiana |

**Genomic Object Editor: llo3179**

| PB id | Ident % | Eval | Gene | Description | EC number | Keywords | Organism |
|---|---|---|---|---|---|---|---|
| D3HME5 | 100 | 0 | rkpA | Malonyl CoA-acyl carrier protein transacylase | _ | Multifunctional enzyme, NADP, Phosphopantetheine, Phosphoprotein, Reference proteome, Transferase | Legionella longbeachae serogroup 1 (strain NSW150) |
| A0A1G0H3Y5 | 48.92 | 0 | _ | Malonyl CoA-acyl carrier protein transacylase | _ | Multifunctional enzyme, NADP, Phosphopantetheine, Phosphoprotein, Transferase | Gammaproteobacteria bacterium RIFCSPHIGHO2_12_FULL_38_11 |
| A0A1G0GZW1 | 49.27 | 0 | _ | Malonyl CoA-acyl carrier protein transacylase | _ | Multifunctional enzyme, NADP, Phosphopantetheine, Phosphoprotein, Transferase | Gammaproteobacteria bacterium RIFCSPHIGHO2_12_FULL_40_19 |
| A0A1G0G9V3 | 47.94 | 0 | _ | Malonyl CoA-acyl carrier protein transacylase | _ | Multifunctional enzyme, NADP, Phosphopantetheine, Phosphoprotein, Transferase | Gammaproteobacteria bacterium RIFCSPHIGHO2_02_FULL_39_13 |
| K2C9B0 | 47.94 | 0 | _ | Carrier domain-containing protein | _ | Multifunctional enzyme, NADP, Phosphopantetheine, Phosphoprotein, Transferase | uncultured bacterium |
| A0A1G0WYA8 | 46.27 | 0 | _ | Malonyl CoA-acyl carrier protein transacylase | _ | Coiled coil, Multifunctional enzyme, NADP, Phosphopantetheine, Phosphoprotein, Transferase | Legionellales bacterium RIFCSPHIGHO2_12_FULL_37_14 |
| A0A1T4X2T5 | 41.79 | 0 | _ | Malonyl CoA-acyl carrier protein transacylase | _ | Coiled coil, Multifunctional enzyme, NADP, Phosphopantetheine, Phosphoprotein, Reference proteome, Transferase | Thiothrix eikelboomii |
| A0A2G6HQX1 | 41.42 | 0 | _ | Malonyl CoA-acyl carrier protein transacylase | _ | Multifunctional enzyme, NADP, Phosphopantetheine, Phosphoprotein, Transferase | Thiothrix nivea |
| A0A2G6DKI5 | 41.28 | 0 | _ | Malonyl CoA-acyl carrier protein transacylase | _ | Multifunctional enzyme, NADP, Phosphopantetheine, Phosphoprotein, Transferase | Proteobacteria bacterium |
| A0A1H9YTQ4 | 40.46 | 0 | _ | Malonyl CoA-acyl carrier protein transacylase | _ | Multifunctional enzyme, NADP, Phosphopantetheine, Phosphoprotein, Transferase | Nitrosomonas europaea |
| Q82UT4 | 40.46 | 0 | rkpA | Malonyl CoA-acyl carrier protein transacylase | _ | Multifunctional enzyme, NADP, Phosphopantetheine, Phosphoprotein, Reference proteome, Transferase | Nitrosomonas europaea (strain ATCC 19718 / CIP 103999 / KCTC 2705 / NBRC 14298) |

| | | | | | | | |
|---|---|---|---|---|---|---|---|
| A0A238XMI4 | 40.34 | 0 | _ | Malonyl CoA-acyl carrier protein transacylase | _ | Multifunctional enzyme, NADP, Phosphopantetheine, Phosphoprotein, Transferase | Methylobacillus rhizosphaerae |
| Q6E7K0 | 30.12 | 0 | _ | 3-hydroxyacyl-CoA dehydrogenase | 1.1.1.35 | 3D-structure, Coiled coil, Multifunctional enzyme, NADP, Nucleotide-binding, Phosphopantetheine, Phosphoprotein, Transferase | Lyngbya majuscula |
| A0A0A0WDX2 | 31.31 | 0 | puwB | PuwB | _ | Coiled coil, Multifunctional enzyme, NADP, Phosphopantetheine, Phosphoprotein, Transferase | Cylindrospermum alatosporum CCALA 988 |
| F4Y426 | 28.41 | 0 | _ | CurJ | _ | 3D-structure, Phosphopantetheine, Phosphoprotein, Reference proteome, Transferase | Moorea producens 3L |
| Q6DNE3 | 28.41 | 0 | curJ | CurJ | _ | 3D-structure, Phosphopantetheine, Phosphoprotein, Transferase | Lyngbya majuscula |
| Q5EGQ8 | 31.94 | 0 | rkpA | Malonyl CoA-acyl carrier protein transacylase | _ | Multifunctional enzyme, Phosphopantetheine, Phosphoprotein, Transferase | Rhizobium fredii |
| Q6ZY03 | 44.68 | 0 | pks4 | Polyketide synthase I | _ | Phosphopantetheine, Phosphoprotein, Transferase | uncultured bacterium |
| A0A1B3TNB2 | 35.66 | 0 | hapB | Malonyl CoA-acyl carrier protein transacylase | _ | Coiled coil, Multifunctional enzyme, NADP, Phosphopantetheine, Phosphoprotein, Transferase | Byssovorax cruenta |
| Q9KIZ7 | 34.07 | 0 | epoD | Malonyl CoA-acyl carrier protein transacylase | _ | Coiled coil, Multifunctional enzyme, Phosphopantetheine, Phosphoprotein, Transferase | Sorangium cellulosum |
| Q3KRU5 | 25.72 | 0 | pksX1 | PKSX1 | _ | Acyltransferase, Coiled coil, Methyltransferase, Multifunctional enzyme, NADP, Oxidoreductase, Phosphopantetheine, Phosphoprotein, Transferase | Xylaria sp. BCC 1067 |
| Q8RJY0 | 34.6 | 0 | stiG | Malonyl CoA-acyl carrier protein transacylase | _ | Coiled coil, Phosphopantetheine, Phosphoprotein, Transferase | Stigmatella aurantiaca |
| Q93TW8 | 34.15 | 0 | mxaD | Malonyl CoA-acyl carrier protein transacylase | _ | Coiled coil, Phosphopantetheine, Phosphoprotein, Transferase | Stigmatella aurantiaca |

**Genomic Object Editor: llo3180**

| PB id | Ident % | Eval | Gene | Description | EC number | Keywords | Organism |
|---|---|---|---|---|---|---|---|
| D3HME6 | 100 | 0 | capI | Protein capI | _ | Reference proteome | Legionella longbeachae serogroup 1 (strain NSW150) |
| D5BMZ1 | 63.17 | 3,00E-152 | _ | Putative nucleotide sugar epimerase | 5.1.3.- | Isomerase, Reference proteome | Puniceispirillum marinum (strain IMCC1322) |
| A0A149VYE0 | 65.06 | 4,00E-152 | rfbB_2 | dTDP-glucose 4,6-dehydratase | 4.2.1.46 | Lyase, Reference proteome | Ferrovum sp. Z-31 |
| A0A1Q8YJ37 | 63.64 | 2,00E-151 | _ | NAD-dependent epimerase/dehydratase | _ | Reference proteome | Rhodoferax antarcticus ANT.BR |
| A0A1F4K1Q8 | 61.76 | 4,00E-150 | _ | Protein CapI | _ | _ | Burkholderiales bacterium RIFCSPLOWO2_12_FULL_61_40 |
| A0A2E1QD17 | 60.66 | 4,00E-148 | _ | Epimerase domain-containing protein | _ | _ | Euryarchaeota archaeon |
| A0A1F9MKL1 | 62.54 | 2,00E-146 | _ | Capsular biosynthesis protein CpsI | _ | _ | Deltaproteobacteria bacterium RIFOXYD12_FULL_53_23 |
| A0A1H7HPY1 | 60.36 | 2,00E-146 | _ | UDP-glucuronate 4-epimerase | _ | _ | Roseateles sp. YR242 |
| A0A1F9KZI9 | 62.84 | 1,00E-145 | _ | Epimerase domain-containing protein | _ | _ | Deltaproteobacteria bacterium RIFOXYD12_FULL_56_24 |
| Q6U8B8 | 59.16 | 1,00E-138 | _ | Putative nucleotide sugar epimerase | _ | _ | Raoultella terrigena |
| Q9RP53 | 57.66 | 6,00E-134 | wbnF | NAD-dependent epimerase | 4.2.1.46 | Lyase | Escherichia coli |
| Q4GY28 | 55.52 | 5,00E-127 | wbnF | UDP-sugar epimerase | _ | _ | Erwinia amylovora |
| Q6URR1 | 54.63 | 9,00E-127 | nse | Putative epimerase | _ | _ | Xenorhabdus nematophila |
| O68979 | 55.22 | 3,00E-121 | wcvA | Nucleotide sugar epimerase | _ | _ | Vibrio vulnificus |
| Q56626 | 53.45 | 1,00E-116 | _ | Nucleotide sugar epimerase | _ | _ | Vibrio cholerae O139 |
| P96481 | 48.18 | 5,00E-100 | cap1J | Putative epimerase | _ | _ | Streptococcus pneumoniae |
| Q70PA0 | 44.11 | 3,00E-90 | _ | Epimerase domain-containing protein | _ | _ | Melittangium lichenicola |
| I1VCA9 | 45.24 | 6,00E-90 | GAE1 | UDP-D-glucuronate 4-epimerase 1 | _ | _ | Arabidopsis thaliana |
| Q6K9M5 | 44.97 | 1,00E-88 | _ | Os02g0791500 protein | _ | Membrane, Reference proteome, Transmembrane, Transmembrane helix | Oryza sativa subsp. japonica |
