## Supplemental Table S2 for "The unique *Legionella longbeachae* capsule favors intracellular replication and immune evasion"

**Supplementary Table 2**

CPS gene identitiy among *L. longbeachae* strains compared to NSW150.

| **Percent identity compared to NSW150** | | | | | | | | | | | | | | | | | | |
|---|---|---|---|---|---|---|---|---|---|---|---|---|---|---|---|---|---|---|
| **Strain** | **serogroup** | ***llo3148*** | ***llo3149*** | ***llo3150*** | ***llo3151*** | ***llo3152*** | ***llo3153*** | ***llo3154*** | ***llo3155*** | ***llo3156*** | ***llo3157*** | ***llo3158*** | ***llo3159*** | ***llo3160*** | ***llo3161*** | ***llo3162*** | ***llo3163*** | ***llo3164*** |
| *L. longbeachae* F1157CHC | sg1 | 98.306 | 97.658 | 98.003 | 91.154 | 97.442 | 99.365 | 99.768 | 99.743 | 99.896 | 99.608 | 96.319 | 99.940 | 99.855 | 99.054 | 98.238 | 99.786 | 97.002 |
| *L. longbeachae* 13.8300 | sg2 | 98.238 | 97.387 | 98.157 | 91.282 | 96.617 | 98.571 | 99.227 | 99.829 | 99.637 | 98.529 | -- | 100.000 | 100.000 | 100.000 | 100.000 | 100.000 | 97.002 |
| *L. longbeachae* B3526CHC | sg1 | 98.238 | 97.387 | 98.157 | 91.282 | 96.617 | 98.571 | 99.149 | 99.829 | 98.445 | 99.477 | 96.185 | 99.819 | 99.565 | 98.318 | 97.504 | 99.359 | 97.002 |
| *L. longbeachae* 13.8301 | sg2 | 98.238 | 97.387 | 98.157 | 91.282 | 96.617 | 98.571 | 99.227 | 99.829 | 99.637 | 98.529 | -- | 100.000 | 100.000 | 100.000 | 100.000 | 100.000 | 97.002 |
| *L. longbeachae* 13.8297 | sg2 | 98.238 | 97.387 | 98.157 | 91.282 | 96.617 | 98.571 | 99.227 | 99.829 | 99.637 | 98.529 | 96.319 | 100.000 | 100.000 | 100.000 | 100.000 | 100.000 | 97.002 |
| *L. longbeachae* B1445CHC | sg1 | 98.306 | 97.658 | 98.003 | 91.154 | 97.442 | 99.286 | 99.845 | 100.000 | 99.948 | 98.529 | 96.319 | 100.000 | 100.000 | 99.895 | 100.000 | 100.000 | 97.002 |
| *L. longbeachae* D-4968 | sg1 | 98.306 | 97.658 | 98.003 | 91.154 | 97.442 | 99.286 | 99.845 | 100.000 | 99.948 | 99.869 | 96.319 | 100.000 | 100.000 | 99.895 | 100.000 | 100.000 | 97.002 |
| *L. longbeachae* NSW150 | sg1 | 100.000 | 100.000 | 100.000 | 100.000 | 100.000 | 100.000 | 100.000 | 100.000 | 100.000 | 100.000 | 100.000 | 100.000 | 100.000 | 100.000 | 100.000 | 100.000 | 100.000 |
| *L. longbeachae* B41211CHC | sg1 | 100.000 | 100.000 | 100.000 | 100.000 | 100.000 | 100.000 | 99.923 | 100.000 | 100.000 | 100.000 | 100.000 | 100.000 | 100.000 | 100.000 | 100.000 | 100.000 | 100.000 |
| *L. longbeachae* NCTC11477 | sg1 | 98.374 | 97.477 | 98.003 | 91.282 | 97.442 | 99.524 | 99.923 | 100.000 | 99.948 | -- | -- | 100.000 | 100.000 | 99.895 | 100.000 | 100.000 | 97.002 |
| *L. longbeachae* FDAARGOS 201 | sg1 | 98.374 | 97.477 | 98.003 | 91.282 | 97.442 | 99.524 | 99.923 | 100.000 | 99.948 | 98.529 | 96.319 | 100.000 | 100.000 | 99.895 | 100.000 | 100.000 | 97.002 |
| *L. longbeachae* FDAARGOS 1481 | sg1 | 98.374 | 97.477 | 98.003 | 91.282 | 97.442 | 99.524 | 99.923 | 100.000 | 99.948 | 98.529 | 96.319 | 100.000 | 100.000 | 99.895 | 100.000 | 100.000 | 97.002 |

| **Percent identity compared to NSW150** | | | | | | | | | | | | | | | | | |
|---|---|---|---|---|---|---|---|---|---|---|---|---|---|---|---|---|---|
| **Strain** | **serogroup** | ***llo3165*** | ***llo3166*** | ***llo3167*** | ***llo3168*** | ***llo3169*** | ***llo3170*** | ***llo3171*** | ***llo3172*** | ***llo3173*** | ***llo3174*** | ***llo3175*** | ***llo3176*** | ***llo3177*** | ***llo3178*** | ***llo3179*** | ***llo3180*** |
| *L. longbeachae* F1157CHC | sg1 | -- | -- | 85.659 | -- | -- | 99.880 | 99.336 | 100.000 | 100.000 | 100.000 | 99.872 | 100.000 | 100.000 | 100.000 | 100.000 | 100.000 |
| *L. longbeachae* 13.8300 | sg2 | -- | -- | 85.756 | -- | -- | 99.640 | 99.225 | 99.604 | 99.689 | 99.431 | 99.489 | 98.115 | 98.424 | 98.712 | 99.224 | 99.705 |
| *L. longbeachae* B3526CHC | sg1 | -- | -- | 85.756 | -- | -- | 99.520 | 99.225 | 99.604 | 99.689 | 99.431 | 99.489 | 98.115 | 98.424 | 98.712 | 99.224 | 99.705 |
| *L. longbeachae* 13.8301 | sg2 | -- | -- | 85.756 | -- | -- | 99.640 | 99.225 | 99.604 | 99.689 | 99.431 | 99.489 | 98.115 | 98.424 | 98.712 | 99.224 | 99.705 |
| *L. longbeachae* 13.8297 | sg2 | -- | -- | 85.756 | -- | -- | 99.640 | 99.225 | 99.604 | 99.689 | 99.431 | 99.489 | 98.115 | 98.424 | 98.712 | 99.224 | 99.705 |
| *L. longbeachae* B1445CHC | sg1 | -- | -- | 85.756 | -- | -- | 99.880 | 99.779 | 99.901 | 100.000 | 100.000 | 100.000 | 100.000 | 100.000 | 99.924 | 100.000 | 100.000 |
| *L. longbeachae* D-4968 | sg1 | -- | -- | 85.756 | -- | -- | 99.880 | 99.779 | 99.901 | 100.000 | 100.000 | 100.000 | 100.000 | 100.000 | 100.000 | 100.000 | 100.000 |
| *L. longbeachae* NSW150 | sg1 | 100.000 | 100.000 | 100.000 | 100.000 | 100.000 | 100.000 | 100.000 | 100.000 | 100.000 | 100.000 | 100.000 | 100.000 | 100.000 | 100.000 | 100.000 | 100.000 |
| *L. longbeachae* B41211CHC | sg1 | 100.000 | 100.000 | 100.000 | 100.000 | 100.000 | 100.000 | 100.000 | 100.000 | 100.000 | 100.000 | 100.000 | 100.000 | 100.000 | 100.000 | 100.000 | 100.000 |
| *L. longbeachae* NCTC11477 | sg1 | -- | -- | 85.756 | -- | -- | 99.880 | 99.779 | 99.901 | 100.000 | 100.000 | 100.000 | 100.000 | 100.000 | 100.000 | 100.000 | 100.000 |
| *L. longbeachae* FDAARGOS 201 | sg1 | -- | -- | 85.756 | -- | -- | 99.880 | 99.779 | 99.901 | 100.000 | 100.000 | 100.000 | 100.000 | 100.000 | 100.000 | 100.000 | 100.000 |
| *L. longbeachae* FDAARGOS 1481 | sg1 | -- | -- | 85.756 | -- | -- | 99.880 | 99.779 | 99.901 | 100.000 | 100.000 | 100.000 | 100.000 | 100.000 | 100.000 | 100.000 | 100.000 |
